# When Deeper Analysis Weakens Aesthetic Experience: Behavioral and Brain Network Evidence

**DOI:** 10.64898/2026.08.06.743328

**Authors:** Lei Ha, Chuyang Sun, Rixin Tang

## Abstract

Analysis does not always enhance aesthetic experience. Philosophical accounts have long suggested that decomposing an aesthetic experience into determinate components may weaken it, yet this possibility has rarely been tested experimentally. To examine whether, when, and how analysis produces divergent effects on aesthetic experience, we conducted two experiments manipulating analysis depth. Experiment 1 showed that, during affective analysis of visual art, deep analysis produced a significantly weaker increase in aesthetic ratings than shallow analysis. In Experiment 2, we selected this condition to investigate the underlying mechanism. The behavioral effect was replicated: deep analysis removed the increase produced by shallow analysis without reducing ratings below the image baseline. Frequency-resolved brain network analysis further revealed a stronger task-related component and higher spatial entropy within the default mode network under deep analysis. Network-behavior correlations observed under shallow analysis were absent under deep analysis, suggesting reduced correspondence between the default-mode network (DMN) organization and aesthetic experience. Exploratory analyses further showed that spatial weights in the lateral temporal cortex and inferior parietal lobule were associated with smaller increases in aesthetic ratings. Together, these findings indicate that deeper analysis can selectively weaken improvements in aesthetic experience by altering how affective information is organized within the DMN.

## 1. Introduction

“Sweet is the lore which nature brings; / Our meddling intellect / Mishapes the beauteous forms of things; / —We murder to dissect” (Wordsworth, 1800, lines 25–28). In “The Tables Turned,” Wordsworth voiced a longstanding concern: when analysis becomes too specific, aesthetic experience may be weakened. The concern does not imply that analysis is harmful in itself; it suggests instead that the effect of analysis depends on how it is carried out. Imagine following a guide through a gallery. A broad remark about a painting may heighten your sense of its beauty, whereas a more detailed, itemized commentary may leave the experience flatter than before. The present study asks what distinguishes these two cases. Answering it bears on how aesthetic experience is communicated and on how such communication might be improved. More fundamentally, examining how and why aesthetic experience is disrupted may reveal what it requires in order to occur at all.

This phenomenon is not readily explained by existing experimental findings, which have primarily reported positive effects of analysis on aesthetic experience. Titles and background information enhance people’s understanding and evaluation of artworks (Millis, 2001; Russell, 2003; Leder et al., 2006). Guiding viewers to consider the emotions and intentions of depicted figures increases aesthetic appreciation of visual art (Iosifyan, 2021). Guided-viewing and slow-looking tasks direct attention toward specific features and encourage more careful observation, thereby enhancing understanding, engagement, and appreciation (Tishman, 2017; Estrada Gonzalez et al., 2025). Collectively, these studies indicate that analysis can facilitate aesthetic experience. However, they have compared the presence versus absence of analysis, or contrasted different forms of viewing guidance, without varying the depth of the analysis itself. The possibility that analysis might, past some point, cease to help has therefore not been tested directly.

Classical aesthetic theory suggests why such a point might exist. Kant located aesthetic judgment in the free harmony of imagination and understanding, unconstrained by determinate concepts (Kant, 1790/2000); subsuming an experience under a fixed set of concepts is, on this view, precisely what aesthetic judgment is not. Schopenhauer described aesthetic experience as a temporary release from ordinary conceptual judgment and practical concern (Schopenhauer, 1818/1969), and Dewey emphasized its unity and completeness (Dewey, 1934). From each of these perspectives, analysis that decomposes an experience into determinate parts may constrain, interrupt, or fragment it. Predictive-processing accounts converge on a similar prediction from a different direction: aesthetic pleasure arises in part from forming predictions and reducing uncertainty or prediction error (Van de Cruys & Wagemans, 2011; Muth & Carbon, 2013). Some analysis may facilitate this process, but analysis pursued further may expose new uncertainty rather than resolve it. Notably, this implies that the experimentally documented benefits of analysis and the theoretically anticipated costs may both hold, and may be reconciled by the depth at which the analysis operates. The present study therefore asks whether, under what conditions, and why the level of analysis produces differences in aesthetic experience — and in particular whether finer-grained analysis removes the aesthetic gain that analysis otherwise produces.

To address these questions, we manipulated the depth of analysis in an aesthetic task. According to construal level theory, the same object can be represented at different levels of abstraction: abstract representations are more general and more distant from direct experience, whereas concrete representations contain more detailed information and are closer to it (Trope & Liberman, 2010; see also Rosch et al., 1976; Vallacher & Wegner, 1987). Building on this distinction, we manipulated whether the information guiding analysis concerned a broad dimension of aesthetic experience or the specific subdimensions within that dimension. Aesthetic experience comprises three broad dimensions — perceptual, affective, and semantic (Leder et al., 2004; Chatterjee & Vartanian, 2014; Pelowski et al., 2017). The perceptual dimension comprises structure, color, spatiality, and complexity; the affective dimension comprises pleasure, impact, relaxation, and harmony; and the semantic dimension comprises abstractness versus realism, imaginative evocation, and profundity of meaning (Marković, 2012). Analysis directed at a broad dimension was defined as shallow analysis, and analysis directed at its constituent subdimensions as deep analysis. We emphasize that depth here refers to the level of abstraction at which the same information is construed, not to the amount of processing or the level of encoding in the sense of Craik and Lockhart (1972): participants in both conditions engaged the same class of semantic–affective information, and differed in whether it was construed as a single global evaluation or decomposed into separately reportable attributes.

Because different types of aesthetic objects may be differentially suited to different aesthetic dimensions, and because analysis depth may interact with both, we additionally manipulated image category. Viewing different image categories recruits partially distinct neural systems. Viewing artworks selectively engages the frontal pole, dorsomedial prefrontal cortex, and inferior frontal gyrus (Vessel et al., 2019); the dorsomedial prefrontal cortex and inferior frontal gyrus are implicated in emotional experience, evaluation, and regulation, and the frontal pole and adjacent anterior medial prefrontal regions have been associated with aesthetic value processing in visual art (Ochsner et al., 2002; Kober et al., 2008; Tabei et al., 2015; Hu et al., 2020). Viewing natural landscapes, in contrast, selectively engages the ventral occipitotemporal cortex (Isik & Vessel, 2021), which together with adjacent visual regions, supports the representation of perceptual information such as scene layout, object category, and visual form (Epstein & Kanwisher, 1998; Haxby et al., 2001; Isik & Vessel, 2021). These findings suggest a potential match between artworks and the affective dimension, and between natural landscapes and the perceptual dimension. More importantly, analysis depth may interact with image category and dimension: making affective responses more explicit through labeling or classification has been associated with attenuated subjective and neural emotional responding (Lieberman et al., 2007, 2011; Herbert & Kissler, 2010), whereas further perceptual processing may support more complete perceptual organization (Muth & Carbon, 2013), and additional semantic information may facilitate understanding and evaluation (Millis, 2001; Russell, 2003; Leder et al., 2006; Liu et al., 2022). Any cost of deeper analysis might therefore be expected in the artwork–affective combination in particular, rather than across conditions generally.

We further asked what neural change accompanies such a cost. Network-level analysis offers a useful vantage point on aesthetic experience. Existing work indicates that aesthetic experience engages both neural systems specialized for processing stimulus content and the default mode network (DMN), which is closely associated with self-referential and personally relevant processing (Buckner et al., 2008; Vessel et al., 2012; Andrews-Hanna et al., 2014; Yeshurun et al., 2021). Viewing different categories of visual stimuli selectively recruits different sensory-related systems (Vessel et al., 2019; Isik & Vessel, 2021), whereas the DMN represents aesthetic appeal across visual categories (Vessel et al., 2019). Critically, the two classes of system respond differently: activity in sensory-related regions varies continuously and gradually with aesthetic ratings, whereas DMN-related frontal regions respond in a step-like manner, primarily during the most intense aesthetic experiences (Vessel et al., 2012). Yet existing neural work has almost exclusively characterized these systems in the direction of aesthetic experience emerging, intensifying, or unfolding over time (Vessel et al., 2012, 2019; Belfi et al., 2019); how they change when aesthetic experience is instead weakened remains largely unexamined. A manipulation that removes an aesthetic gain therefore provides an opportunity to approach the same question from the direction of disruption. Given the distinct response profiles of the two systems, comparing how each changes when the gain is removed may indicate whether the loss arises in stimulus-specific content processing or in the higher-order integration supported by the DMN, and thereby test whether the DMN is critical to sustaining aesthetic experience.

A further consideration shapes how such a change should be measured. The function of a brain network depends on the specific spatial organization of its constituent subsystems and on the coordination among them (Andrews-Hanna et al., 2010), and dynamic changes in aesthetic experience have been shown to be reflected in DMN activity patterns (Belfi et al., 2019). The effect of deeper analysis on the DMN may accordingly appear not as a simple increase or decrease in overall activation, but as a change in how the network’s contribution is spatially distributed. We therefore propose that the loss of the aesthetic gain produced by shallow analysis arises from a change in the frequency-specific spatial organization of the DMN, and a consequent decoupling between DMN configuration and subjective aesthetic experience.

Across two experiments, we show that the benefit of analysis has a boundary in depth, and that this boundary is set by what is analyzed rather than by analysis itself. In Experiment 1, deeper analysis removed the aesthetic gain in one condition only — the affective dimension of visual art — while for natural landscapes, it enlarged the gain on all three dimensions. Experiment 2 replicated this dissociation in an independent sample and, using frequency-resolved generalized eigendecomposition of fNIRS data, found it accompanied by a selective rise in the spatial entropy of the DMN together with the loss of the network–behavior associations present under shallow analysis. On this evidence, what determines whether analysis helps is not how much processing it recruits but how the DMN organizes what is processed.

## 2. Experiment 1

### 2.1 Participants

Thirty-six undergraduate and graduate students (15 males, *M* = 22.32 years, *SD* = 2.27) participated in Experiment 1. Participants were randomly assigned to either the Shallow Analysis group or the Deep Analysis group, with 18 participants in each group. All participants reported normal or corrected-to-normal vision and provided informed consent prior to participation. They received appropriate compensation for their participation. The study was approved by the Ethics Committee of Nanjing University and was conducted in accordance with the Declaration of Helsinki (World Medical Association, 2013).

### 2.2 Materials

The visual stimuli consisted of two categories of color images: artworks and natural landscapes. The artwork images were selected from the Catalog of Art Museum Images Online (CAMIO) following the stimulus selection procedure described by Vessel et al. (2012), whereas the natural landscape images were selected from the Nature-related Image System (NIS) developed by Gao et al. (2025).

All images were presented as high-resolution color photographs with identical display parameters. Images containing obvious textual information, watermarks, or other distracting visual elements were excluded.

For each participant, the artwork and natural-landscape images were shuffled separately. Within each image category, eight images were assigned to the image-baseline task and 18 to the formal task. The 18 formal-task images were then allocated equally across the perceptual, affective, and semantic dimensions, with six images per dimension. The order of the two baseline blocks and the formal trials was randomized for each participant. The complete stimulus set is provided in Supplementary Material 1.

### 2.3 Experimental Design

Experiment 1 employed a 2 (analysis depth: shallow vs. deep, between-subjects) × 2 (image category: artwork vs. natural landscape, within-subjects) × 3 (analytical dimension: perceptual, affective, and semantic, within-subjects) mixed factorial design. Participants were randomly assigned to either the shallow group or the deep group, whereas image category and analytical dimension were manipulated within participants.

To quantify changes in aesthetic experience following analytical processing, all participants first completed an image-baseline task, during which they viewed artwork and natural landscape images without performing any analytical task and provided only overall aesthetic ratings.

During the formal experiment, each participant completed analytical evaluations under all six combinations of image category and analytical dimension (i.e., artwork–perceptual, artwork– affective, artwork–semantic, landscape–perceptual, landscape–affective, and landscape– semantic), with six trials in each condition (36 trials in total). To ensure that participants actively engaged in analytical processing rather than passive viewing, they were required to complete analytical judgments before providing an overall aesthetic rating.

Analysis depth was operationalized by manipulating the granularity of analytical judgments within each analytical dimension while keeping the analytical dimension itself constant. In the shallow-analysis condition, participants completed a single judgment corresponding to the assigned analytical dimension (i.e., visual expression, emotional expression, or semantic expression). In the deep-analysis condition, participants evaluated multiple predefined subdimensions before providing the overall judgment. Specifically, the perceptual dimension comprised structure, color, spatial composition, and visual complexity; the affective dimension comprised happiness, impact, relaxation, and harmony; and the semantic dimension comprised abstractness–realism, imagination, and meaningfulness. Thus, the two analysis depth conditions differed in the granularity of analytical processing while maintaining identical image stimuli and analytical dimensions.

The primary behavioral outcome was the baseline-corrected change in overall aesthetic ratings. For each participant, the mean aesthetic rating in each formal-task condition was calculated and then baseline-corrected by subtracting that participant’s mean rating from the corresponding image-category baseline block. The resulting change scores were compared between the shallow and deep groups.

#### 2.3.1 Apparatus and Setup

The experimental program was developed in Python using the Tkinter graphical user interface toolkit and the Pillow image-processing library for stimulus presentation and behavioral data collection.

Visual stimuli were presented on a 37-inch Samsung monitor (S37D702EAC) with a resolution of 3840 × 2160 pixels and a refresh rate of 120 Hz. Participants were seated approximately 80 cm from the display with their eyes aligned to the center of the screen. All images were presented centrally against a black background (RGB = 0, 0, 0). They were resized proportionally while preserving their original aspect ratios, with both width and height limited to 80% of the screen dimensions.

The experiment was conducted individually in a quiet laboratory under dim lighting conditions. The laboratory door remained closed during testing to minimize external distractions, and participants completed the task alone. Participants completed all responses using a computer mouse.

#### 2.3.2 Procedure

The experiment consisted of two consecutive phases: an image-baseline phase followed by a formal experimental phase (see **Figure 1**).

**Figure 1.**
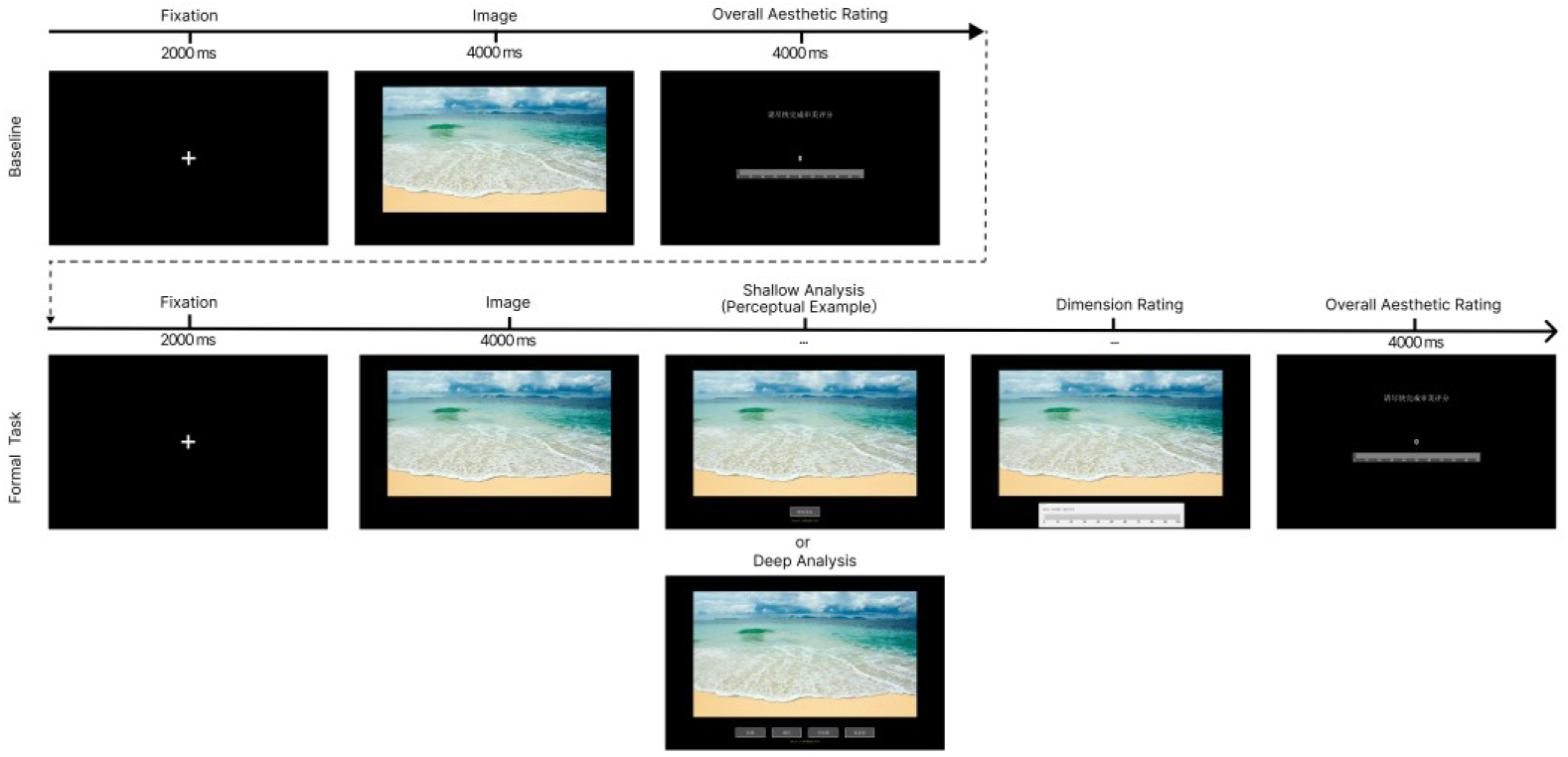
Experimental procedure of Experiment 1. Participants first completed the image baseline and then the formal task. In the formal task, the shallow group evaluated one overall dimension, whereas the deep group evaluated its specific subdimensions before providing an aesthetic rating.

During the image-baseline phase, participants completed two baseline blocks corresponding to the artwork and natural landscape image categories. The order of the two blocks was randomized across participants. Within each block, the eight baseline images were presented sequentially. Each trial began with a 2-s fixation cross, followed by a 4-s presentation of an image. Participants then rated the overall aesthetic quality of the image within 4 s using a continuous on-screen slider ranging from 0 (“not at all beautiful”) to 100 (“extremely beautiful”). A 2-s fixation cross preceded the next trial. Participants were instructed to view the images naturally without performing any analytical task. A 15-s rest period was provided between the two baseline blocks, followed by another 30-s rest period before the formal experimental phase.

During the formal experimental phase, 36 images (18 artworks and 18 natural landscapes) were presented in a randomized order. Each trial began with a 2-s fixation cross, followed by a 4-s presentation of the image. The image remained visible while participants completed the assigned analytical judgment(s). Participants in the shallow group completed one analytical judgment corresponding to the assigned analytical dimension, whereas participants in the deep group successively completed multiple subdimension judgments within the same analytical dimension. After completing the analytical task, participants rated the overall aesthetic quality of the image within 4 s using the same 0–100 on-screen slider. Each trial ended with a 2-s fixation cross before the next trial. Trial order was randomized independently for each participant.

#### 2.3.3 Behavioral Data Acquisition

During the formal experiment, the program automatically recorded participants’ ratings for each analytical question together with the corresponding completion time. These completion times were recorded solely to verify that participants completed the required analytical task and were not included in subsequent statistical analyses.

For the overall aesthetic evaluation, both the aesthetic rating and the corresponding completion time were recorded. Participants were required to complete the aesthetic rating within the specified time window. Ratings without a recorded response latency were treated as missing during preprocessing. All behavioral data were automatically exported as comma-separated value (CSV) files for subsequent statistical analyses.

#### 2.3.4 Data Analysis

Analyses were performed in R. For each participant, the mean rating in each image category × Dimension condition was baseline-corrected by subtracting the mean rating from the corresponding image-category baseline block. The baseline-corrected aesthetic-rating change was submitted to a 2 (image category: Visual Art vs. Natural Landscapes) × 3 (Dimension: Perceptual, Semantic, Affective) × 2 (analysis depth: Shallow vs. Deep Analysis) mixed-design ANOVA (Type III sums of squares), with image category and Dimension as within-subject factors, and analysis depth as a between-subjects factor. The Greenhouse–Geisser correction was applied where sphericity was violated (Greenhouse & Geisser, 1959), and partial eta-squared (*η*^2^) was reported as the effect-size estimate. Because Experiment 1 was designed to *screen* for the specific condition in which deeper processing yields a smaller change than shallow processing (Deep < Shallow), we decomposed the three-way interaction with simple-effect contrasts comparing Deep vs. Shallow Analysis within each image category × Dimension cell. These planned contrasts served as a hypothesis-generating screen and are reported uncorrected; the single cell meeting the Deep < Shallow criterion was carried forward as the a priori target condition for Experiment 2.

In addition, to characterize what each condition did relative to passive viewing rather than relative to the other group, the change score in each image category × Dimension × Group cell was tested against zero with a one-sample *t*-test. Because a non-significant test cannot by itself support the claim that a condition produced no change, each non-significant cell was followed by two one-sided tests of equivalence (TOST; Lakens, 2017; Lakens et al., 2018) against symmetric bounds of ±5 rating points, i.e. 5% of the 0–100 scale. This bound is smaller than the largest change observed anywhere in Experiment 1 (5.83 points) and is therefore conservative with respect to the effects the design was intended to detect. To allow readers to apply their own criterion, we additionally report for each cell the smallest symmetric bound at which equivalence would be established, which is equivalent to the wider limit of the 90% confidence interval.

### 2.4 Results

The baseline-corrected aesthetic-rating change, defined as the mean formal-task rating minus the mean rating from the corresponding image-category baseline block, was entered into a 2 (image category: Visual Art [VA] vs. Natural Landscapes [NL]) × 3 (Dimension: Perceptual, Semantic, Affective) × 2 (analysis depth: Shallow Analysis vs. Deep Analysis) mixed-design ANOVA, with image category and Dimension as within-subject factors, and analysis depth as a between-subjects factor (see **Figure 2**).

**Figure 2.**
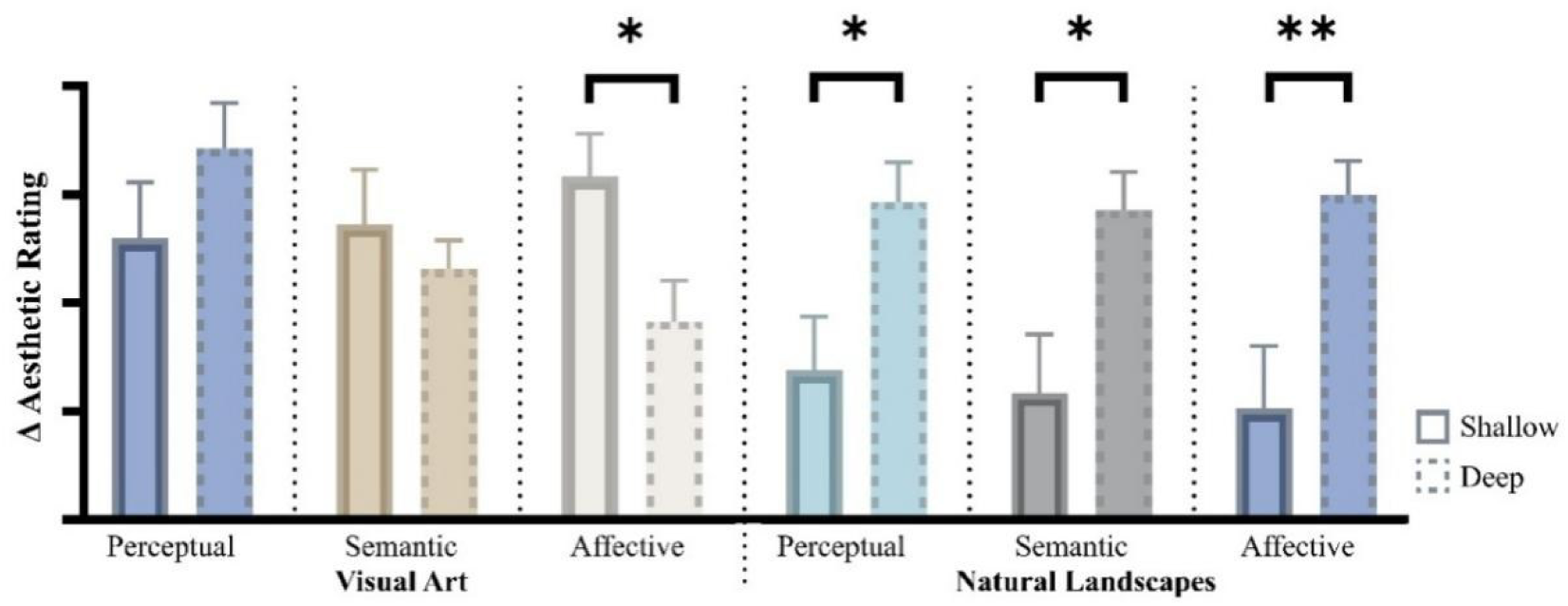
Aesthetic-rating change across image categories, dimensions, and analysis depth (Experiment 1). Mean baseline-corrected aesthetic-rating change for Visual Art (VA) and Natural Landscapes (NL) across the Perceptual, Semantic, and Affective dimensions, shown separately for the Shallow and Deep Analysis groups (*N* = 36). Deeper analysis selectively reduced the affective aesthetic-rating change for VA (Deep < Shallow) while enhancing the change for NL, reflecting the significant image category × Dimension × analysis depth interaction. Error bars denote SE. Asterisks denote uncorrected simple-effect contrasts (* *p* < 0.05, ** *p* < 0.01).

Mauchly’s test indicated a violation of sphericity for the Dimension factor, *W* = 0.82, χ²(2) = 6.71, *p* = 0.035; Greenhouse–Geisser corrected degrees of freedom are therefore reported for all effects involving Dimension. The between-subjects main effect of analysis depth was only a nonsignificant trend, *F*(1, 34) = 3.53, *p* = 0.069, ηp² = 0.094. Critically, analysis depth interacted with image category, *F*(1, 34) = 8.12, *p* = 0.007, ηp² = 0.193, further qualified by a significant image category × Dimension × analysis depth interaction, *F*(1.79, 60.75) = 5.98, *p* = 0.006, ηp² = 0.15. No other effect reached significance (all *p* ≥ 0.095).

To decompose the three-way interaction, we contrasted Deep vs. Shallow Analysis within each image category × Dimension cell, seeking the condition in which deeper analysis produced a smaller change in aesthetic rating (Deep < Shallow) to serve as the a priori target for Experiment 2. This pattern emerged selectively for Visual Art on the Affective dimension, Deep − Shallow = −6.69, *t*(34) = −2.42, *p* = 0.021, *d* = −0.81; the Semantic and Perceptual dimensions did not differ (Semantic: −2.06, *t*(34) = −0.73, *p* = 0.469, *d* = −0.24; Perceptual: +4.12, *t*(34) = 1.24, *p* = 0.223, *d* = 0.41). For Natural Landscapes, the direction reversed, deeper analysis producing larger positive changes in all three dimensions (Affective: +9.87, *t*(34) = 3.01, *p* = 0.005, *d* = 1.00; Semantic: +8.43, *t*(34) = 2.61, *p* = 0.013, *d* = 0.87;

Perceptual: +7.74, *t*(34) = 2.54, *p* = 0.016, *d* = 0.85). Deep analysis thus selectively suppressed the affective enhancement elicited by Visual Art while amplifying the change for Natural Landscapes, indicating that the aesthetic cost of deeper analysis is domain-specific. We therefore adopted the Visual Art × Affective condition as the focus of Experiment 2.

#### Change relative to the image baseline

The contrasts above compare the two groups with one another; to establish what each group did relative to passive viewing, each cell was additionally tested against zero. In the shallow group, a single cell differed from baseline: for Visual Art on the Affective dimension, ratings rose by 5.83 points, *t*(17) = 2.92, *p* = 0.01, *d* = 0.6. No other cell reached significance (|*t*| ≤ 1.69, all *p* ≥ 0.109). In the deep group, the pattern differed. Ratings rose above baseline for Visual Art on the Perceptual dimension, *M* = 7.13, *t*(17) = 3.39, *p* = 0.004, *d* = 0.8, and for all three Natural Landscape dimensions (Perceptual: *M* = 4.66, *t*(17) = 2.55, *p* = 0.021, *d* = 0.6; Semantic: *M* = 4.28, *t*(17) = 2.45, *p* = 0.026, *d* = 0.58; Affective: *M* = 5.00, *t*(17) = 3.22, *p* = 0.005, *d* = 0.76). The two Visual Art cells in which deep analysis produced no gain were both statistically equivalent to zero: Affective, *M* = −0.86, TOST *p* = 0.022; Semantic, *M* = 1.58, TOST *p* = 0.010. For the Affective cell — the condition carried forward to Experiment 2 — equivalence held down to a bound of ±4.18 points (*d* = 0.52). Deep analysis of Visual Art on the Affective dimension therefore did not push aesthetic ratings below the image baseline; it left them at baseline, removing the increase that shallow analysis produced in the same condition. These twelve tests decompose an already-significant three-way interaction, and per-cell statistics are given in **Table S1**.

## 3. Experiment 2

### 3.1 Participants

Participants were assigned between subjects to a Deep Analysis group or a Shallow Analysis group, which differed in the depth of emotional processing engaged during aesthetic evaluation (for the depth manipulation, see Experimental Design, Section 3.3, and Procedure, Section 3.3.2). Eighteen participants were analyzed in the deep group (7 males, *M* = 24.3 years, *SD* = 4.3). In the shallow group, one participant was excluded prior to analysis owing to a data-acquisition/equipment failure during recording, leaving 16 participants (6 males, *M* = 22.6 years, *SD* = 1.8); one further participant was removed from the network-level analyses because the residual eigenspectrum was non-finite across frequencies (see Data Acquisition and Analysis, subsection d), yielding 15 shallow participants for those analyses.

Because the behavioral, network, and brain–behavior analyses have different data requirements, they draw on partially non-overlapping subsamples:

Behavioral replication (*N* = 34; deep *n* = 18, shallow *n* = 16). Includes every participant with a valid aesthetic-change score, irrespective of fNIRS quality; the shallow participant dropped from the network analyses for a non-finite residual eigenspectrum is retained here because the behavioral response was usable.

Network-level analyses (*N* = 33; deep *n* = 18, shallow *n* = 15). Includes every participant with usable fNIRS data, irrespective of behavioral quality.

Brain–behavior analyses (*N* = 31; deep *n* = 17, shallow *n* = 14). These require paired neural and behavioral data. Of the 33 participants in the network sample, one per group had usable fNIRS data but no usable aesthetic-change score — the behavioral response was lost to a mid-session equipment failure while the neural recording was retained — and therefore could not be entered into the correlations; these two were removed list-wise.

All participants provided written informed consent, and the study was approved by the Ethics Committee of Nanjing University and conducted in accordance with the Declaration of Helsinki (World Medical Association, 2013).

### 3.2 Materials

The visual stimuli consisted of 26 artwork images and eight abstract patterns. The artwork images were selected from the same source as in Experiment 1, following the procedure described by Vessel et al. (2012). The abstract patterns were selected from the stimulus set developed by Jacobsen and Höfel (2002) and served as non-aesthetic control stimuli in the cognitive-baseline task.

For each participant, eight artwork images were assigned to the image-baseline task and 18 to the formal task. The eight abstract patterns were used in the cognitive-baseline task. Image assignment and presentation order were randomized independently for each participant. The complete set of experimental stimuli is provided in the Supplementary Material 1.

### 3.3 Experimental Design

Experiment 2 employed a one-factor between-subjects design, in which analysis depth (shallow vs. deep) was manipulated between participants.

Based on the findings of Experiment 1, only the artwork–affective analysis condition was included in Experiment 2, as this was the only condition in which analysis depth significantly modulated baseline-corrected aesthetic ratings. Accordingly, Experiment 2 focused on this condition to investigate the neural mechanisms underlying the behavioral effect of analysis depth.

To ensure that participants actively engaged in affective analysis rather than passive viewing, they were required to complete affective judgments before providing an overall aesthetic rating. In the shallow-analysis condition, participants completed a single judgment of the emotional expression of each artwork. In the deep-analysis condition, participants successively evaluated four affective subdimensions (happiness, impact, relaxation, and harmony) before providing the overall judgment. Thus, the two groups differed in the granularity of affective analysis while viewing identical image stimuli.

Before the formal experimental task, all participants completed an image-baseline task and a cognitive-baseline task. The primary behavioral outcome was the baseline-corrected change in aesthetic ratings, calculated for each participant as the mean rating in the formal artwork task minus the mean rating in the artwork image-baseline block. These change scores were subsequently compared between the shallow and deep groups.

#### 3.3.1 Apparatus and Setup

The experimental apparatus, stimulus presentation, and laboratory setting were identical to those used in Experiment 1. In addition, cortical hemodynamic activity was continuously recorded using a NIRSport2 functional near-infrared spectroscopy (fNIRS) system (NIRx Medical Technologies LLC, Glen Head, NY, USA). The system employed dual wavelengths of 760 and 850 nm and sampled the signals at 16.7 Hz. The probe set consisted of 16 light sources and 16 detectors, forming 57 measurement channels covering the bilateral frontal, temporal, parietal, and occipital cortices. Based on signal quality assessment, 28 channels were retained for subsequent analyses. These channels were grouped into nine predefined regions of interest (ROIs): the left and right lateral temporal cortex (LTC), left and right inferior frontal gyrus (IFG), orbitofrontal cortex (OFC), frontal pole (FP), left and right inferior parietal lobule (IPL), and the lateral occipito-parietal area (LOPPA). Oxygenated hemoglobin (HbO) signals were used for all subsequent analyses (see Supplementary Material 2 for detailed coordinates).

#### 3.3.2 Procedure

The experiment consisted of three consecutive phases: an image-baseline phase, a cognitive-baseline phase, and a formal experimental phase.

During the image-baseline phase, participants viewed eight artwork images without performing any analytical task and rated the overall aesthetic quality of each image.

During the cognitive-baseline phase, participants completed the affective-analysis task on eight abstract patterns. Each cognitive-baseline trial followed the same sequence as a formal-task trial. This phase provided a neural baseline for affective processing without artwork content.

During the formal experimental phase, 18 artwork images were presented in a randomized order. Each trial began with a 2-s fixation cross and a 4-s presentation of the artwork. The artwork remained visible during the affective judgment, followed by a 4-s aesthetic rating. Participants in the shallow group completed a single affective judgment of the emotional expression of each artwork, whereas participants in the deep group successively completed judgments of the four affective subdimensions (happiness, impact, relaxation, and harmony) before providing the overall aesthetic rating. Trial order was randomized independently for each participant (Figure 3).

**Figure 3.**
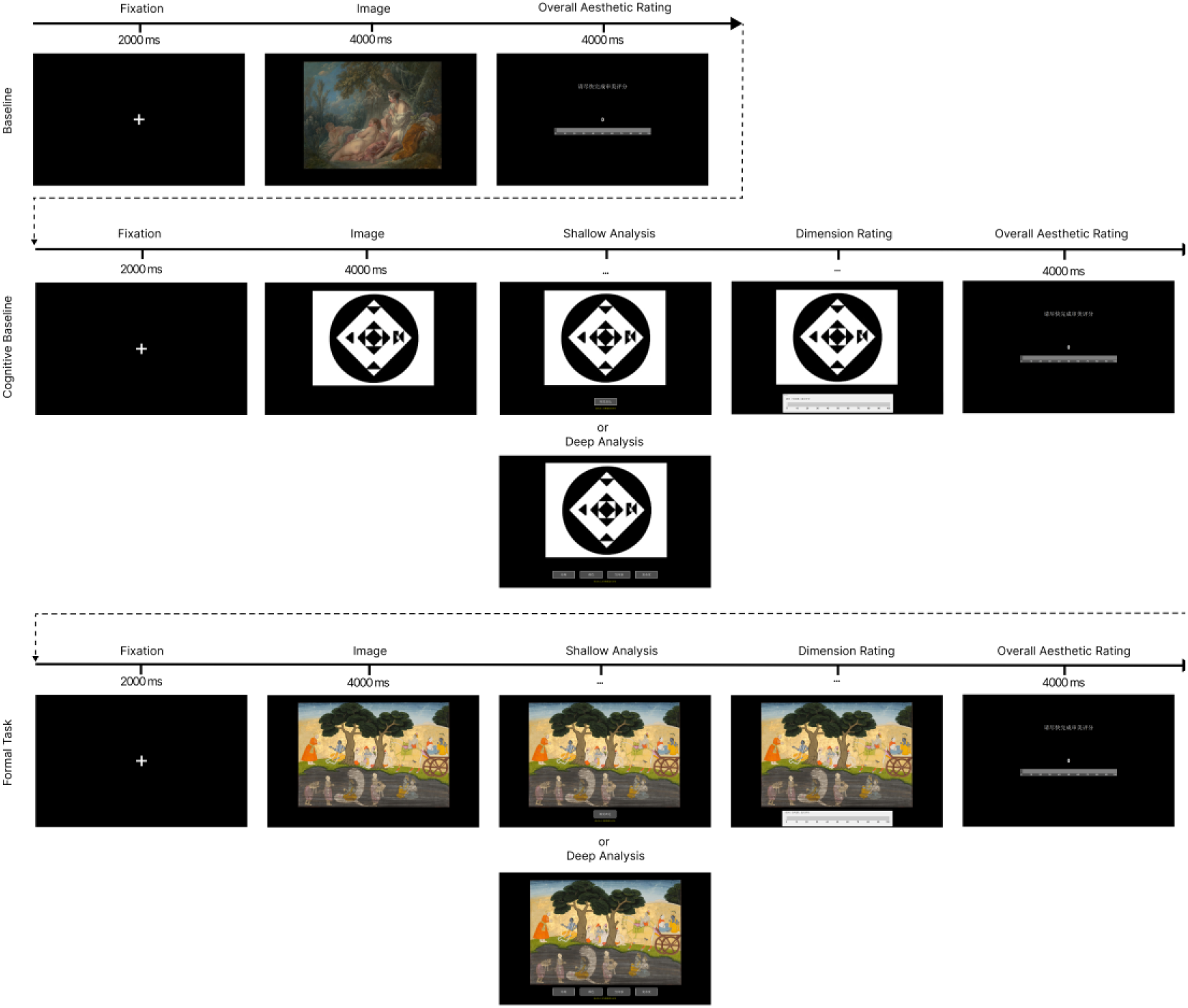
Experimental procedure of Experiment 2. Participants completed the image baseline, cognitive baseline, and formal artwork task. The shallow group made one overall affective judgment, whereas the deep group evaluated four affective subdimensions before providing an aesthetic rating.

#### 3.3.3 Data Acquisition and Analysis

##### Behavioral Data

The behavioral index was a baseline-corrected aesthetic-rating change score, calculated for each participant as the mean aesthetic rating in the formal artwork task minus the mean rating in the artwork image baseline block. Positive values denote a processing-induced increase in aesthetic evaluation. This change-score definition is identical to that used in Experiment 1. Behavioral scores were available for 31 of the 33 participants entering the network analyses (deep *n* = 17, shallow *n* = 14); the remaining two — one per group — had usable fNIRS data but no usable behavioral score (see Section 3.1), and were therefore excluded from the brain–behavior analyses only. Each group was also compared with the image baseline by a one-sample t-test, followed by two one-sided tests of equivalence against the ±5-point bound used in Experiment 1 (see Section 2.3.4); the smallest bound at which equivalence would be established is also reported. Bayesian one-sample t-tests were computed with a default JZS prior (*r* = 0.707; Rouder et al., 2009) as a complement to the equivalence tests.

##### fNIRS Data

###### (a) Preprocessing

Hemodynamic activity was recorded with continuous-wave functional near-infrared spectroscopy and sampled at 16.7 Hz. Preprocessing was performed in MATLAB with HOMER-based routines, in the following order:

1. Concentration time courses were converted to optical density (hmrConc2OD) and back to light intensity, and channels were re-ordered to match the probe (SD) geometry.
2. Poor channels were pruned (enPruneChannels) using a dynamic-range criterion of [0 3], a signal-to-noise-ratio threshold of 2, and an admissible source–detector separation of [0 50] mm.
3. Intensity was converted to optical density (hmrIntensity2OD).
4. Global physiological noise was attenuated with a spatial principal-component filter retaining the components that explained 80% of the variance (enPCAFilter, nSV = 0.8).
5. Optical density was converted to hemoglobin concentration (hmrOD2Conc).
6. Head-motion artifacts were corrected with the correlation-based signal improvement method (hmrMotionCorrectCbsi).

Oxygenated hemoglobin (HbO) was retained for all subsequent analyses.

Critically, no band-pass filter was applied at any preprocessing stage. Because the frequency-resolved analysis below is designed to identify functionally relevant frequencies in a data-driven manner across the full 0.01–1 Hz range, pre-selecting a frequency band would impose a priori spectral assumptions and undermine that rationale. Frequency specificity was instead achieved through the narrow-band generalized-eigendecomposition (GED) scan (see subsection b).

The 28 channels were grouped into nine regions of interest (ROIs) for the subsequent subnetwork analyses: the left and right LTC, left and right IPL, left and right IFG, OFC, FP, and left LOPPA (see subsection g).

###### (b) FREQ-NESS-based network extraction

Frequency-specific multivariate networks were estimated with a frequency-resolved generalized eigendecomposition pipeline following FREQ-NESS (Rosso et al., 2025), using GED as the source-separation engine (Cohen, 2022). For each participant and block, the 28-channel time series was amplitude-normalized by an order-of-magnitude rescaling to condition the covariance estimates.

A broadband reference covariance matrix **R** was computed from the rescaled data and shrinkage-regularized:

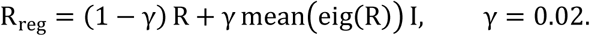

The analysis swept 496 center frequencies from 0.01 to 1 Hz in 0.002-Hz steps (following Sun et al., 2026). At each center frequency *f*, a narrow-band signal covariance **S(***f*) was computed from the data band-pass filtered at *f* with a frequency-domain Gaussian filter (filterFGx; Cohen, 2014) of full width at half maximum (FWHM) 0.02 Hz; for the five ultra-low frequencies below 0.02 Hz, the FWHM was reduced to 0.01 Hz to prevent leakage across 0 Hz. The generalized eigendecomposition

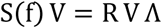

was solved at each frequency, yielding spatial filters (generalized eigenvectors **V**) ordered by their generalized eigenvalues *λ*; the leading 10 components were retained. Eigenvalues were expressed as a percentage of their sum across components,

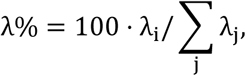

to index relative network prominence.

For each component, the channel-space activation pattern (topography) was recovered as the forward model **a** = **S v** and sign-normalized to its maximum magnitude channel. Activation patterns, rather than the spatial filters themselves, were used for all topographic interpretation: filter weights are not interpretable as topographies, because a large filter weight may serve to suppress noise rather than to reflect a source, whereas the activation pattern reflects the projection of the component onto channel space (Haufe et al., 2014; Cohen, 2022).

###### (c) Effective dimensionality and component-number selection

To set the number of the nuisance subspaces removed from the formal task, the effective dimensionality (ED) of each baseline GED spectrum was quantified by the participation ratio:

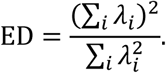

ED was computed per participant per frequency and averaged across frequencies; the group-level ED was the across-participant mean, and the number of components removed was the ceiling of this mean (a conservative choice). This produced *K*_1_ from the image baseline (pooled across groups), and, because analysis depth applies to the cognitive baseline, group-specific values *K*_2,deep_ and *K*_2,shallow_ from cognitive baseline (computed within the deep and shallow groups).

###### (d) Dual-subspace projection and residual GED of the task network

To isolate task-specific network activity, at each frequency and within each participant, the joint subspace spanned by the leading baseline components was removed before the task network was re-estimated. The top-*K*_1_ eigenvectors of the image baseline (**W**_1_) and the top-*K*_2_ eigenvectors of the cognitive baseline (**W**_2_; *K*_2_=*K*_2,deep_ or *K*_2,shallow_ according to group) were concatenated, and an orthonormal basis **Q** of their joint span was obtained by orthogonalization,

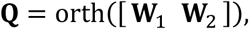

which automatically accommodates any overlap or rank deficiency between the two baseline subspaces; the number of dimensions actually removed, rank([**W**_1_ **W**_2_]), was recorded per participant and frequency. The residual projector

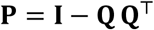

projects onto the orthogonal complement of the baseline subspace.

The narrow-band-filtered formal task data at each frequency were projected through **P**; a residual signal covariance **S**_res_ was computed on the projected data (with a 10^−6^ **I** ridge for numerical stability); and a residual GED,

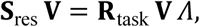

was solved against the shrinkage-regularized broadband task reference. This produced the residual task eigenvalue spectrum (normalized to *λ*%) and the residual component-1 topography, a =**S**_res_ **v**. For the before-removal comparison condition, the identical pipeline was run with **P** set to the identity, so that pre- and post-removal spectra differ only by the subspace-removal step. Participants whose residual component-1 spectrum was non-finite across frequencies were excluded as a data-quality criterion (one shallow participant; see Participants, Section 3.1). Because the two baseline subspaces were disjoint in every participant and at every frequency, the number of dimensions actually removed was constant within each group — 10 in the deep group (*K*₁ = 4, *K*₂,deep = 6) and 9 in the shallow group (*K*₁ = 4, *K*₂,shallow = 5) — leaving residual subspaces of 18 and 19 dimensions respectively.

###### (e) Statistical analysis I — task component-1 eigenvalue (deep vs shallow)

We tested whether the leading residual task network (component 1) differed between groups as a function of frequency, using the post-removal component-1 eigenvalue (*λ*%) at each of the 496 frequencies. At every frequency, an independent-samples *t* statistic (deep vs shallow) was computed, and multiple comparisons across frequencies were controlled with a cluster-based permutation test (Maris & Oostenveld, 2007). Frequencies whose |*t*| exceeded the two-tailed parametric threshold (*α* = 0.05, df = *n*_deep_+*n*_shallow_−2 = 31) were grouped into contiguous clusters; only clusters spanning at least 6 consecutive frequencies (≥ 0.012 Hz) were retained, with cluster mass defined as the summed *t*. A null distribution was constructed by permuting the group labels 2,000 times and recording, on each permutation, the largest positive and the largest negative cluster mass; a cluster’s Monte-Carlo *p* value was the proportion of permutations whose maximal signed cluster mass equaled or exceeded the observed value. Clusters with *p* < 0.05 were considered significant, and within-cluster effect size was indexed by Cohen’s *d* on the pooled within-cluster values. This analysis used 18 deep and 15 shallow participants.

###### (f) Statistical analysis II — channel spatial weights and brain–behavior correlation

Topographic brain–behavior associations were assessed from the post-removal component-1 activation pattern. For each participant, the pattern was averaged across the 20 frequencies of the significant band (0.050–0.088 Hz; see Section 3.4) and unit-L2-normalized, retaining the direction of the topography while discarding its overall magnitude. The magnitude was discarded because it is not comparable across participants in fNIRS (differential pathlength, optode coupling, partial-volume effects) and because, in GED, network prominence is carried by the eigenvalue rather than by the pattern norm (Cohen, 2022); empirically, pre-normalization norms spanned more than two orders of magnitude across participants. The pattern sign was preserved because the FREQ-NESS sign convention was applied within participant, frequency, and component. No cross-participant sign alignment was performed, because aligning signs to a group template derived from the same data would introduce circularity (Kriegeskorte et al., 2009). For each channel, the signed normalized weight was correlated across participants with the behavioral aesthetic-change score using Spearman correlation, with *p* values obtained from 5,000 sign-/label-preserving permutations of the behavioral vector; correlations were run within each group and on the pooled sample.

###### (g) Statistical analyses III–IV — subnetwork normalized entropy, energy, and group differences

The 28 channels were partitioned a priori into a default-mode (DMN) subnetwork — bilateral LTC and bilateral IPL — and a domain-specific subnetwork — bilateral IFG, OFC, and FP — following the default-mode vs domain-specific dissociation of aesthetic processing reported by Vessel et al. (2019).

For each participant, the component-1 activation pattern was averaged across the significant band and unit-L2-normalized (as in subsection *f*), and two complementary summaries were computed for each subnetwork:

- **Distributedness (normalized Shannon entropy).** With *w_i_*=|*a_i_*| and *p_i_*=*w_i_*/ ∑*_j_ w_j_* over the subnetwork’s channels,

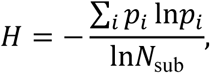

where *N*_sub_ is the number of channels in the subnetwork (Shannon, 1948). *H* ranges from 0 (activity concentrated on a single channel) to 1 (uniform engagement), indexing how distributed vs. focal the subnetwork’s contribution to the leading task network is.

- **Engagement (relative energy).** 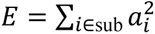 on the unit-norm pattern, i.e. the proportion of total component-1 energy carried by the subnetwork. Because the pattern is unit-norm, the DMN and domain-specific energies are near-complementary; we therefore also report a single network-bias index (DMN minus domain-specific energy) to summarize the relative weighting of the two subnetworks without double-counting.

###### Group differences in entropy (Analysis IV)

For each subnetwork, normalized entropy was compared between the deep and shallow groups using a permutation test: the statistic was the difference in group means (deep − shallow), evaluated against a null built by reassigning group labels 5,000 times; the two-tailed *p* value was the proportion of permutations with |null| ≥ |observed|. Cohen’s *d* (pooled *SD*) indexed effect size, and a Mann–Whitney *U* (rank-sum) test was reported as a small-sample-robust complement.

###### Brain–behavior correlation (Analysis III)

Within each group, subnetwork entropy and subnetwork energy were each correlated with the behavioral aesthetic-change score (Spearman, 5,000 permutations). To test whether the energy–behavior correlation itself differed by group, we additionally fit a moderation model (behavior ∼ group + energy + group × energy, energy *z*-scored) and tested the interaction by permutation (5,000), complemented by a direct permutation test on the difference in group-wise slopes.

### 3.4 Results

#### Behavioral Data

In an independent sample (*N* = 34; Deep Analysis *n* = 18, Shallow Analysis *n* = 16), we tested the effect isolated in Experiment 1 — the baseline-corrected aesthetic-rating change for Visual Art on the Affective dimension — comparing the Deep and Shallow Analysis groups with a Welch-corrected independent-samples *t*-test (**Figure 4**). Replicating Experiment 1, Shallow Analysis raised affective aesthetic ratings well above baseline, whereas Deep Analysis abolished this increase, leaving ratings essentially unchanged (Shallow *M* = 9.19; Deep *M* = −1.2; mean difference [Deep − Shallow] = −10.39, *SE* = 4.32, 95% CI [−19.19, −1.59]; Welch *t*(31.70) = −2.41, *p* = 0.022, two-tailed; Cohen’s *d* = −0.83).

**Figure 4.**
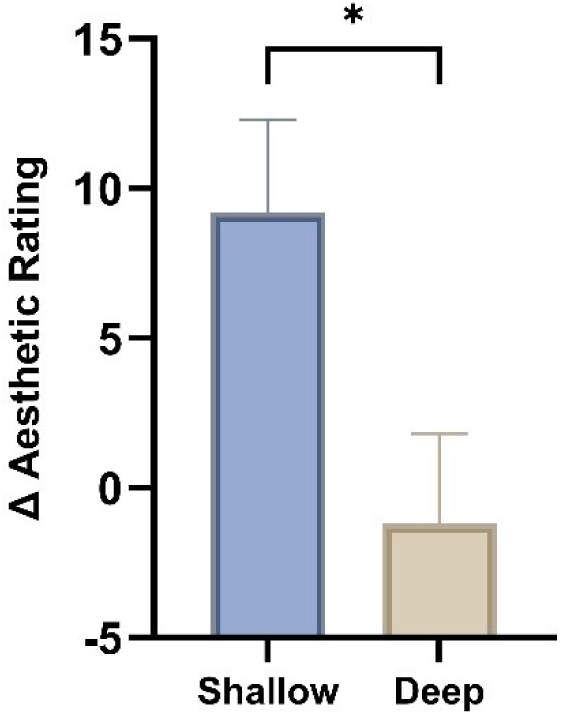
Replication of the VA × Affective effect. Baseline-corrected aesthetic-rating change for Visual Art on the Affective dimension in the Deep and Shallow Analysis groups. Shallow Analysis raised ratings above baseline, whereas Deep Analysis abolished this increase (Welch *t*(31.70) = −2.41, *p* = 0.022, two-tailed; Cohen’s *d* = −0.83). Error bars denote SE.

Each group was also tested against the image baseline. Shallow Analysis produced ratings significantly above baseline, *M* = 9.19, *t*(15) = 2.96, *p* = 0.01, *d* = 0.74, 95% CI [2.57, 15.81], closely reproducing the corresponding effect in Experiment 1 (*d* = 0.69). Deep Analysis left ratings at baseline, *M* = −1.20, *t*(17) = −0.4, *p* = 0.694, *d* = −0.09, 95% CI [−7.53, 5.13].

Equivalence testing against the ±5-point bound used in Experiment 1 was not conclusive in this sample (TOST *p* = 0.111); equivalence to zero was established from a bound of ±6.42 points (*d* = 0.5) upward, which remains smaller than the gain shallow analysis produced in the same experiment. Testing the deep group against a bound set at that gain (±9.19 points) yielded equivalence, TOST *p* = 0.008, as did the corresponding test in Experiment 1 (bound ±5.83, TOST *p* = 0.009). Bayesian one-sample tests (JZS prior, *r* = 0.707) gave converging and consistent evidence across the two experiments: moderate support for the null in the deep groups (BF₀₁ = 3.76 and 3.83) and moderate support for a change in the shallow groups (BF₁₀ = 5.49 and 5.58).

Deep analysis therefore did not reduce aesthetic ratings below the level obtained under passive viewing; it removed the increase that shallow analysis produced, in both samples.

#### fNIRS Data

##### Effective dimensionality and component numbers

Effective dimensionality (participation ratio) of the baseline GED spectra yielded the component numbers used for subspace removal (**Figure 5**). For the image baseline, ED (K₁) = 4. For the cognitive baseline, ED was computed within each group: deep ED (K_2_) = 6; shallow ED (K_3_)= 5.

**Figure 5.**
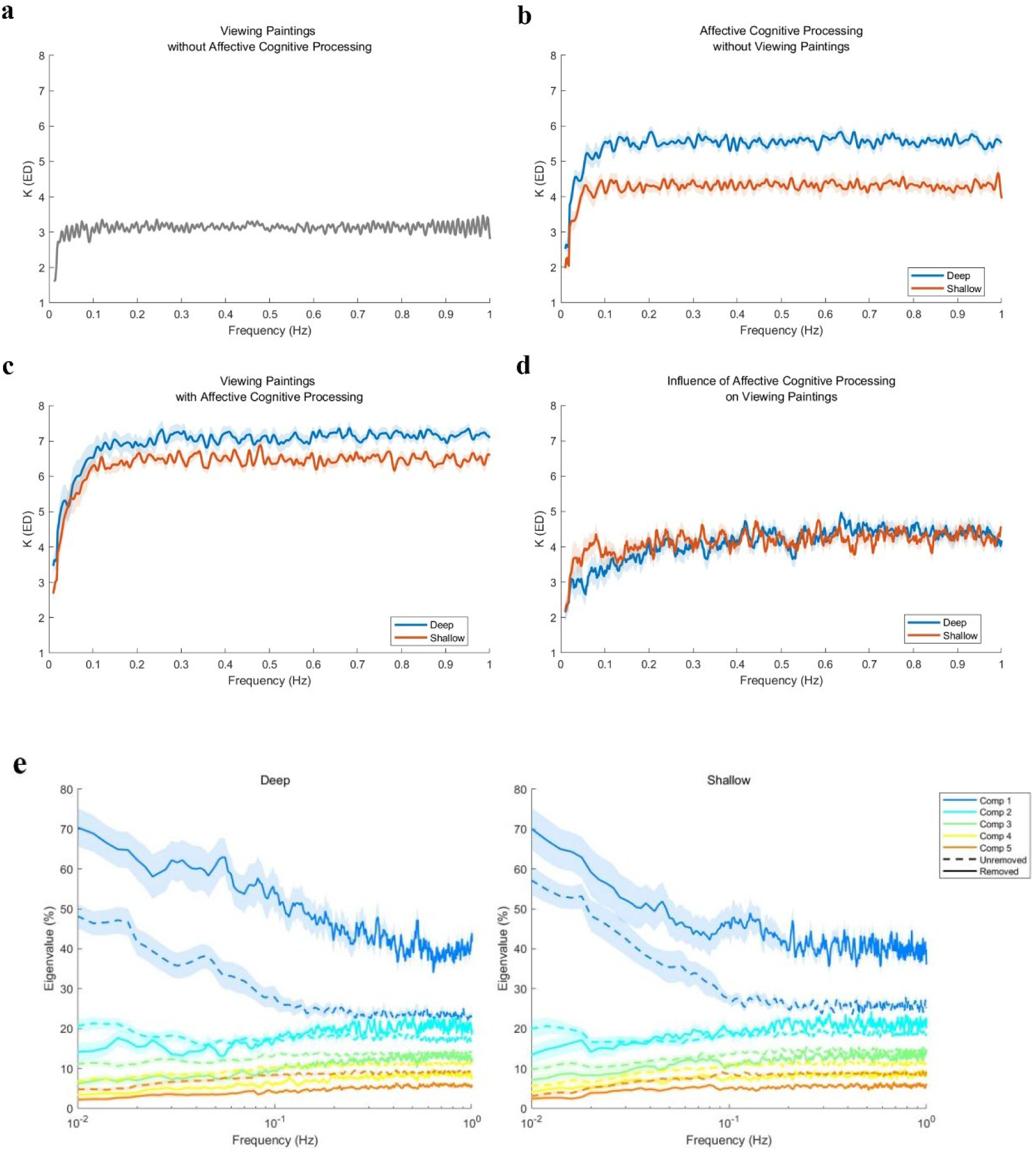
Residual task network after dual-subspace projection (Experiment 2, fNIRS). **(a–d)** GED eigenvalue spectra, effective-dimensionality estimates across frequencies: viewing VA without processing (a), processing without viewing VA (b), viewing VA with processing (c) and pure ED of processing on VA (d). **(e)** Normalized component-1 eigenvalue (λ%) of the task state before versus after removal of the baseline subspace: the leading residual component is enhanced after subspace removal.

The two baseline subspaces were disjoint in every participant and at every frequency, so rank ([**W**₁ **W**₂]) was constant within group: 10 in the deep group and 9 in the shallow group, leaving residual subspaces of 18 and 19 dimensions respectively. Because normalized entropy is computed over a fixed set of subnetwork channels, this one-dimension asymmetry does not enter the entropy normalization directly; its potential indirect influence on the recovered topography is evaluated by simulation in Supplementary Note S1.

##### Task component-1 eigenvalue: group comparison

The cluster-based permutation test on the post-removal component-1 eigenvalue identified a single significant cluster spanning 0.050–0.088 Hz (20 frequencies; cluster-mean *t* = 2.7, *p* = 0.004), within which the component-1 eigenvalue was higher in the deep than in the shallow group (Cohen’s *d* = 0.99; deep *n* = 18, shallow *n* = 15). No other cluster reached significance. This band (0.050–0.088 Hz) was used as the frequency window of interest for all subsequent topographic analyses (**Figure 6**).

**Figure 6.**
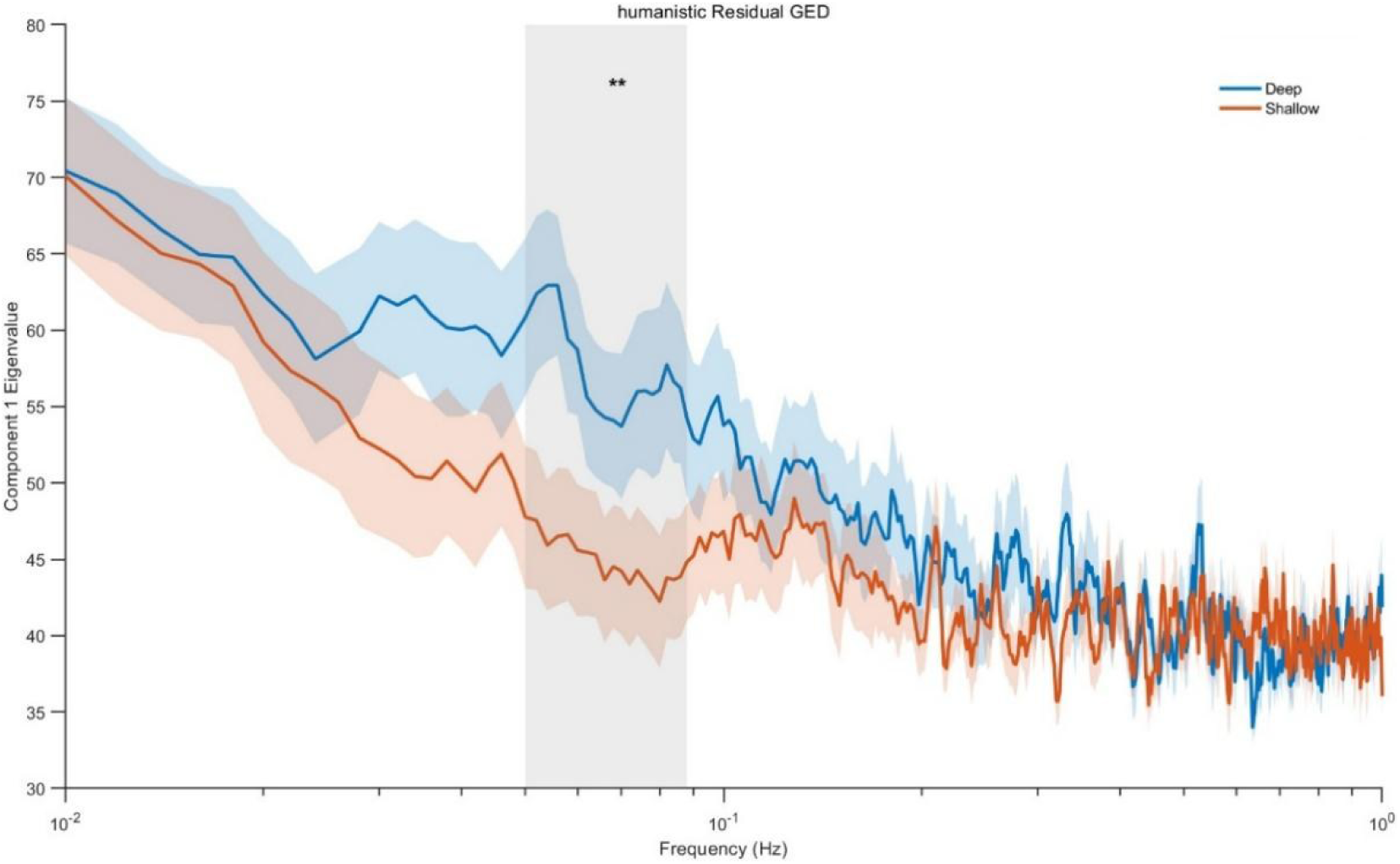
Frequency-resolved group comparison of the task component-1 eigenvalue. Post-removal component-1 eigenvalue (λ%) as a function of frequency (0.01–1 Hz) for the Deep and Shallow Analysis groups (deep *n* = 18, shallow *n* = 15). Cluster-based permutation testing identified a single significant cluster at 0.050–0.088 Hz (20 frequencies; cluster-mean *t* = 2.7, *p* = 0.004; Cohen’s *d* = 0.99), where the eigenvalue was higher in the deep group. Shading marks the significant cluster. Lighter color areas denote SE.

##### Channel spatial weights and brain–behavior correlation

Within the deep group (*n* = 17), the signed normalized component-1 spatial weights of two DMN channels were negatively correlated with the aesthetic-change score: left LTC (CH 7) (ρ = −0.561, *p* = 0.022) and left inferior parietal lobule (CH 40) (ρ = −0.498, *p* = 0.042); right LTC showed a sub-threshold trend (ρ = −0.424, *p* = 0.093). No channel in the domain-specific subnetwork reached significance in the deep group. In the shallow group and in the pooled sample, no channel was significantly correlated with behavior. These channel-level correlations are exploratory and uncorrected for multiple comparisons (**Figure 7**).

**Figure 7.**
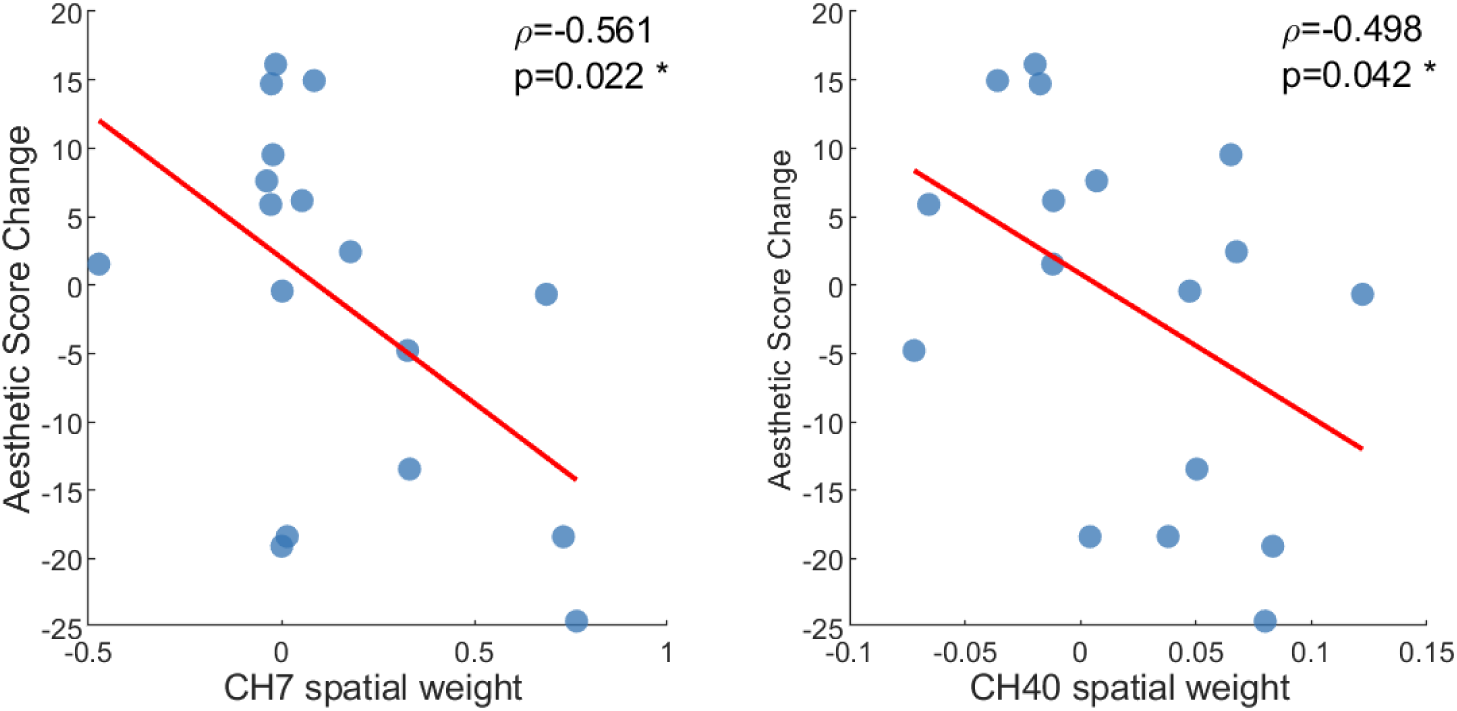
Channel-wise brain–behavior correlations of the component-1 topography (deep group). Signed, unit-normalized component-1 spatial weights (averaged over 0.050– 0.088 Hz) correlated with the aesthetic-rating change score. Two DMN channels showed significant negative associations — left LTC (ρ = −0.561, p = 0.022) and left inferior parietal lobule (ρ = −0.498, p = 0.042); right LTC showed a trend (ρ = −0.424, p = 0.093). No domain-specific channel, and no channel in the shallow group or the pooled sample, reached significance.

##### Subnetwork normalized entropy: group comparison

In the DMN, normalized Shannon entropy was higher in the deep group (0.807 ± 0.124) than in the shallow group (0.678 ± 0.184; permutation *p* = 0.022, Cohen’s *d* = 0.84; Mann– Whitney *U p* = 0.024; deep *n* = 18, shallow *n* = 15). The domain-specific subnetwork showed no group difference (deep 0.744 ± 0.133 vs shallow 0.777 ± 0.088; permutation *p* = 0.42, Cohen’s *d* = −0.29). Because both subnetwork entropies were derived from the same component, the same frequency band, and the same analysis pipeline, the selective DMN effect cannot be attributed to any factor of that pipeline acting uniformly across channels (**Figure 8**).

**Figure 8.**
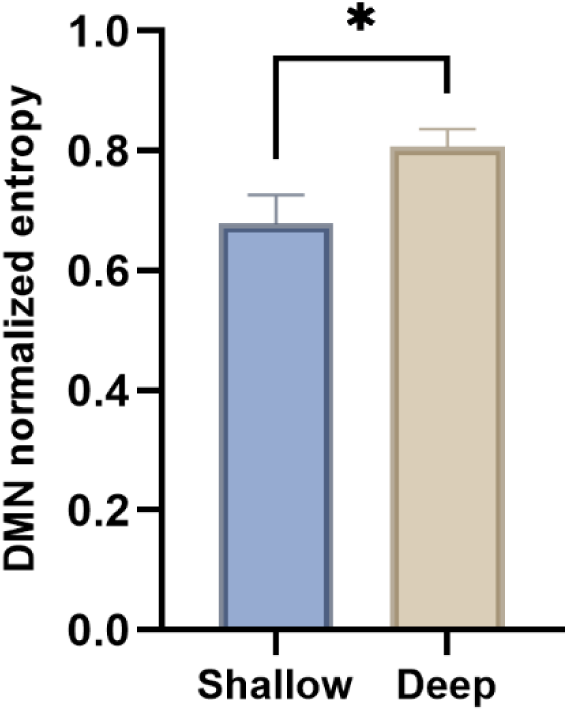
Subnetwork normalized entropy by group. Normalized Shannon entropy of the component-1 topography within the DMN and the domain-specific subnetwork for the deep and shallow groups. DMN entropy was higher in the deep group (0.807 ± 0.124 vs 0.678 ± 0.184; permutation *p* = 0.022, Cohen’s *d* = 0.84; Mann–Whitney *U p* = 0.024); the domain-specific subnetwork did not differ (0.744 ± 0.133 vs 0.777 ± 0.088; *p* = 0.42). Error bars denote SE.

##### Subnetwork entropy, energy, and behavior

###### Entropy × behavior

In the shallow group (*n* = 14), domain-specific subnetwork entropy was positively correlated with the aesthetic-change score (ρ = +0.609, *p* = 0.026). The DMN showed no such relationship in the shallow group, and neither subnetwork’s entropy was correlated with behavior in the deep group.

###### Energy × behavior

In the shallow group, DMN energy was positively correlated with the aesthetic-change score (ρ = +0.701, *p* = 0.007) and domain-specific energy was negatively correlated with it (ρ = −0.604, *p* = 0.023). In the deep group, neither DMN energy (ρ = −0.1, *p* = 0.72) nor domain-specific energy (ρ = +0.08, *p* = 0.77) was correlated with behavior.

###### Moderation by group

For network energy, the moderation model (behavior ∼ group + energy + group × energy) showed an interaction at the trend level (DMN: interaction *p* = 0.085; difference in group-wise slopes, permutation *p* = 0.097; deep slope ≈ 0, shallow slope positive). The domain-specific interaction was not significant (*p* = 0.33). The present evidence therefore establishes a set of mutually consistent network–behavior associations in the shallow group and their absence in the deep group, but does not by itself establish a statistically confirmed difference in coupling strength between groups (**Figure 9**).

**Figure 9.**
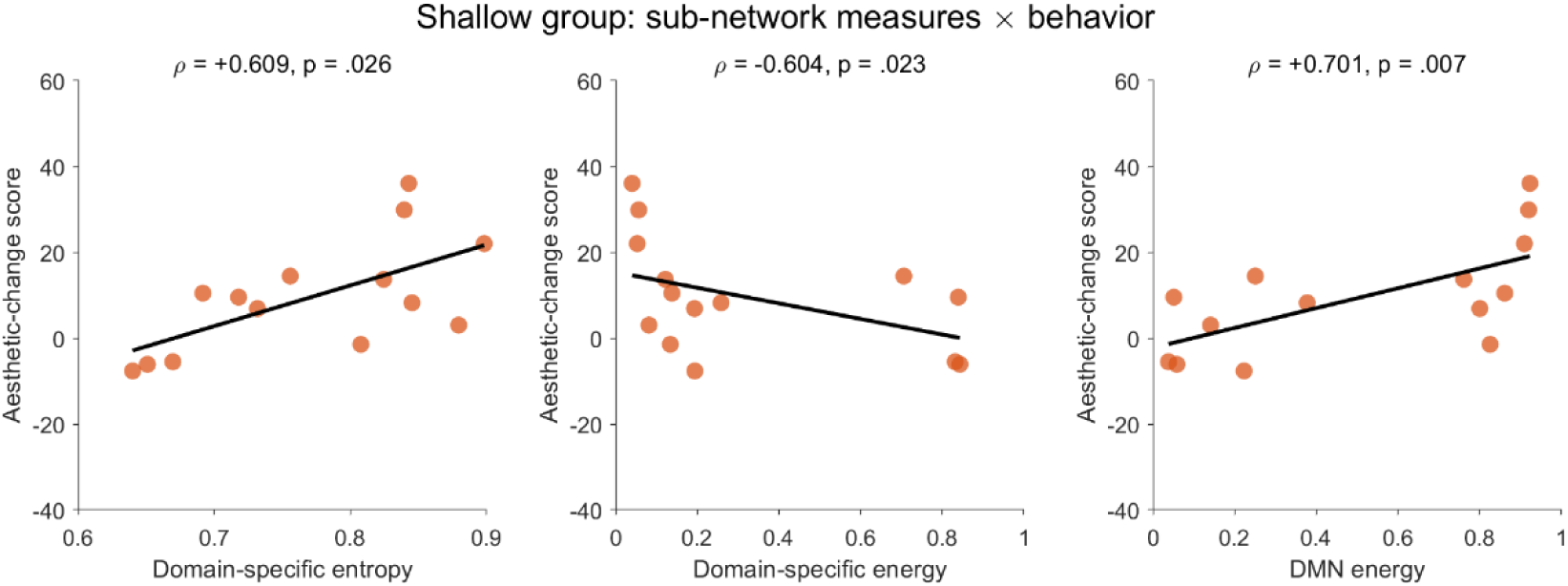
Exploratory associations of subnetwork entropy and energy with aesthetic-rating change. In the Shallow Analysis group, domain-specific subnetwork entropy was positively associated with aesthetic-rating change, DMN energy was positively associated with aesthetic-rating change, and domain-specific subnetwork energy was negatively associated with aesthetic-rating change. Corresponding associations were not statistically significant in the Deep Analysis group. Group differences in correlation strength did not reach statistical significance.

## 4. Discussion

The present study examined whether the depth of analysis affects aesthetic experience, and whether any such effect is confined to particular categories of aesthetic object and particular dimensions of aesthetic response. Deeper analysis proved to have no uniform direction. For visual art, it was costly on the affective dimension alone, removing the gain that shallow analysis produced and returning ratings to the image baseline; for natural landscapes, it instead enlarged the gain on all three dimensions. Experiment 2 replicated the target effect in an independent sample and found it accompanied by a more prominent leading task component at 0.050–0.088 Hz, by higher spatial entropy within the default mode network (DMN) but not within the domain-specific prefrontal subnetwork, and by the absence of the network–behavior associations present under shallow analysis.

### 4.1 Deeper analysis removes the gain from analysis, not aesthetic experience itself

Interpreting these results requires first establishing what deeper analysis actually changed. Titles, contextual information, and viewing guidance help viewers attend to, organize, and comprehend the information available in a work, thereby raising aesthetic evaluation (Millis, 2001; Russell, 2003; Leder et al., 2006). The present findings do not contradict this: shallow analysis raised affective ratings of visual art above the image baseline in both experiments, with near-identical standardized effect sizes.

What changed was the depth at which the analysis was pitched. When affective evaluation moved from a single global judgment to the constituent attributes of pleasure, impact, relaxation, and harmony, ratings returned to baseline rather than falling below it — a conclusion supported by equivalence and Bayesian tests in both samples, and so resting on positive evidence rather than on a null result alone. Deeper analysis therefore did not degrade the aesthetic experience viewers already had; it dissolved the increment that shallow analysis added. What requires explanation is the disappearance of an analytic benefit, not an impairment of experience.

This rules out any account on which analysis is detrimental as such — for instance, that decomposing a judgment into attributes shifts its basis toward information that is easy to verbalize (Wilson & Schooler, 1991). Such accounts over-predict the present data: if deeper analysis itself was the source of the cost, the same attenuation should have appeared for natural landscapes, where the opposite obtained in all three dimensions. Whether deeper analysis exacts a cost thus depends on what is being analyzed more deeply — and the neural findings of Experiment 2 exclude the remaining possibility that the deep group simply invested less (Section 4.3).

### 4.2 Conditional specificity: dimensions, categories, and the functional heterogeneity of category-specific systems

Three connected features of the behavioral pattern bear on this conditionality.

First, the three aesthetic dimensions are not each proprietary to one class of object. For natural landscapes, deeper analysis of the perceptual, semantic, and affective dimensions all yielded larger gains, supporting the multidimensional structure of aesthetic experience, in which evaluation is jointly constituted by sensory–perceptual, affective–evaluative, and semantic– knowledge components (Leder et al., 2004; Marković, 2012; Chatterjee & Vartanian, 2014; Pelowski et al., 2017; Lin et al., 2026). This establishes a negative premise: analyzing a dimension more deeply does not in itself interrupt the experience as a whole.

Second, that all three dimensions can participate does not entail that categories are equally sensitive to them. Visual art and natural landscapes showed different dimension-by-depth correlation, consistent with category-specific neural representation. Vessel et al. (2019) found that category-specific aesthetic information for natural scenes is carried principally along the ventral visual pathway, that aesthetic evaluation of artworks additionally recruits frontopolar, orbitofrontal, and inferior frontal regions, and that the DMN represents aesthetic appeal across visual categories. Liang et al. (2026) subsequently isolated two principal dimensions of aesthetic evaluation — visual–semantic and hedonic — tracked respectively by category-selective ventral visual regions and by medial prefrontal and subcortical circuitry. The difference between image categories here is therefore better read as one of relative reliance than as a claim that art is affective and landscape perceptual.

Third, and critically: the direction of the depth effect reversed across the two putatively matched conditions, indicating that category specificity does not entail functional homogeneity. Were a category-specific system merely the location at which information about that category is carried, analyzing its preferred dimension more deeply should produce effects of the same sign for both image categories. Ventral occipitotemporal cortex principally represents color, shape, spatial layout, and scene structure — information extractable directly from the stimulus (Vessel et al., 2019; Isik & Vessel, 2021) — and along this pathway, aesthetic value is built hierarchically through weighted integration of low- and high-level features (Iigaya et al., 2023). Drawing finer distinctions among the structure, color, spatiality, and complexity of a landscape raises the resolution of content representation, and that finer componential information remains available to global evaluation. The prefrontal system associated with artworks, by contrast, links visual information to affect, meaning, and evaluation (Chatterjee & Vartanian, 2014; Vessel et al., 2019; Liang et al., 2026). The affect an artwork evokes is not the sum of pleasure, impact, relaxation, and harmony; it depends on the relations these bear to one another within the viewer.

We therefore advance a testable proposition: the benefit of deeper analysis depends on the representational form on which the analyzed dimension rests within that category. Where the representation is content-additive — where a global evaluation can be assembled from finer componential information — deeper analysis raises resolution, and the added information continues to feed evaluation. Where it is relationally integrated — where global value derives from the relations among components rather than from the components themselves — deeper analysis rewrites those relations as a set of juxtaposed attributes, altering the organization of the global representation rather than its resolution, and the benefit disappears. The cost of deeper analysis arises not from processing too much, but from processing in the wrong form.

So construed, verbal-overshadowing and fluency effects are neither general laws nor counterexamples but boundary cases. Attribute-wise rating increases the explicitness of specific information and may thereby reweight the sources entering a global judgment (Wilson & Schooler, 1991; Schooler & Engstler-Schooler, 1990) — but only where that judgment is relational in character. Likewise, moderate analysis can facilitate aesthetic experience by organizing information and reducing uncertainty (Reber et al., 2004; Van de Cruys & Wagemans, 2011; Muth & Carbon, 2013), yet analysis extended into multiple specific attributes requires the viewer to reconcile a global impression against its subdimensions, which need not increase fluency and may instead surface new discrepancies. These are the specific costs incurred when a relationally integrated representation is decomposed.

### 4.3 The neural level: not less task processing, but redistribution within the DMN

The neural findings first exclude a parsimonious alternative. After removal of the components shared with passive viewing and with affective processing, the residual leading task component at 0.050–0.088 Hz was more prominent in the deep group, with a large effect size. Task-relevant neural organization was therefore stronger, not weaker, under deeper analysis, and the smaller aesthetic gain cannot be attributed to insufficient engagement. The intensity of aesthetic experience is not predictable from the quantity of task-related processing.

The operational load attaching to depth of analysis was moreover removed by design rather than by argument. The task network was estimated not from the task state directly but from the residual remaining after removal, within each participant and at each frequency, of the joint span of two baseline subspaces: the image baseline, and an affective-analysis baseline in which participants performed exactly the same analytic procedure on abstract graphics containing no aesthetic content — the deep group again rating four affective components, the shallow group again making a single global judgment — estimated separately for each group. Differences in the number of judgments, task switching, and working-memory load were thereby projected out. Group differences in the residual network are attributable only to what an analysis of equivalent depth does when applied to stimuli carrying aesthetic content. More broadly, recent work on non-equilibrium brain dynamics suggests that subjective states cannot be characterized by the magnitude of neural activity alone, but also depend on how spontaneous fluctuations and perturbation-evoked responses are dynamically organized (Berjaga-Buisan et al., 2026). Although the FDT-violation measure used in that study differs from the present measure of spatial entropy, the distinction is relevant here: deeper analysis elicited a more prominent task-related component, yet this stronger network expression did not translate into a comparable aesthetic gain. This raises the possibility that the subjective consequence of analysis depends not only on how much task-related processing is recruited, but also on how that processing is organized within the relevant networks.

Deeper analysis selectively raised the spatial entropy of the DMN subnetwork, while the entropy of the domain-specific subnetwork and the group-level similarity of the overall component topography did not differ. The evidence therefore supports spatial redistribution within the DMN specifically, rather than reorganization of the task network as a whole.

Notably, the domain-specific subnetwork here — bilateral IFG, orbitofrontal cortex, and frontal pole — is precisely the prefrontal system Vessel et al. (2019) associated with aesthetic evaluation of artworks: its spatial organization was unchanged while the DMN’s was not. This converges with the proposition of Section 4.2, that what deeper analysis alters is not content representation but the manner in which content is organized into global aesthetic value. It also serves as an internal control, since both entropies derive from the same component, frequency band, and pipeline, so the selective DMN effect cannot reflect any feature of that pipeline acting uniformly across channels.

The relationship between network configuration and behavior points in the same direction. In the shallow group, the aesthetic increase was greater when the task network was relatively biased toward the DMN, and greater when content representation was distributed across more nodes of the domain-specific network — a pattern matching the model of Vessel et al. (2019), on which category-specific systems supply visual content while the DMN forms a representation of aesthetic appeal generalizing across categories. No such relation held in the deep group. The energy × group interaction reached only a trend level, however, so the evidence establishes a set of mutually consistent associations in the shallow group and their absence in the deep group, rather than a confirmed difference in coupling strength. An exploratory channel-level analysis was compatible with this: within the deep group, higher weights of left LTC and left inferior parietal lobule were associated with smaller aesthetic gains, though uncorrected and unreplicated in the shallow or pooled samples.

The loss of aesthetic benefit under affective analysis of artworks is therefore unlikely to reflect insufficient processing. Instead, the more finely specified affective information may have failed to be organized by the DMN in a way that supports a global aesthetic experience.

## 5. Conclusion

The present study shows that the facilitative effect of analysis on aesthetic experience has a boundary in depth. Decomposing affective evaluation from a global judgment into subordinate attributes did not impair aesthetic experience itself; it removed the gain that analysis would otherwise have conferred. This cost was conditionally specific, arising only on the affective dimension of visual art, whereas deeper analysis of all three dimensions of natural landscapes instead produced larger gains. We propose that the dissociation turns on the representational form on which the analyzed dimension rests within a given stimulus category: deeper analysis of a content-additive representation raises resolution and continues to feed global evaluation, whereas deeper analysis of a relationally integrated representation alters how the whole is organized. The neural evidence accords with this. Fine-grained analysis did not reduce task-related processing but selectively altered the distribution of spatial contribution within the DMN, accompanied by a weakening of its correspondence with subjective aesthetic experience. These findings suggest that the DMN’s role in aesthetic experience lies not in how much stimulus information it represents, but in whether that information can be organized into a global representation bearing subjective value.

