## Supplementary figures and images for "When Deeper Analysis Weakens Aesthetic Experience: Behavioral and Brain Network Evidence"

### 6d1f1a1c-e0ce-46de-b969-5bff83633c1b_2344.jpg

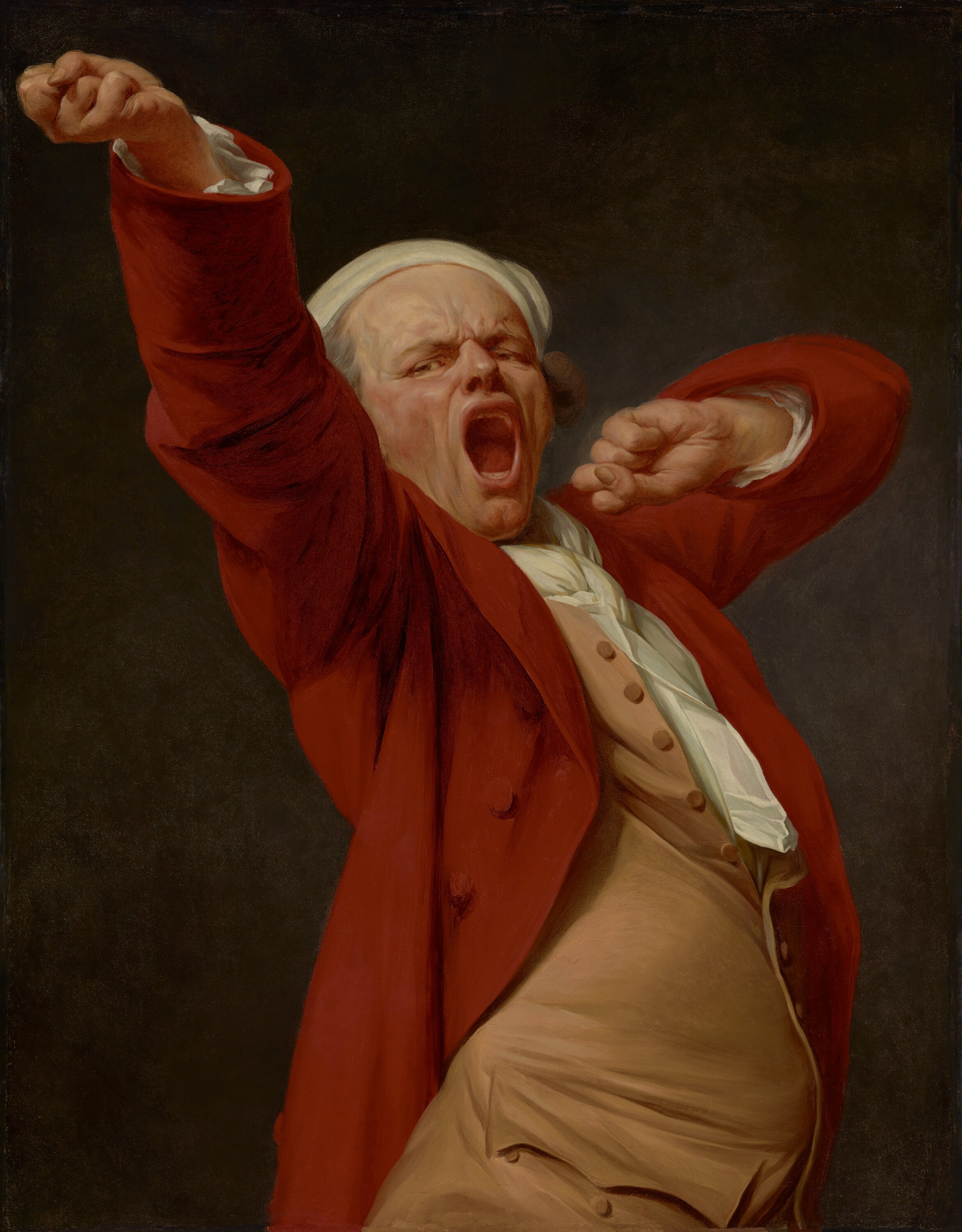

### 72c39c640475cbb086059dabbe49dbfe.png

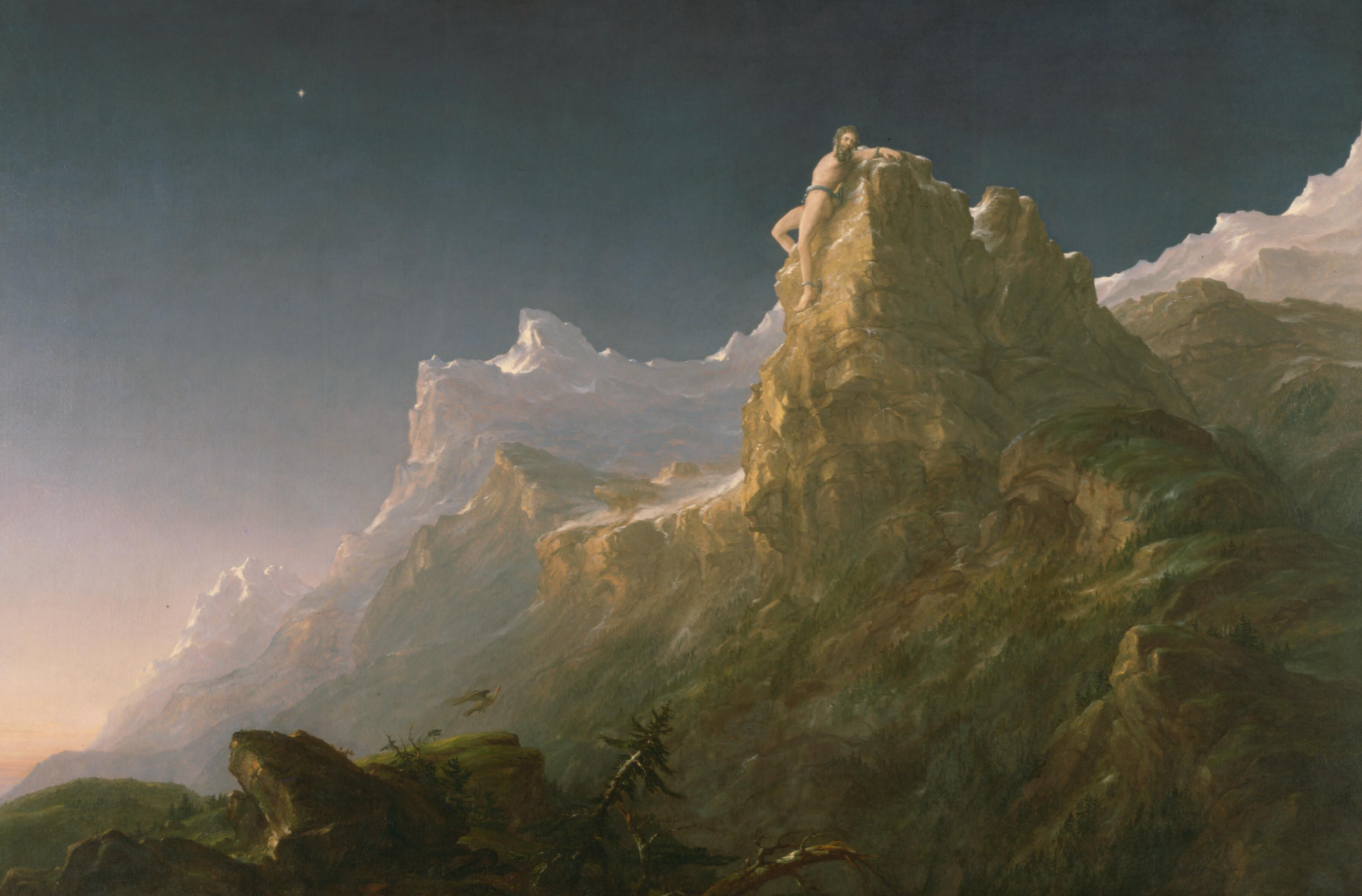

### 1002.jpg

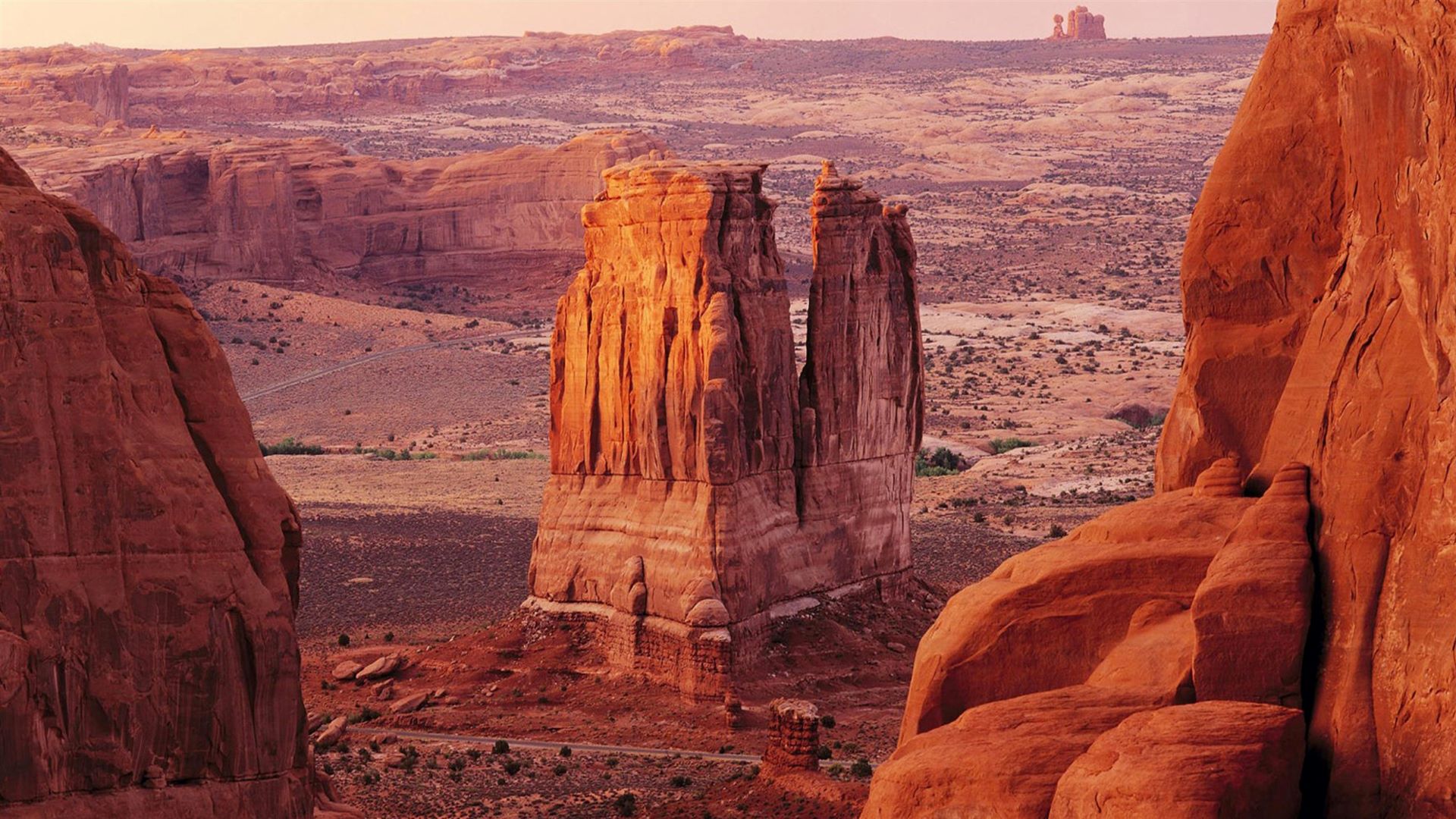

### 1004.jpg

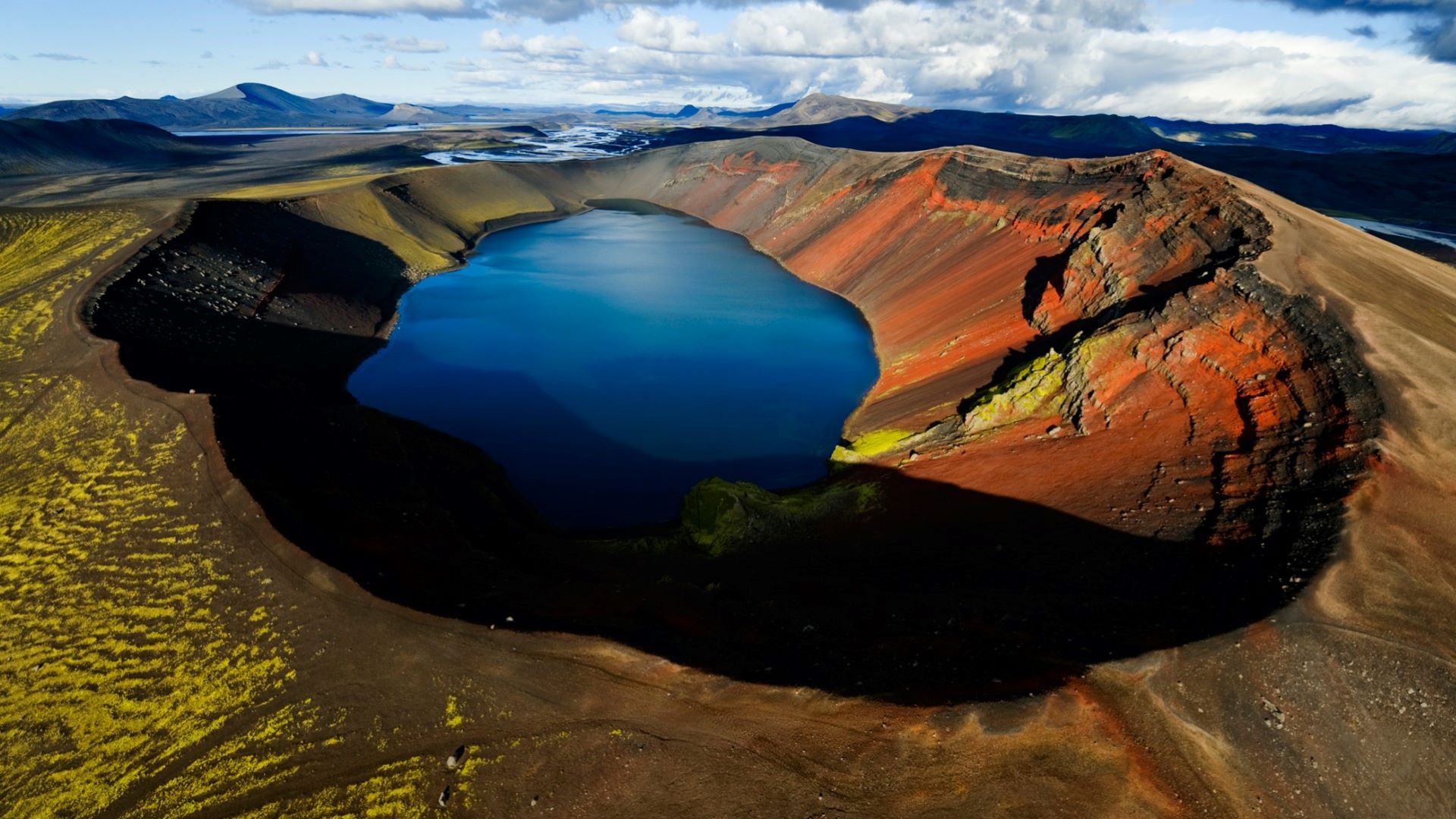

### 1006.jpg

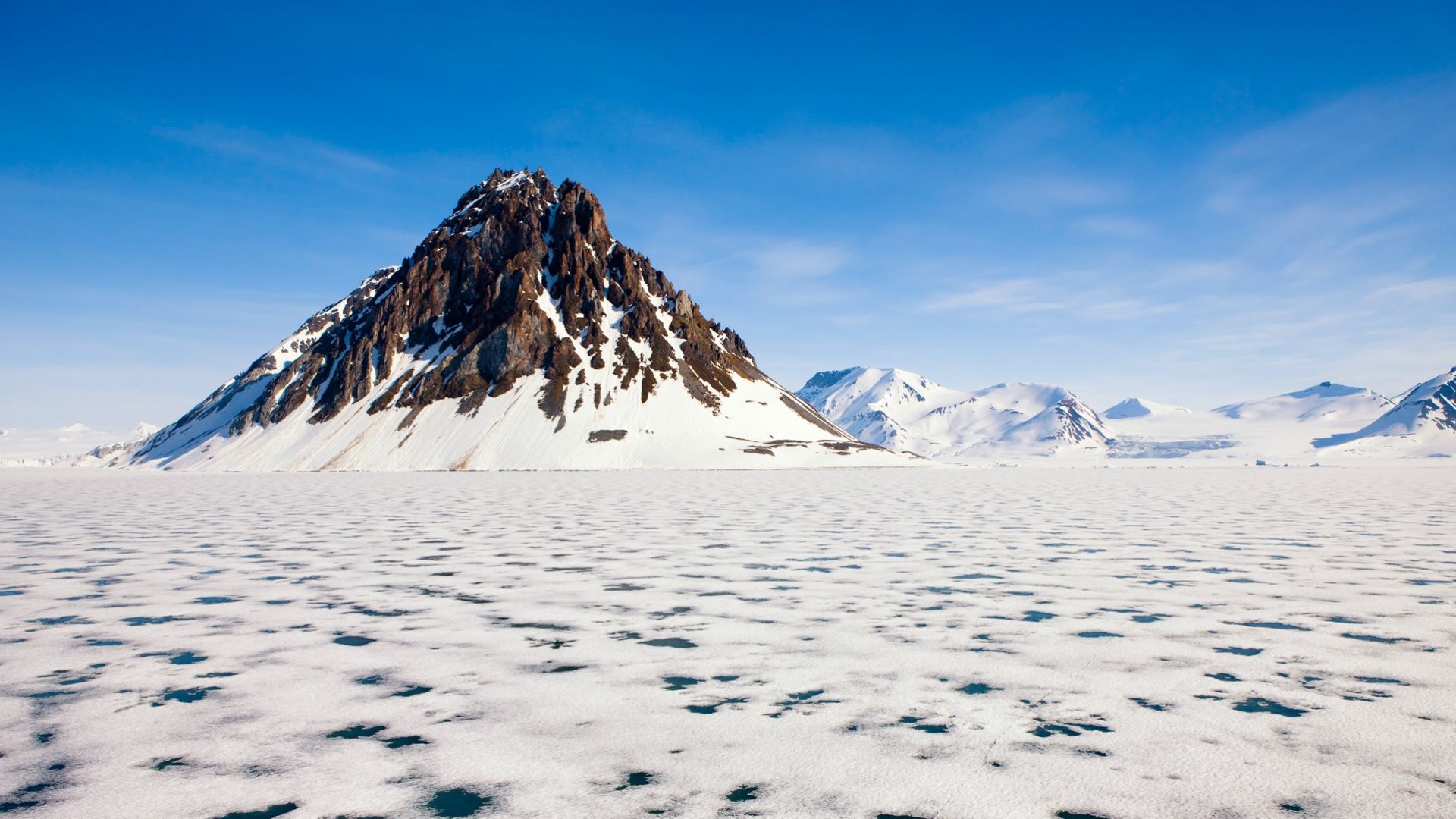

### 1024.jpg

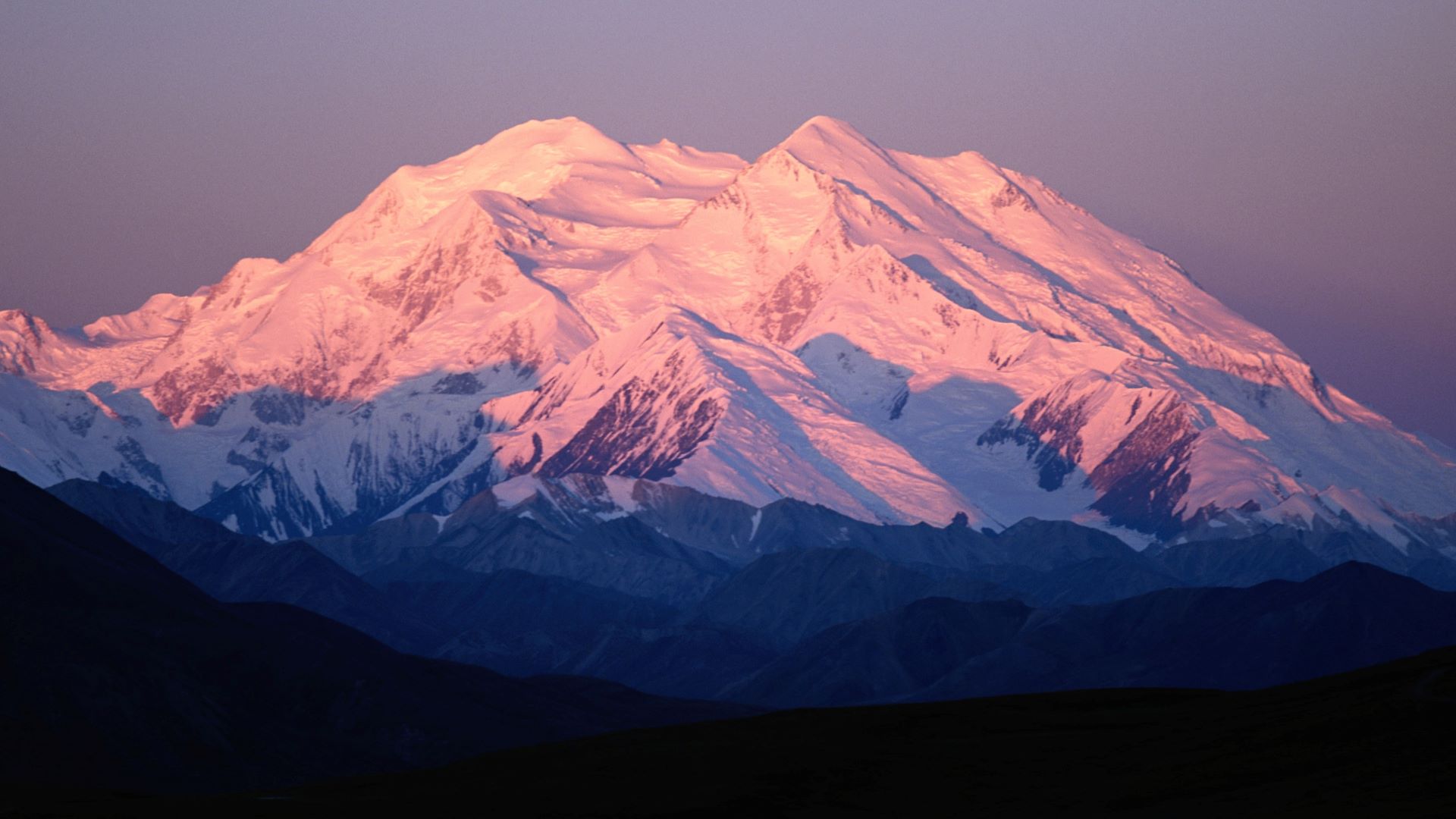

### 1034.jpg

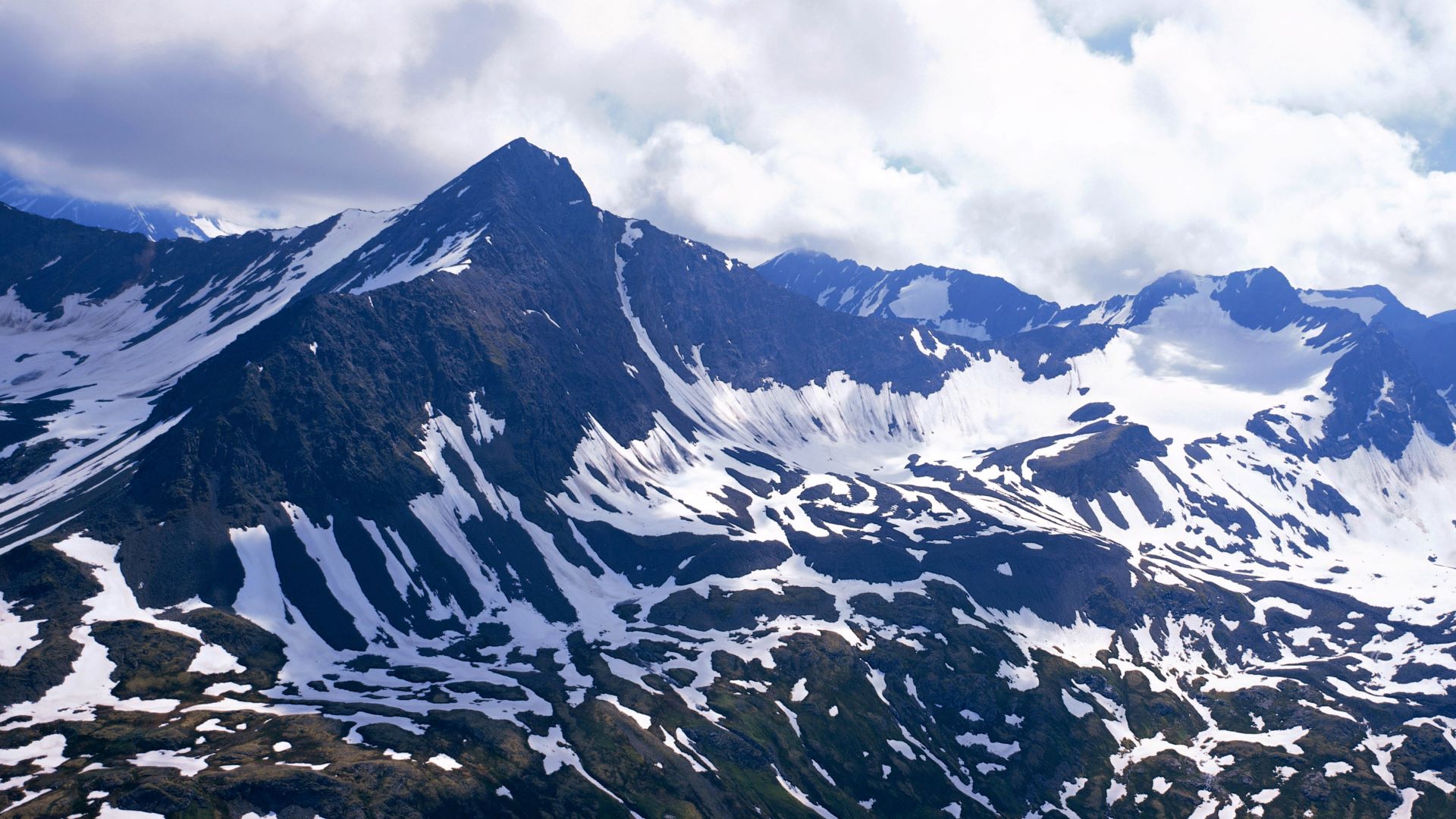

### 1046.jpg

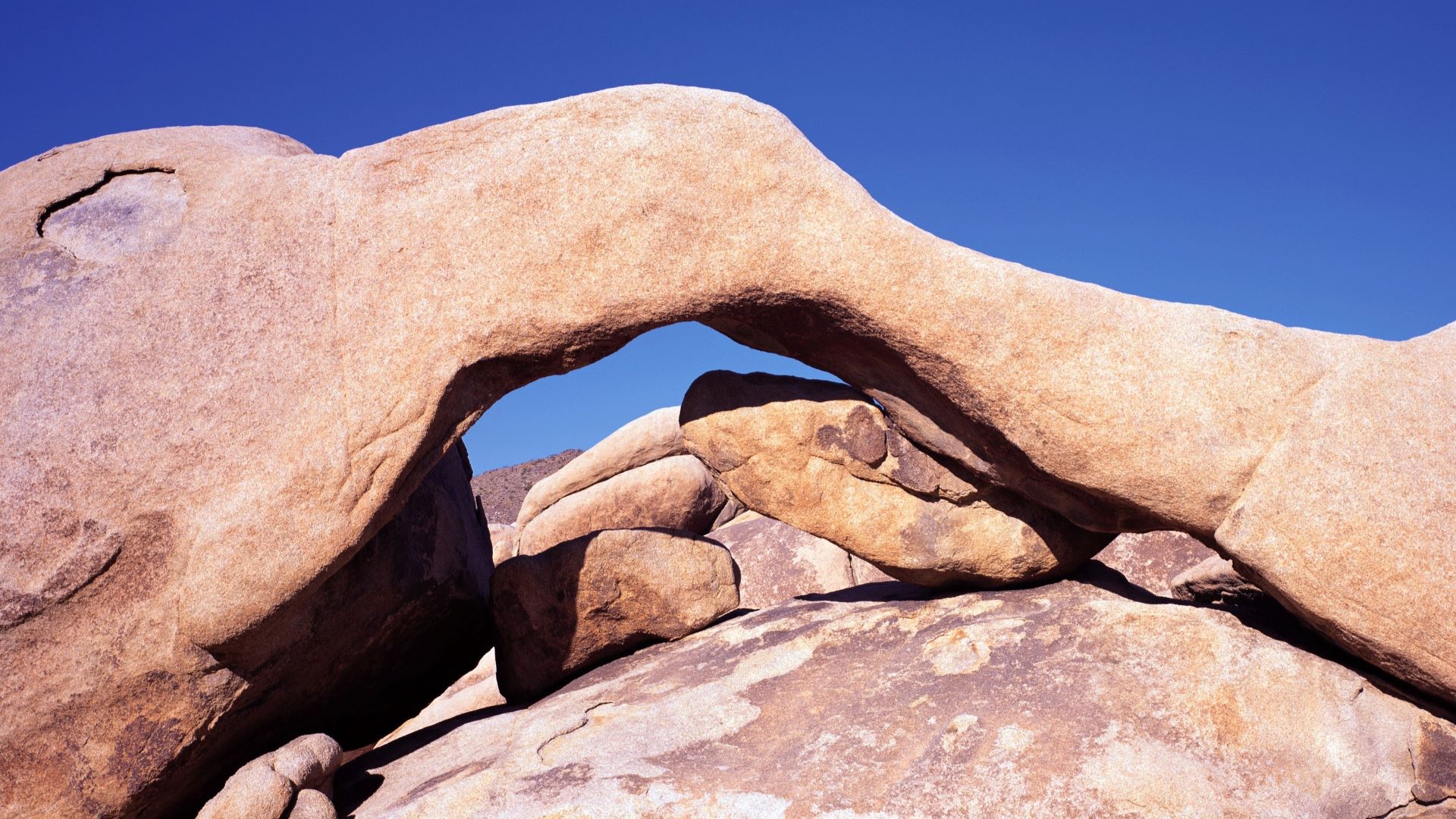

### 1060.jpg

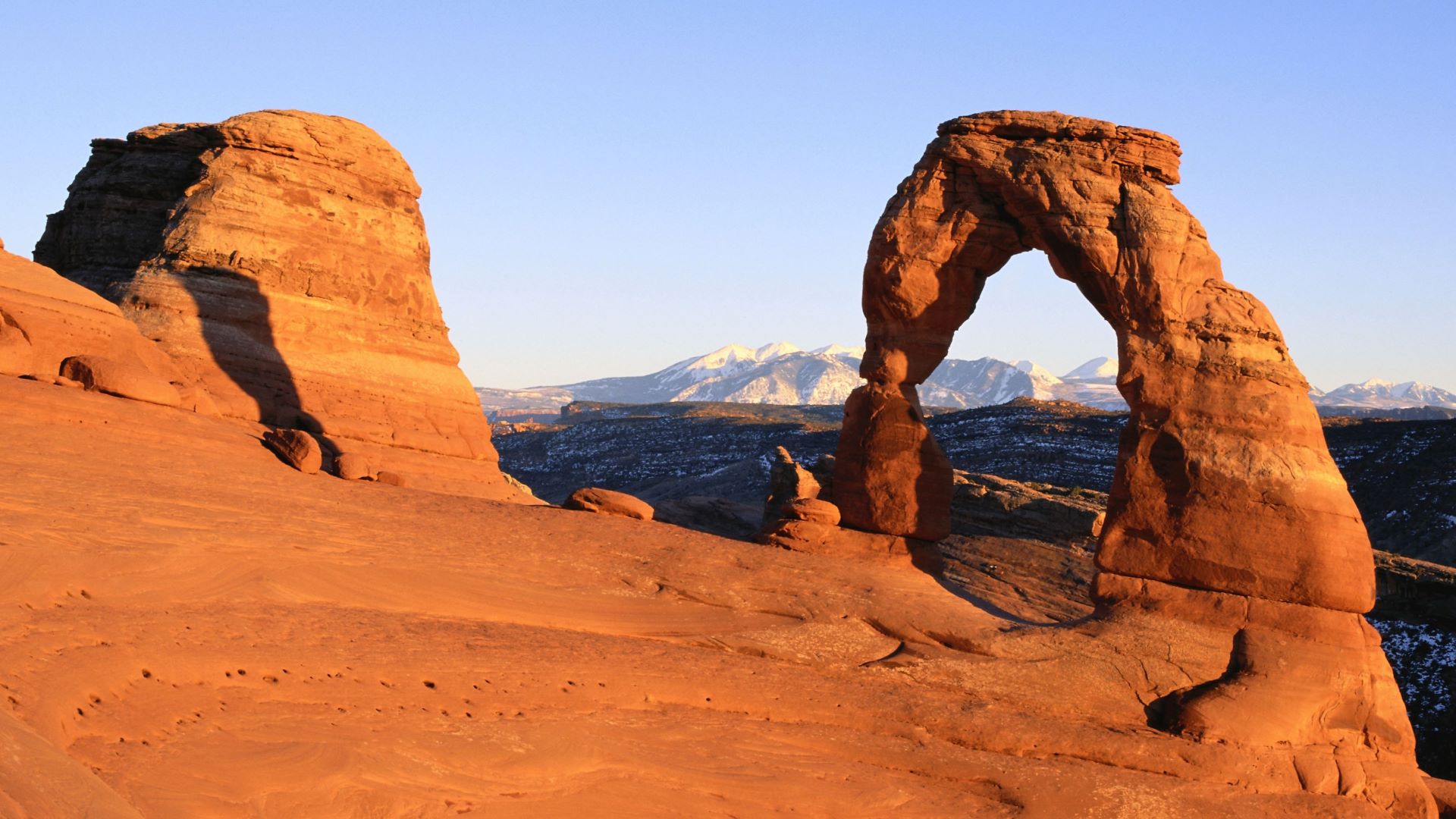

### 1067.jpg

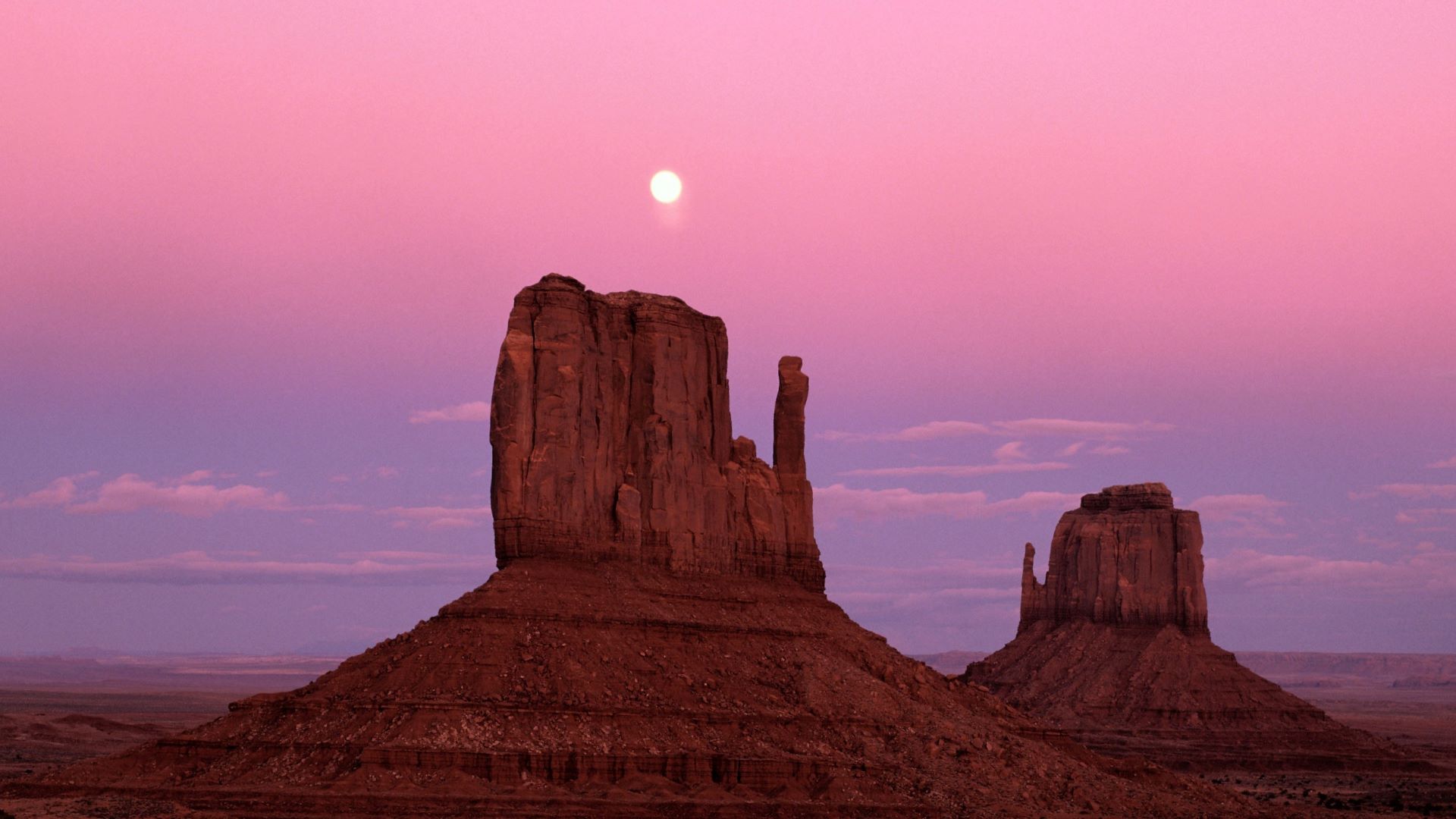

### 1069.jpg

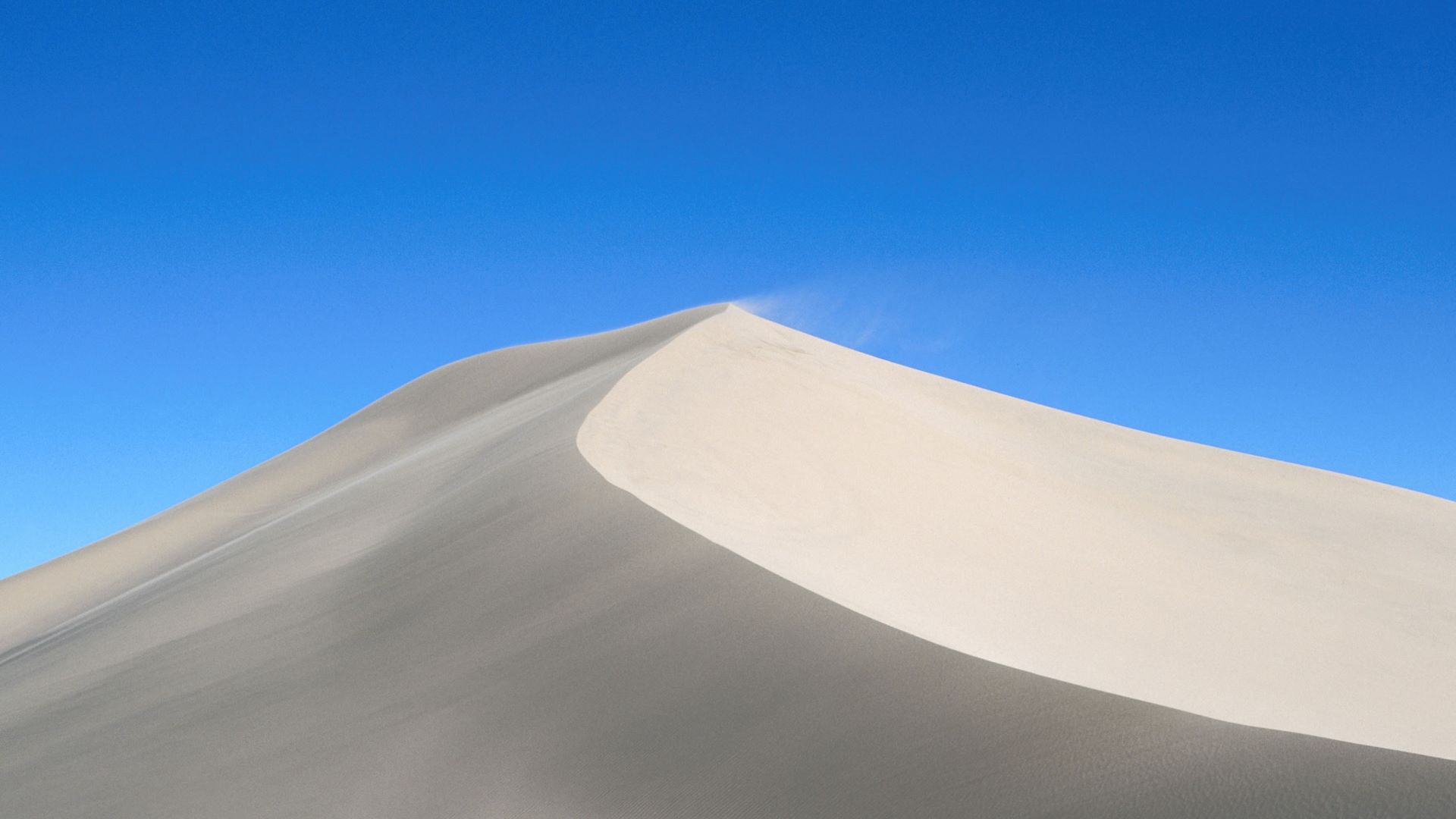

### 1071.jpg

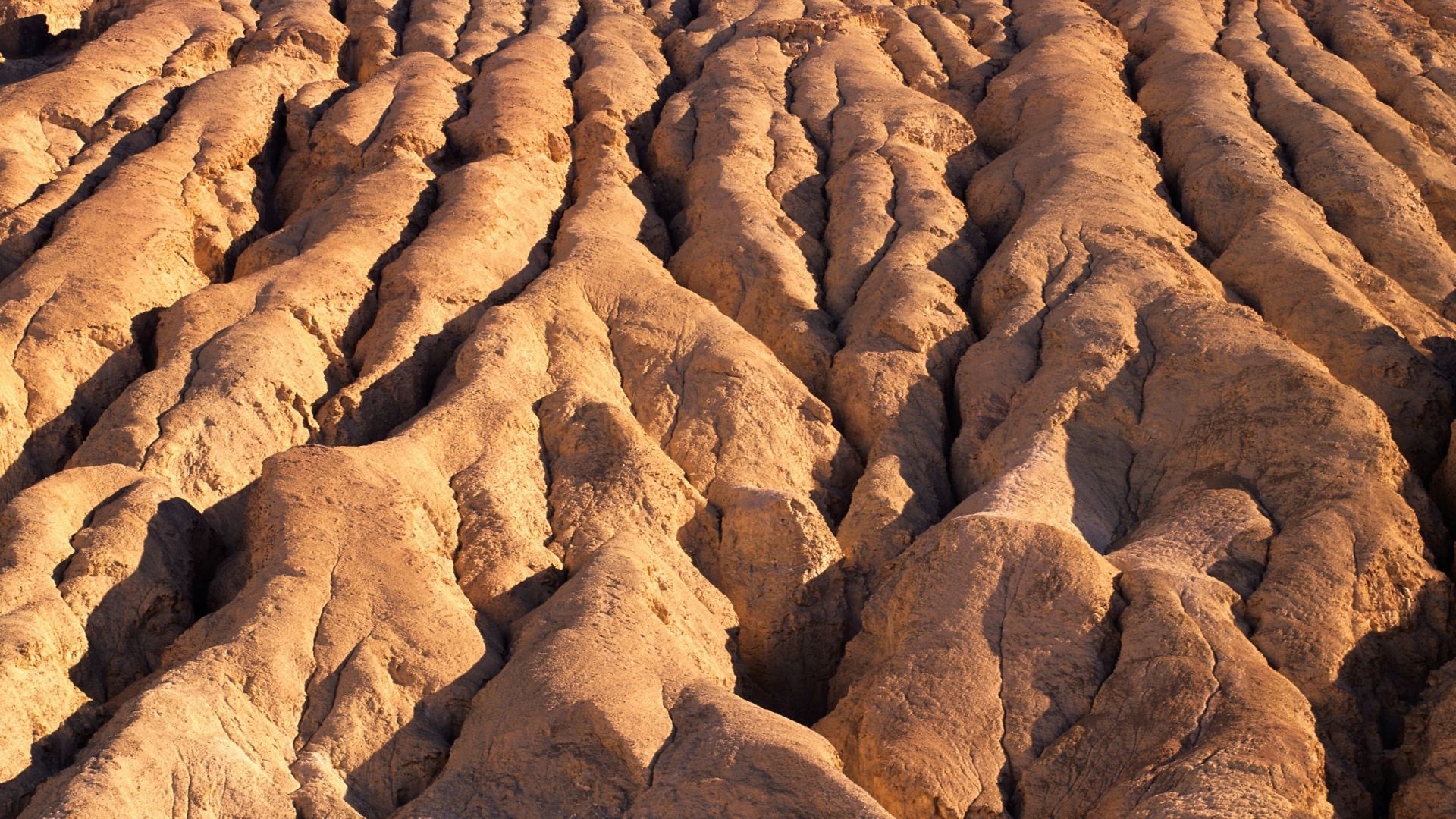

### 1079.jpg

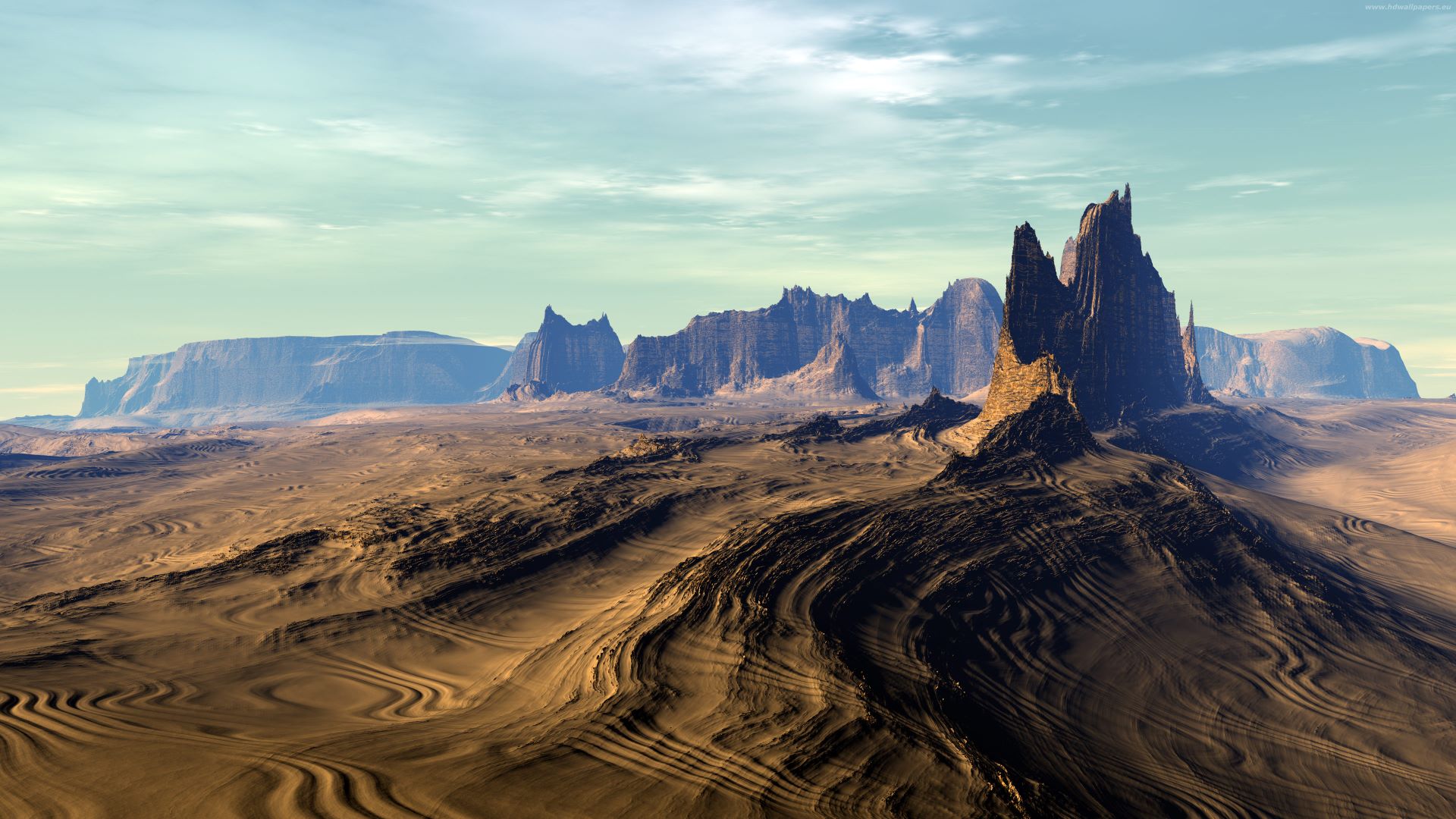

### 1081.jpg

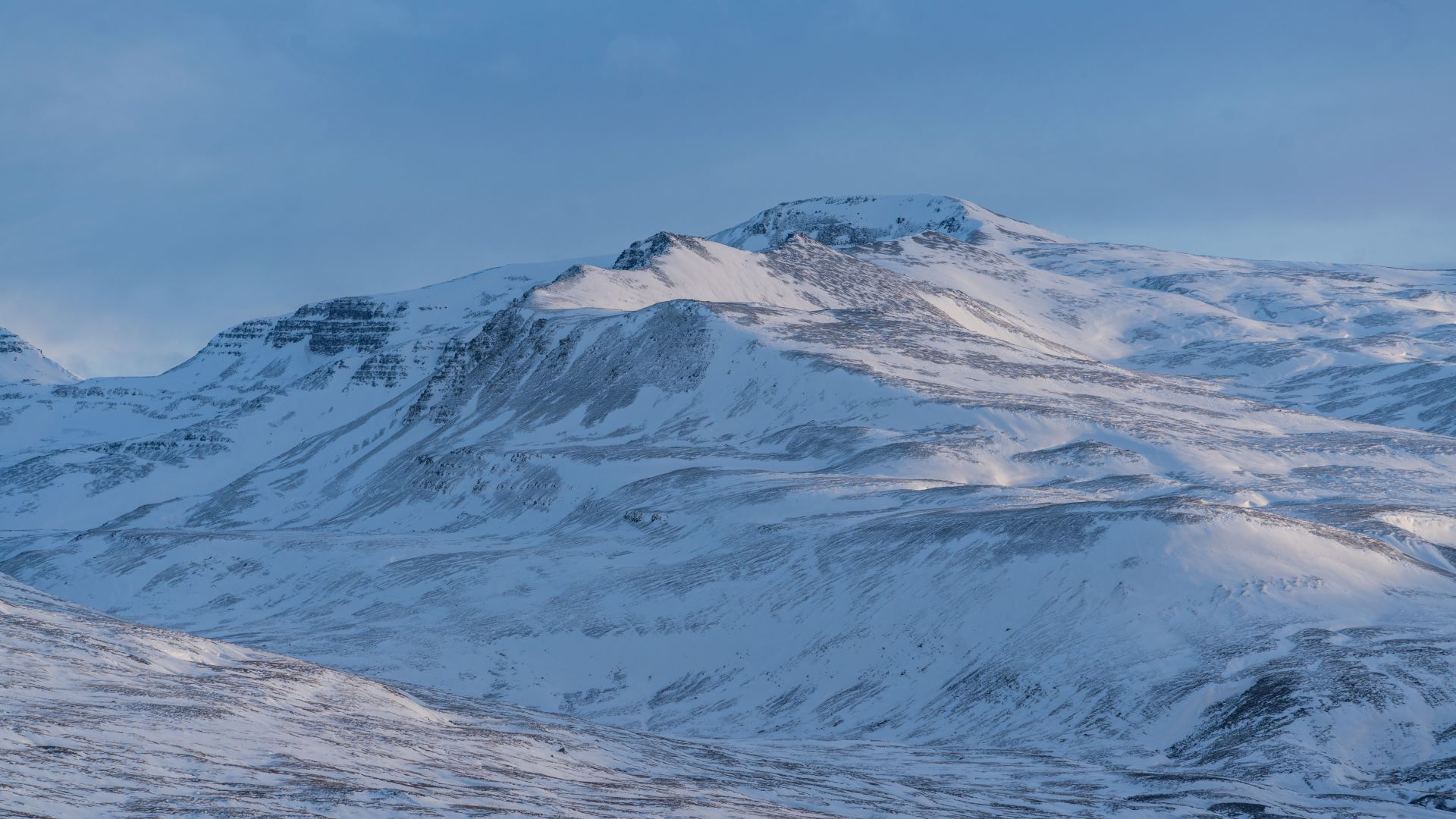

### 1083.jpg

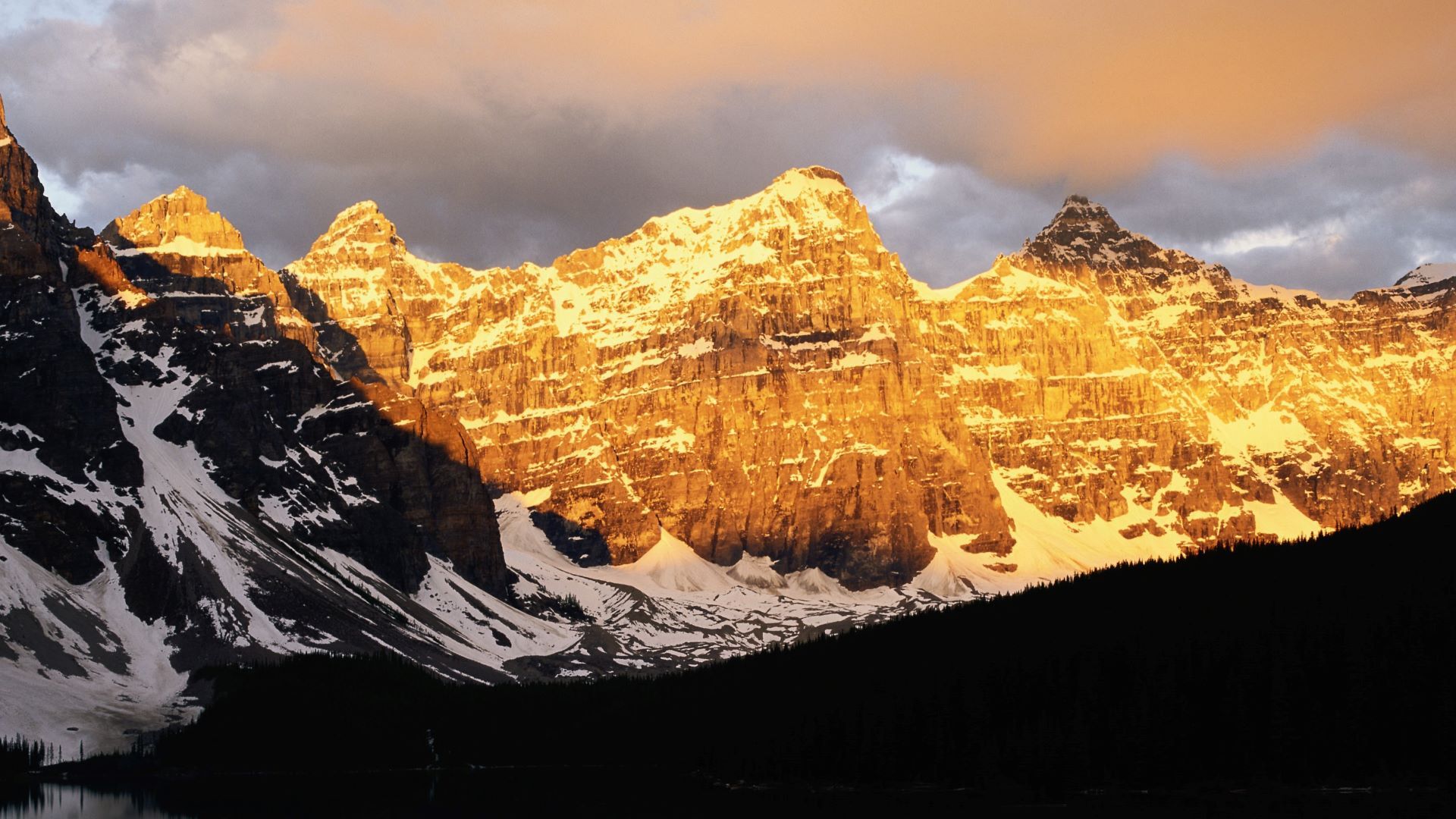

### 1949.541_print.jpg

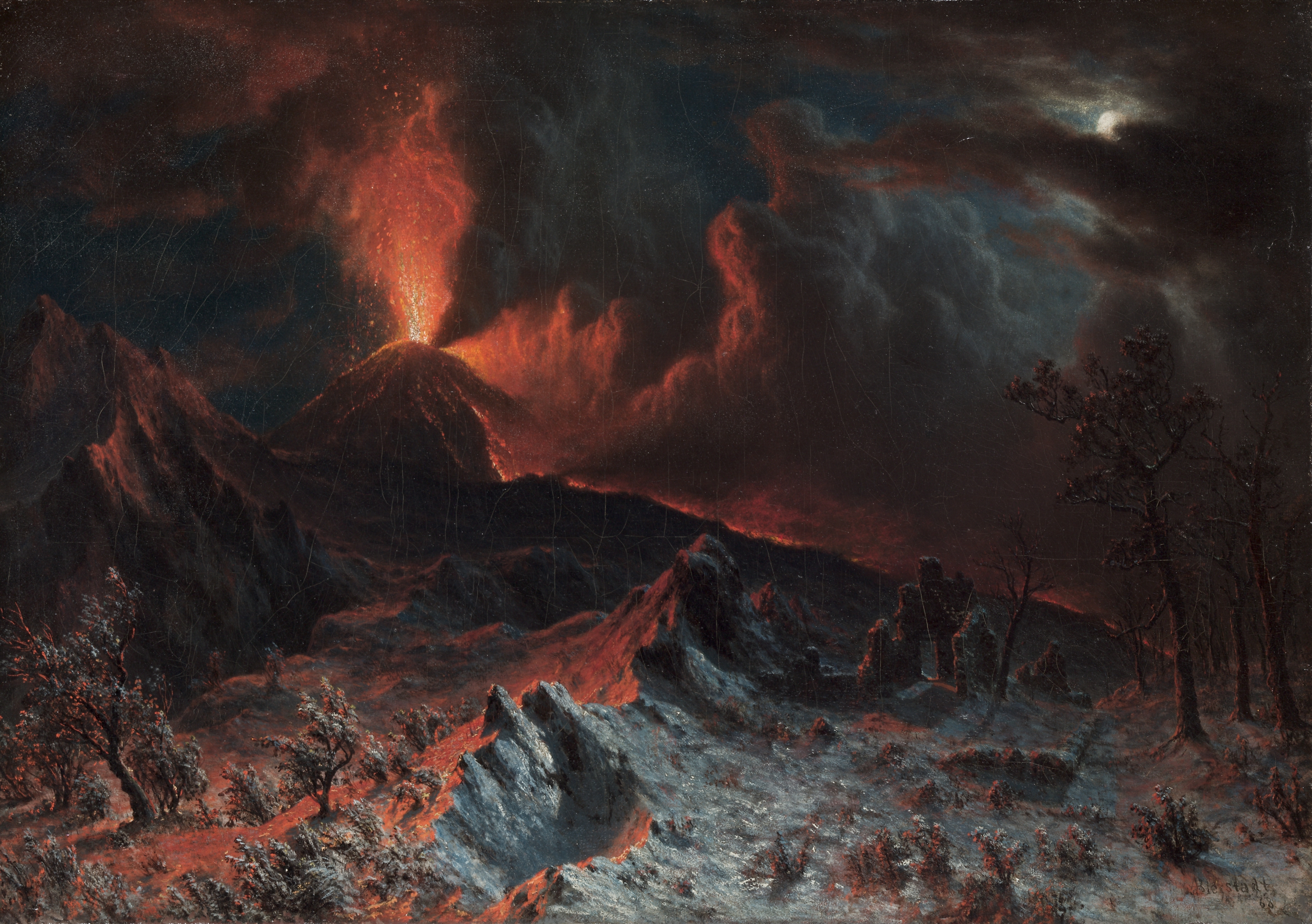

### 1949.585 - Trompe-l'Oeil Still Life with a Flower Garland and....jpg

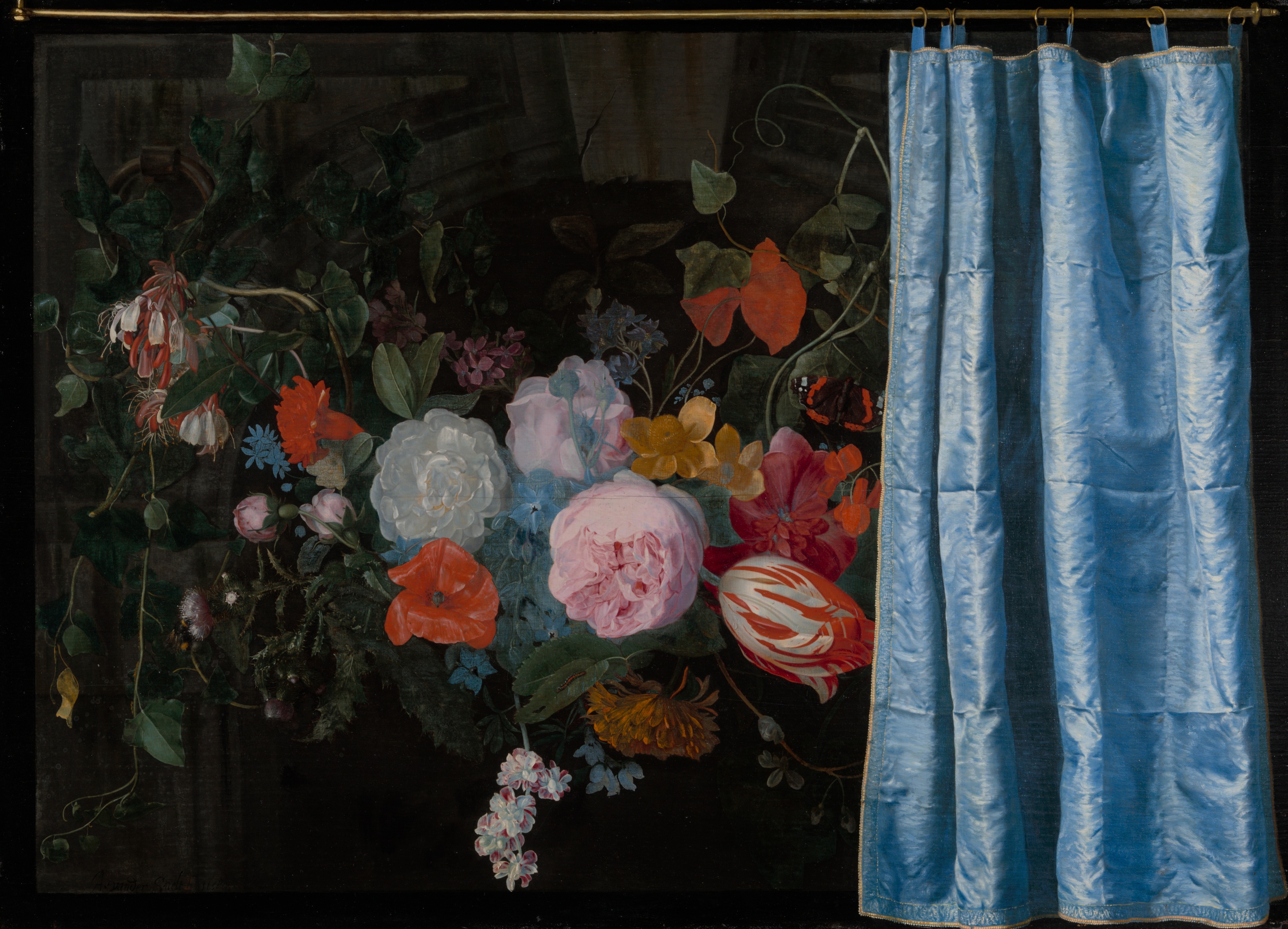

### 1952.235.3_print.jpg

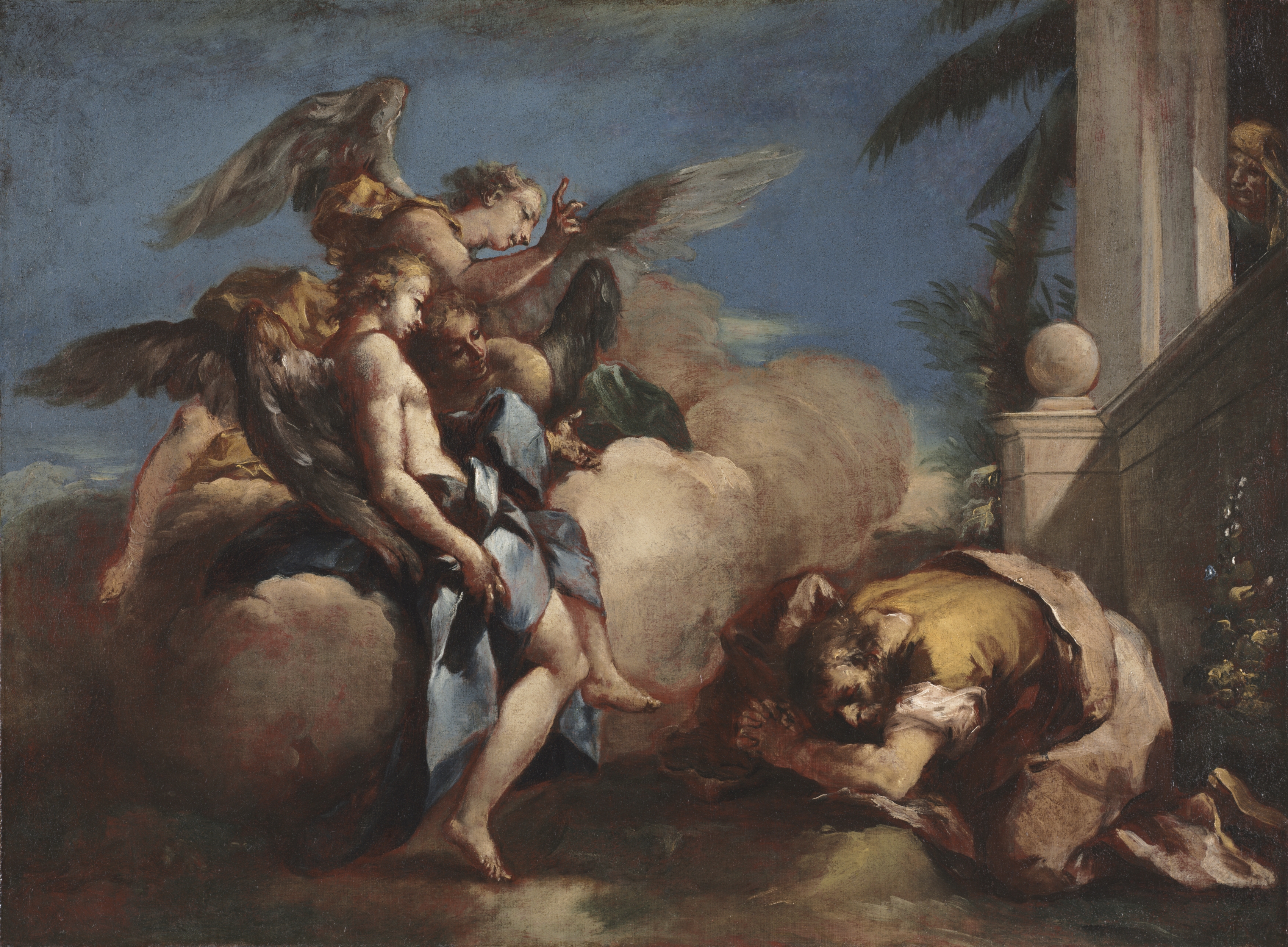

### 1952.542_print.jpg

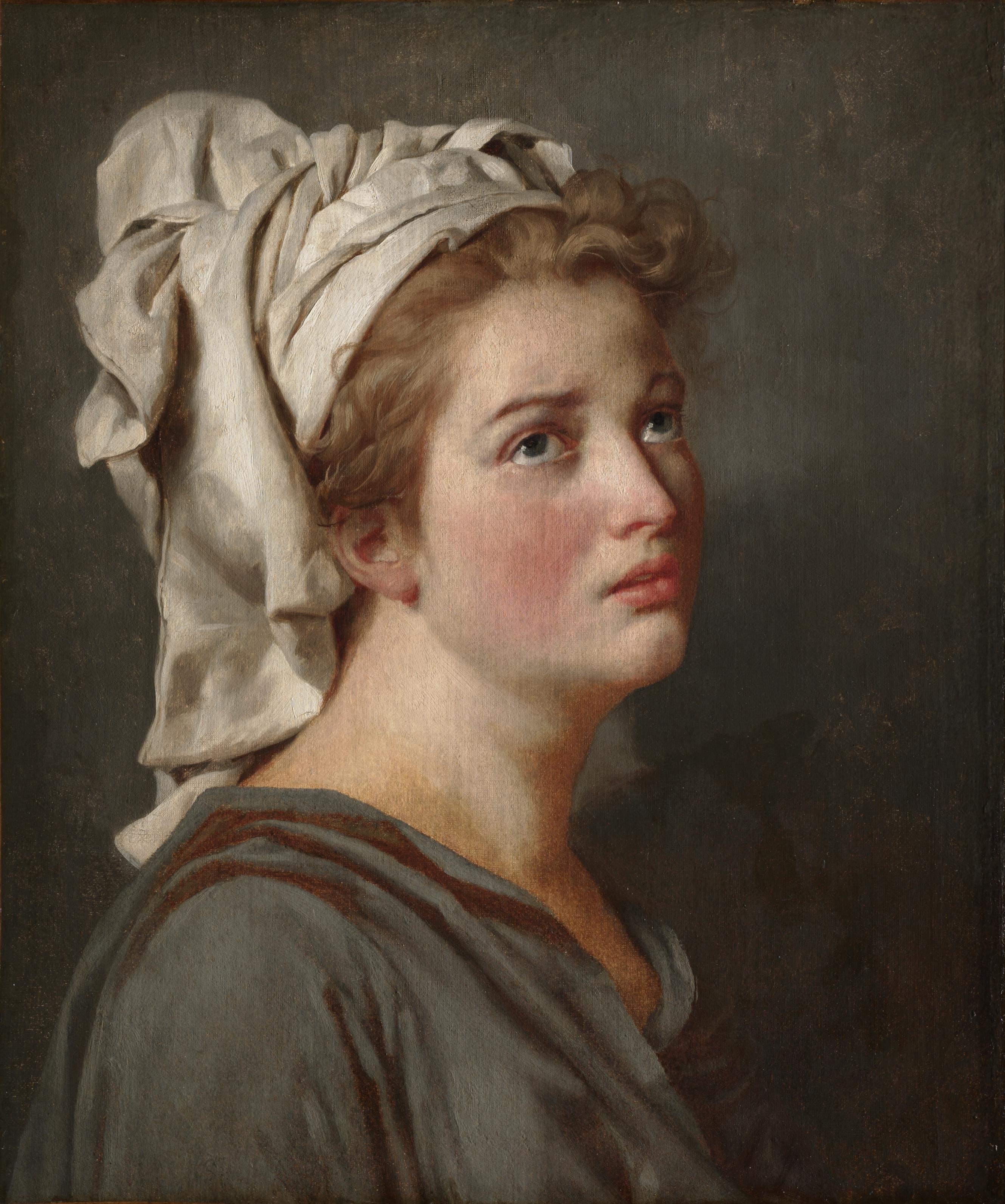

### 1955.865.jpg

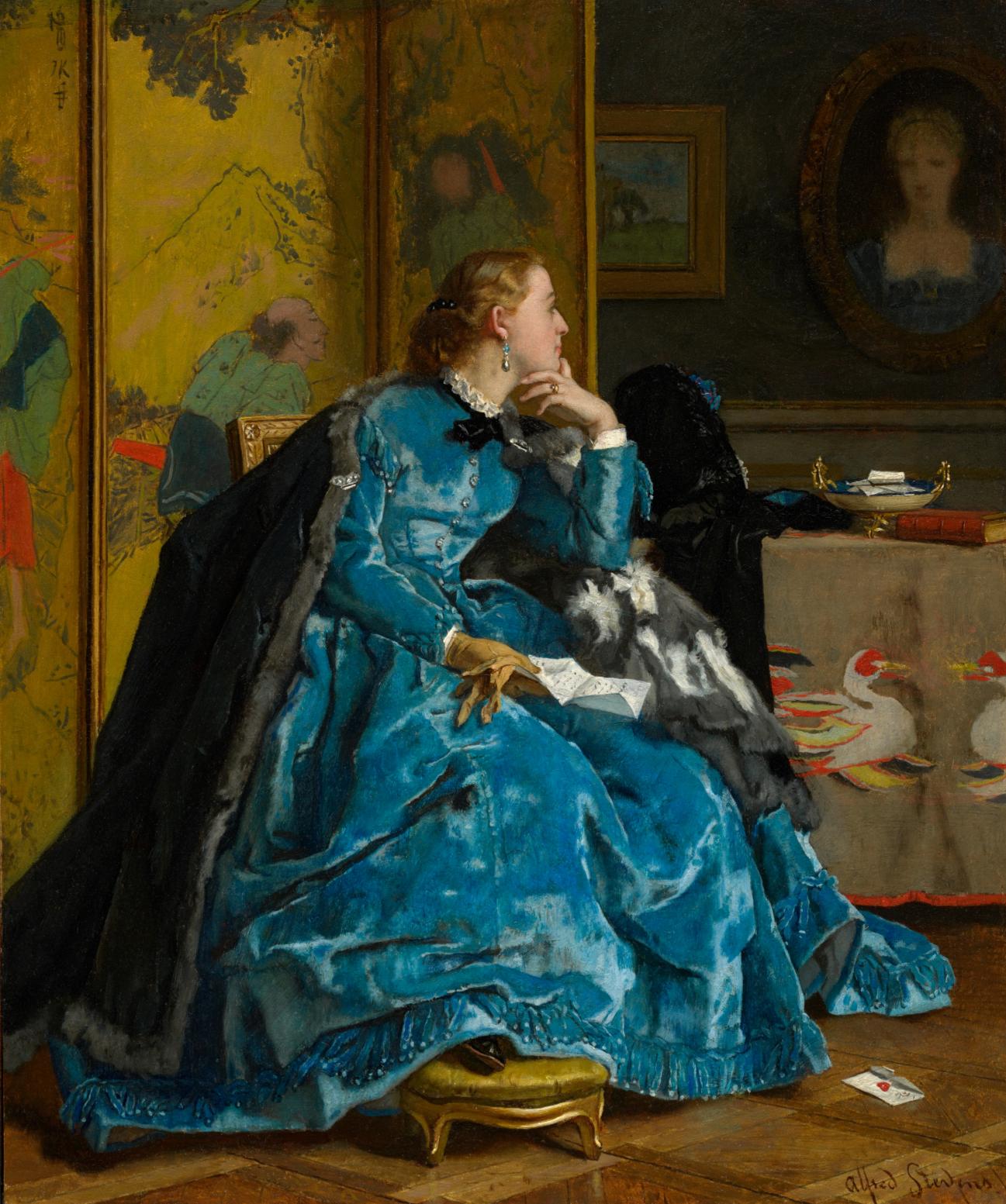

### 1960.108_print.jpg

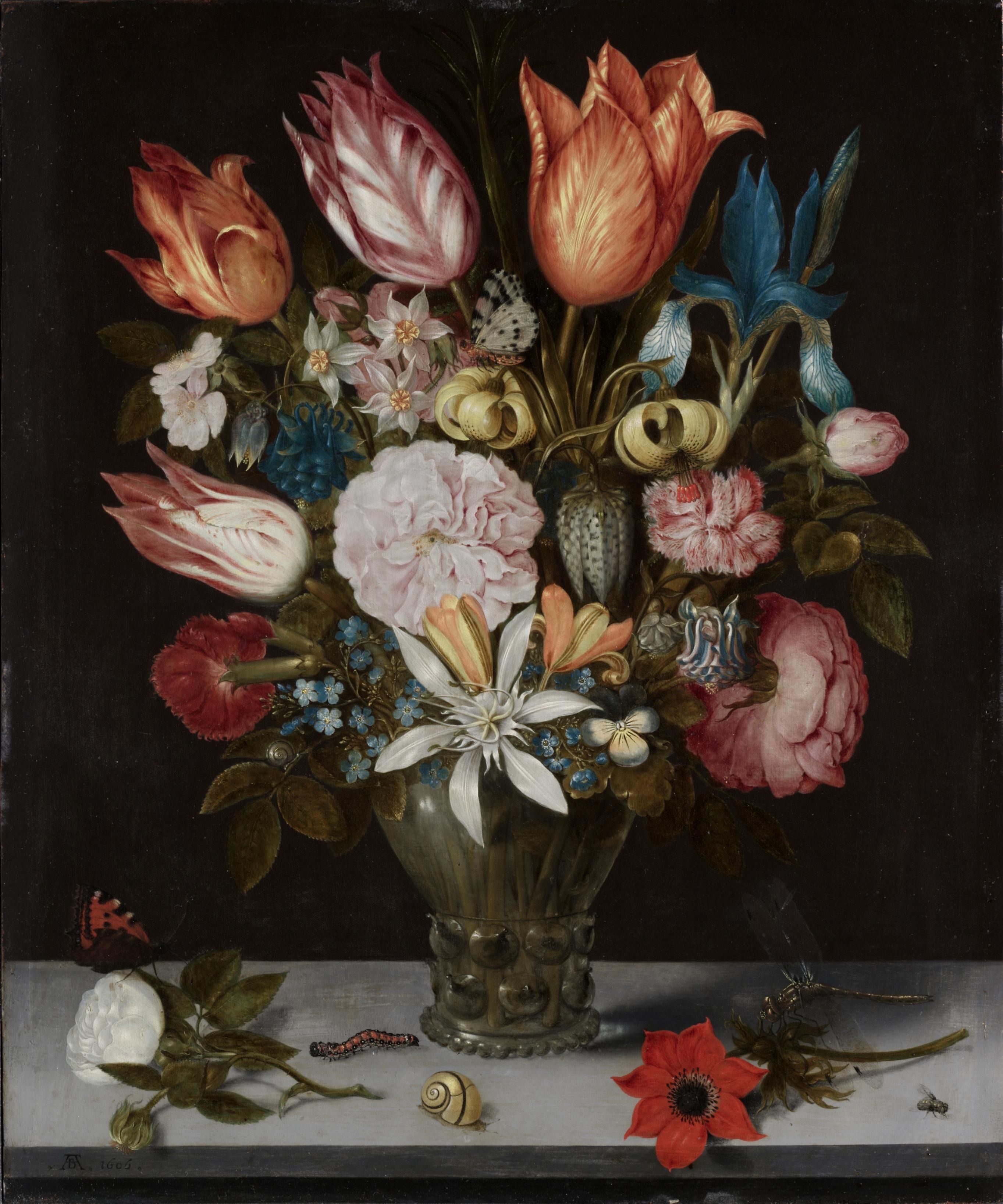

### 1989.333_print.jpg

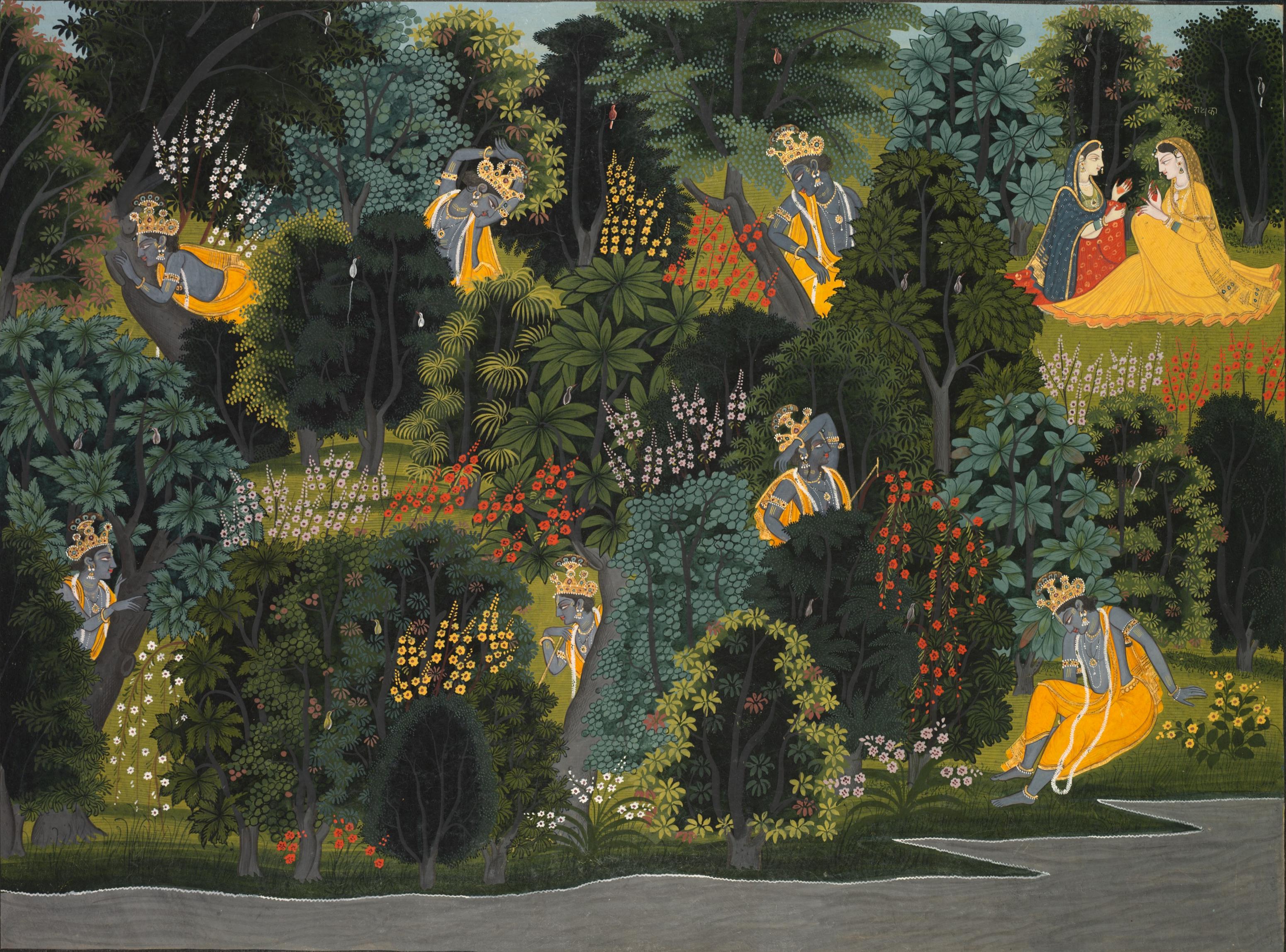

### 2002.jpg

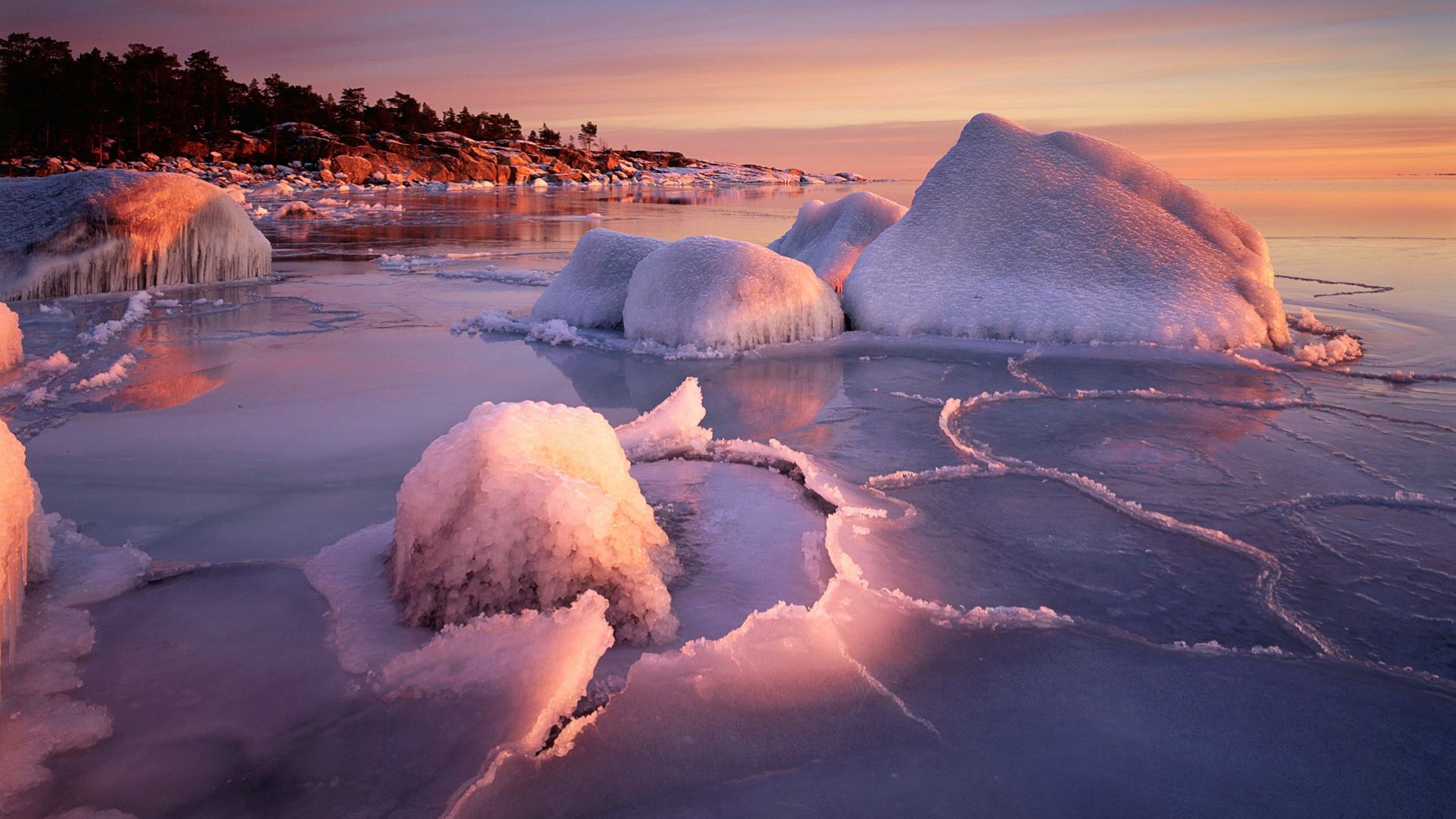

### 2003.jpg

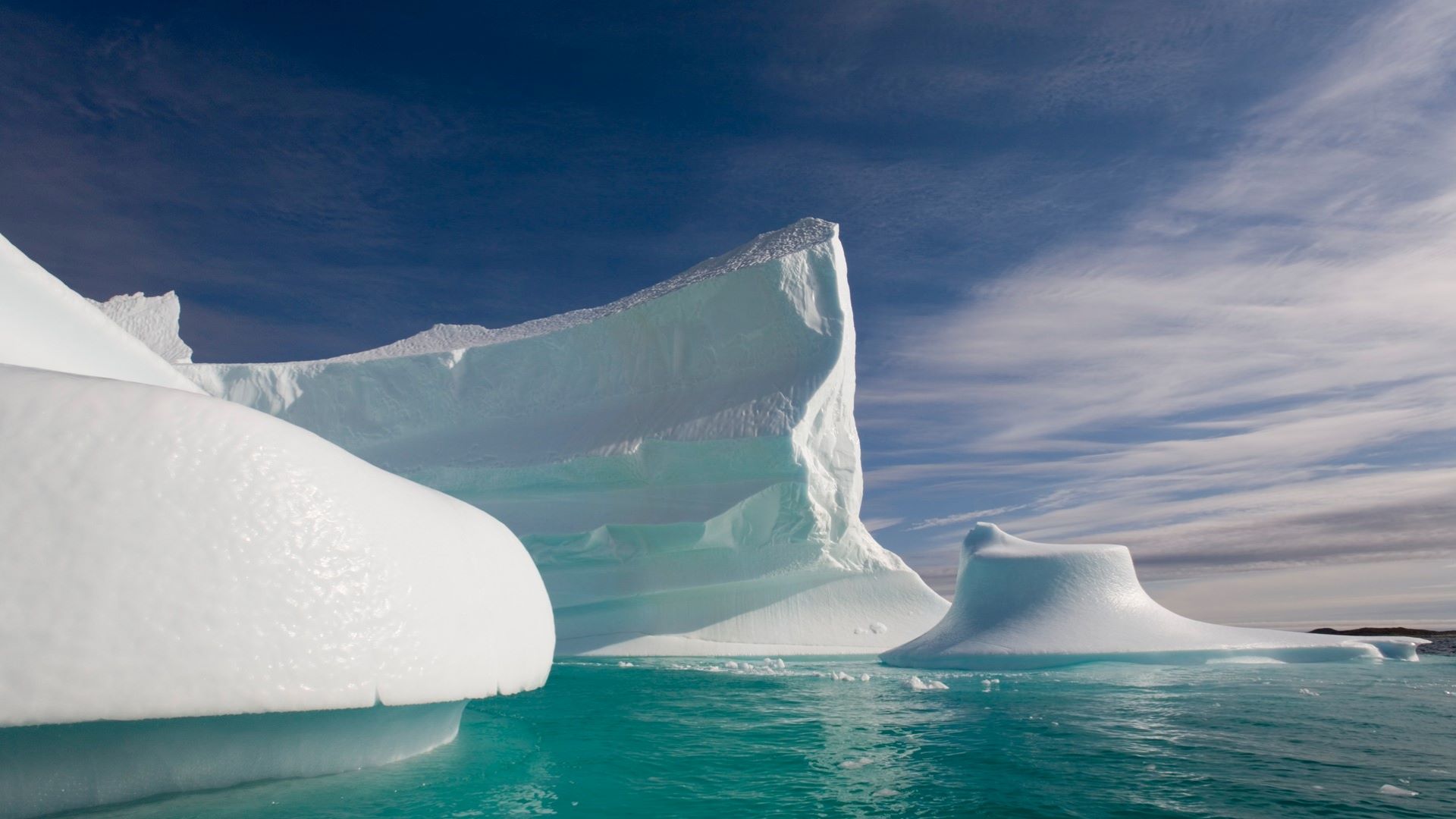

### 2004.jpg

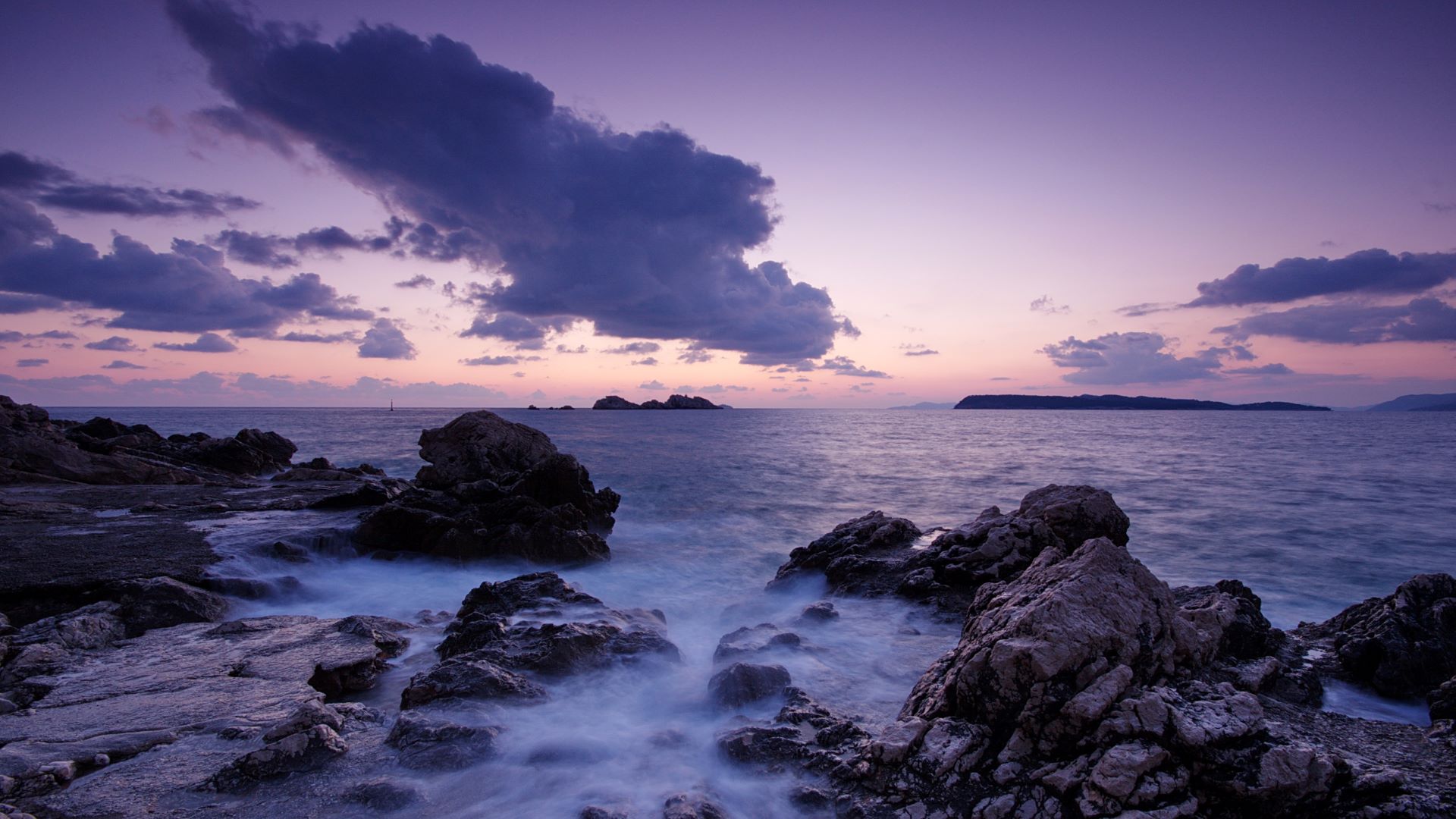

### 2006.jpg

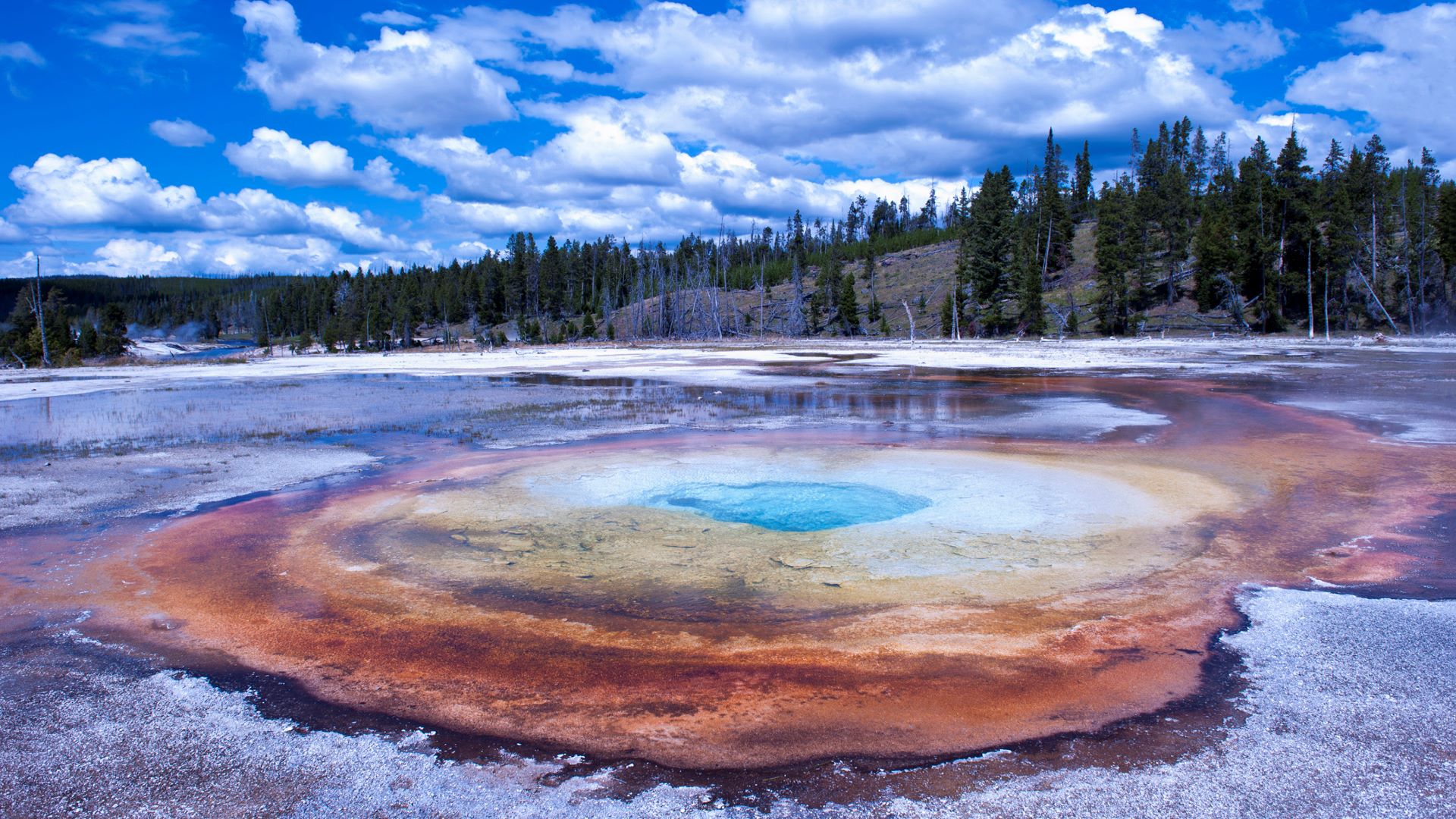

### 2007.8.34.jpg

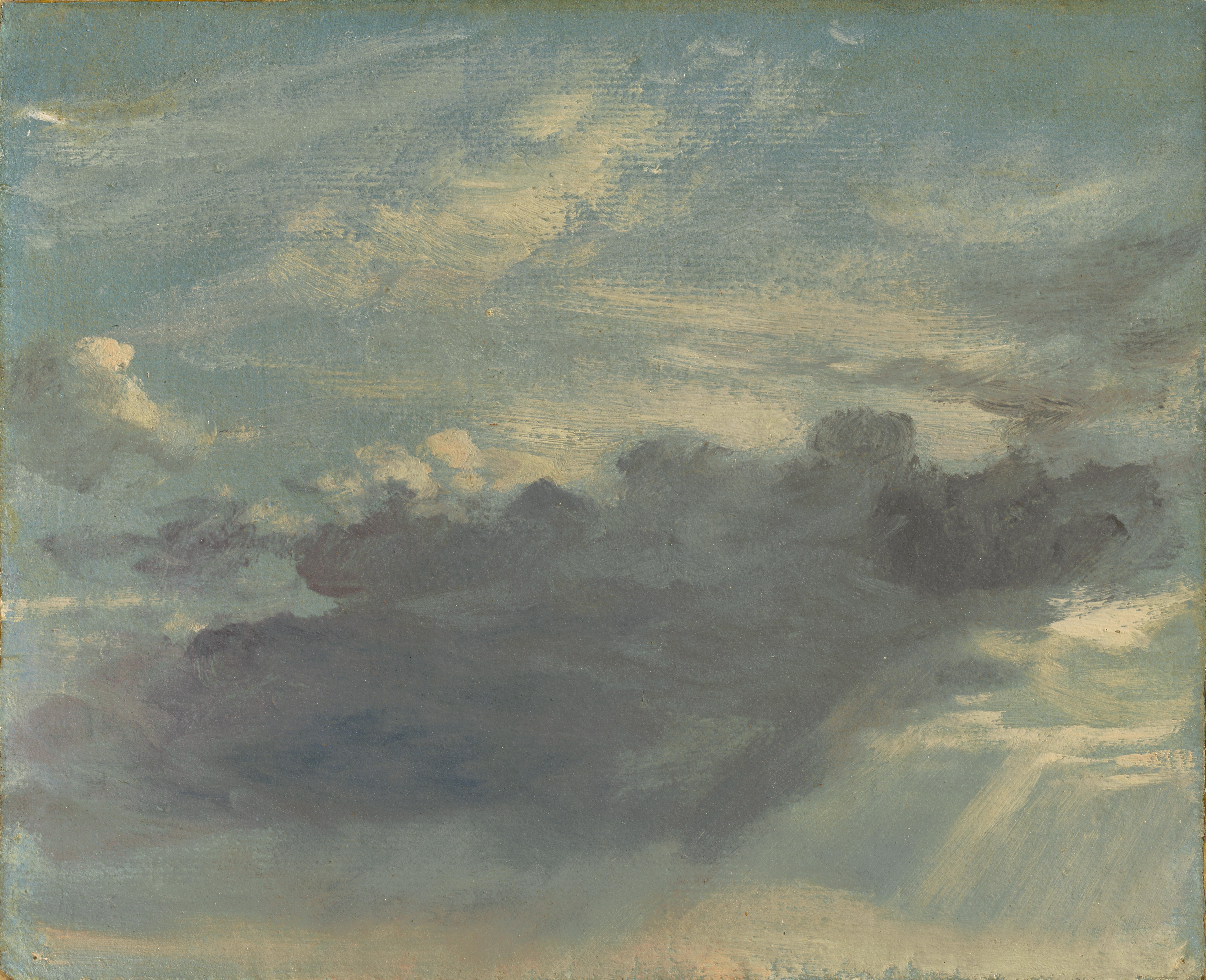

### 2009.jpg

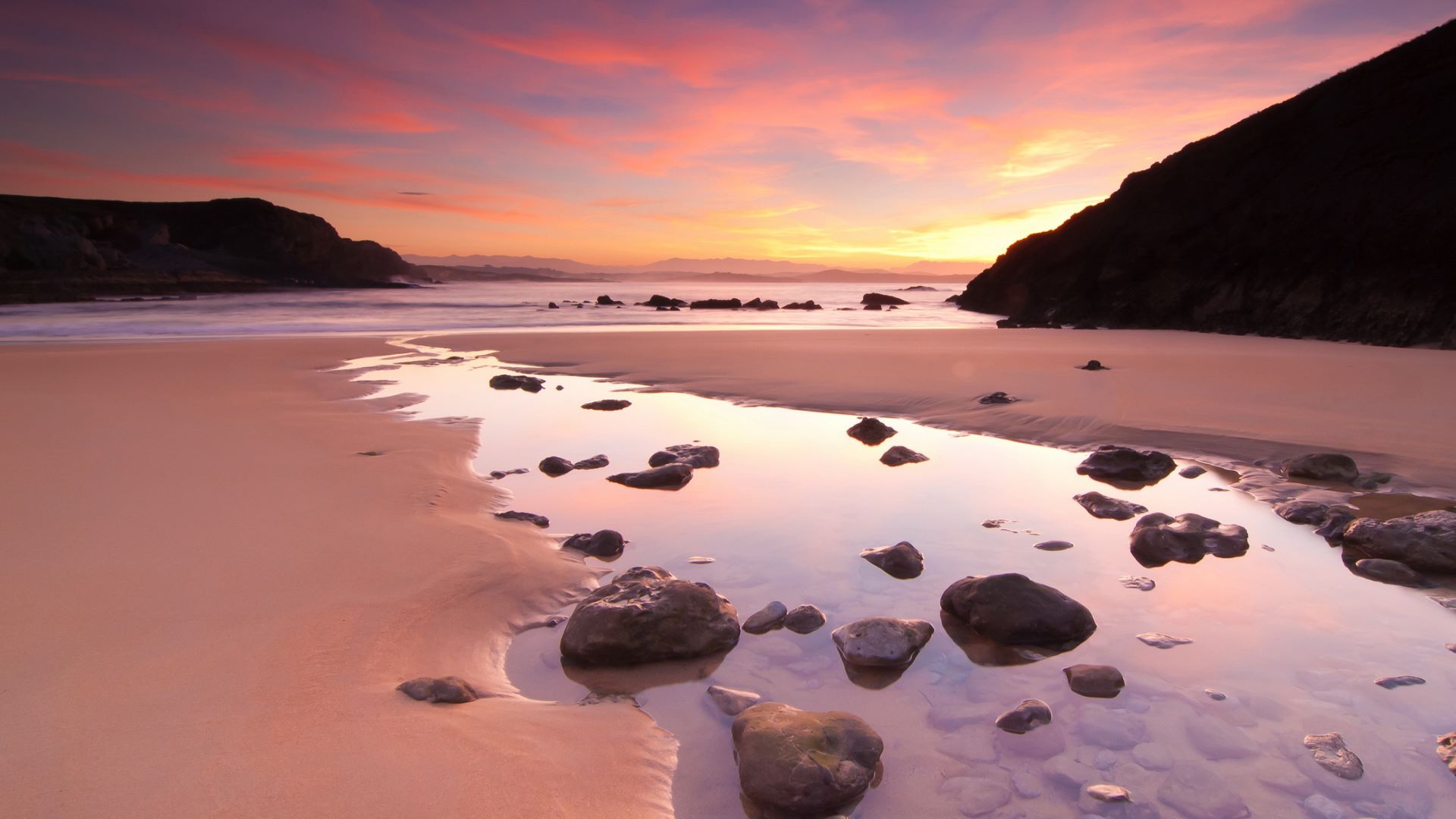

### 2016.jpg

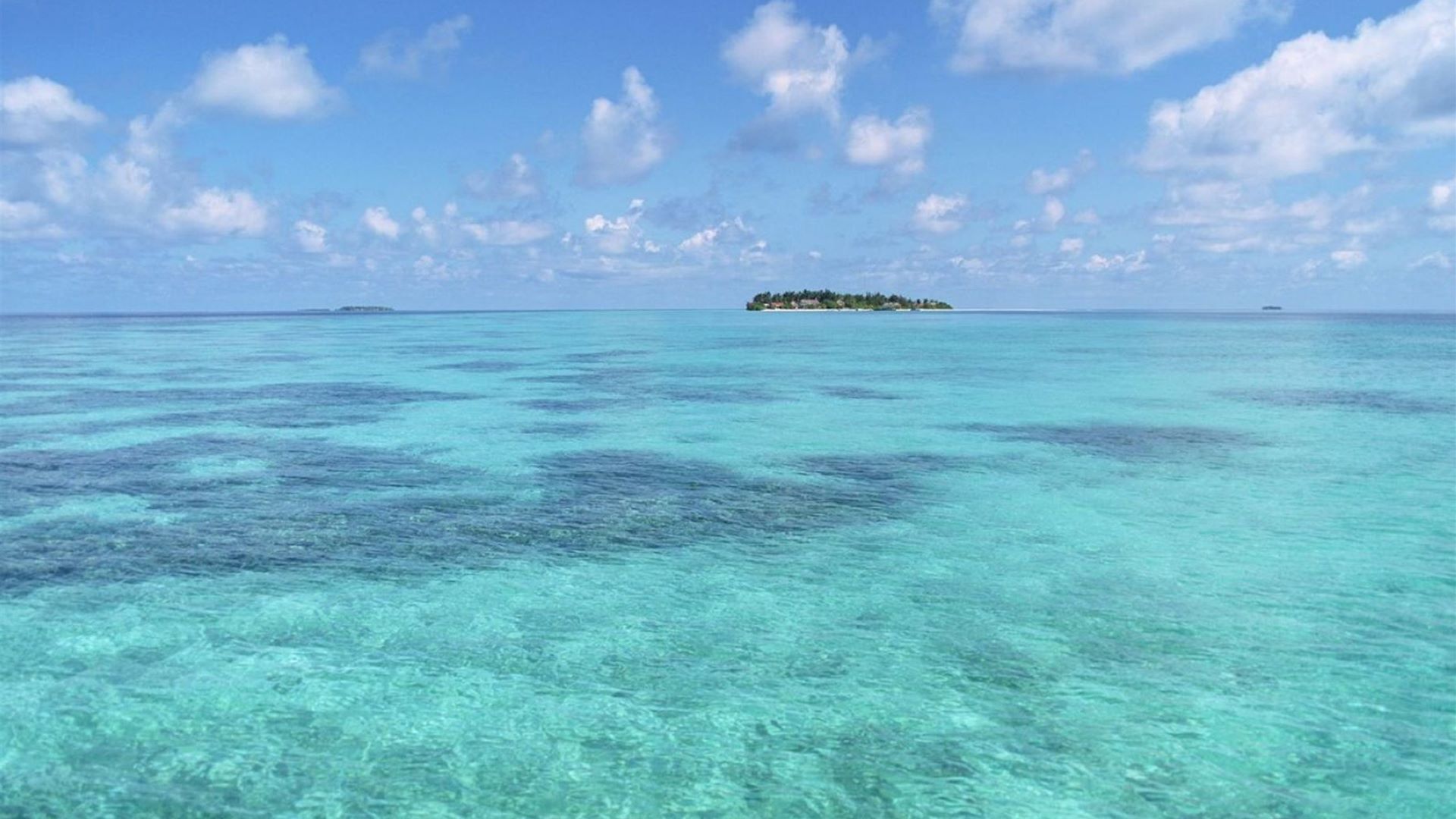

### 2017.jpg

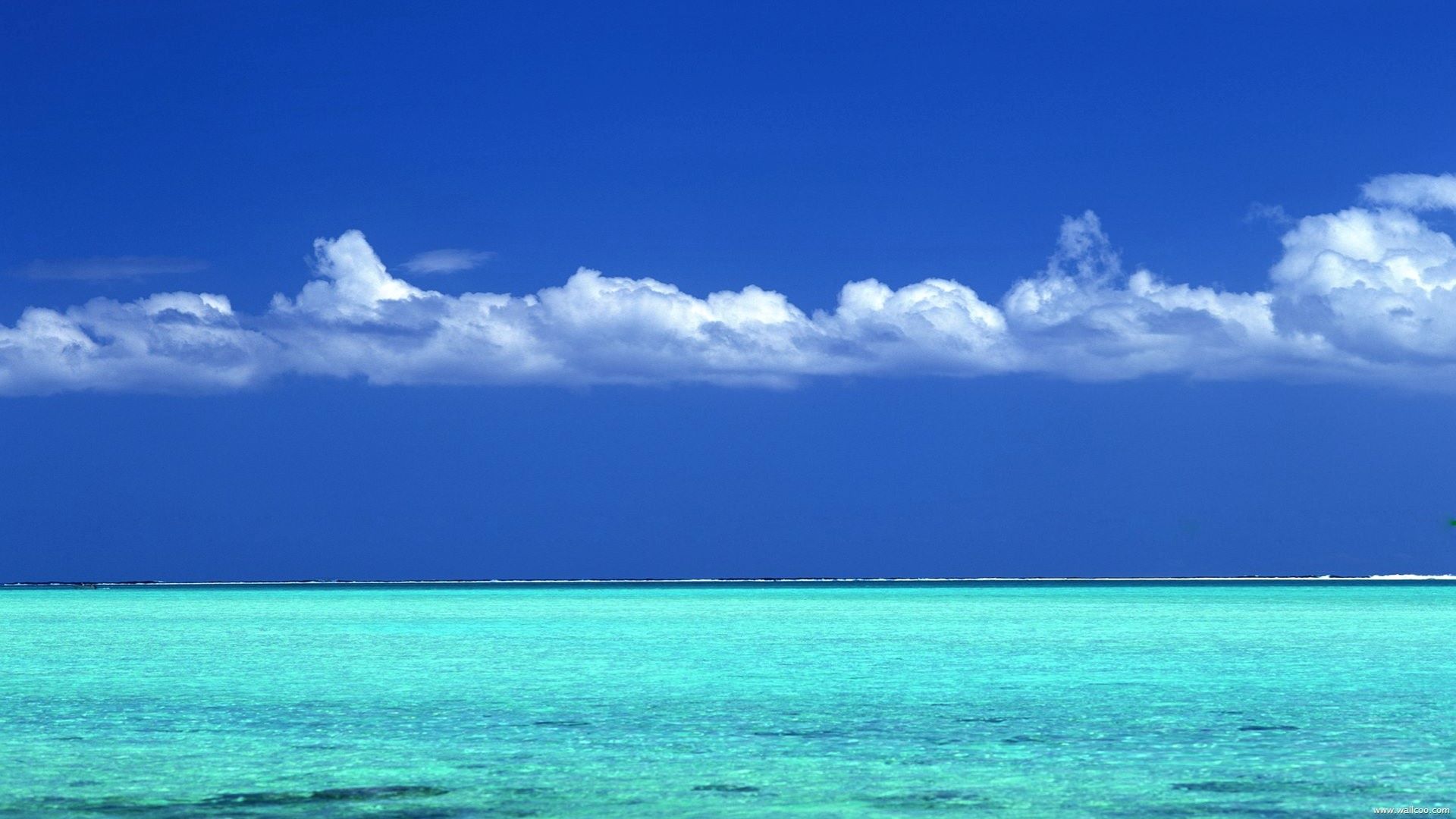

### 2020.jpg

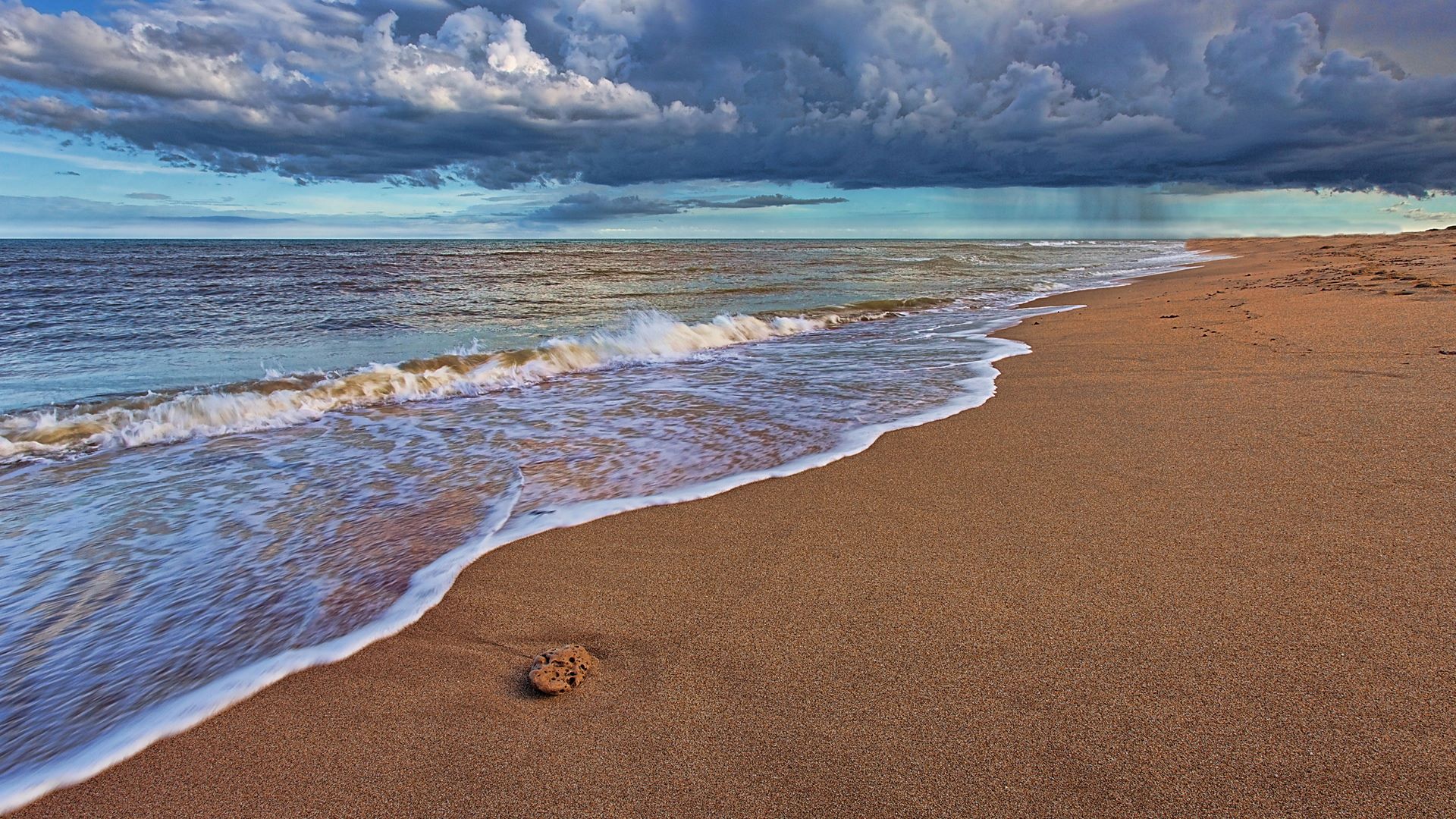

### 2021.jpg

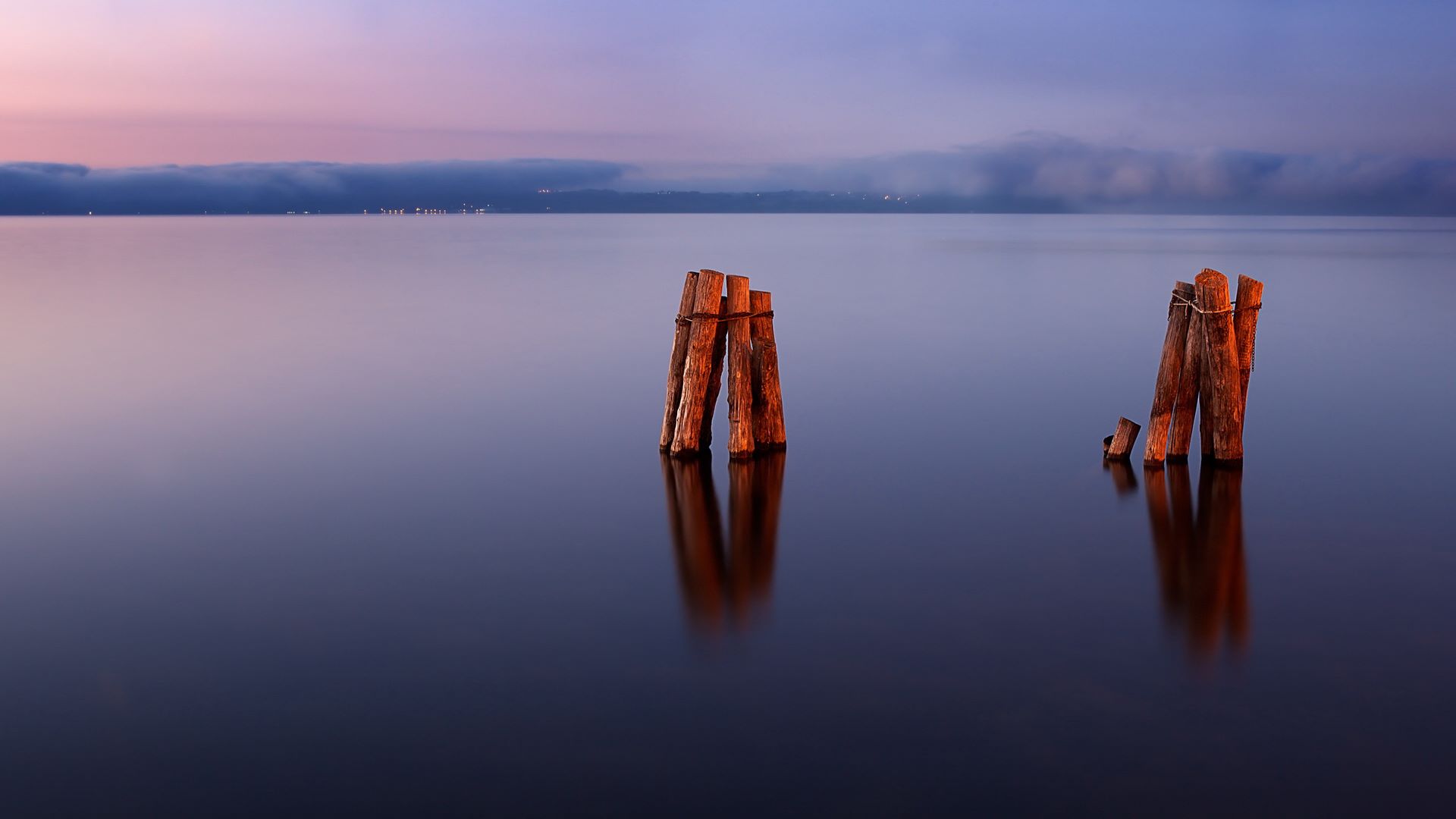

### 2022.jpg

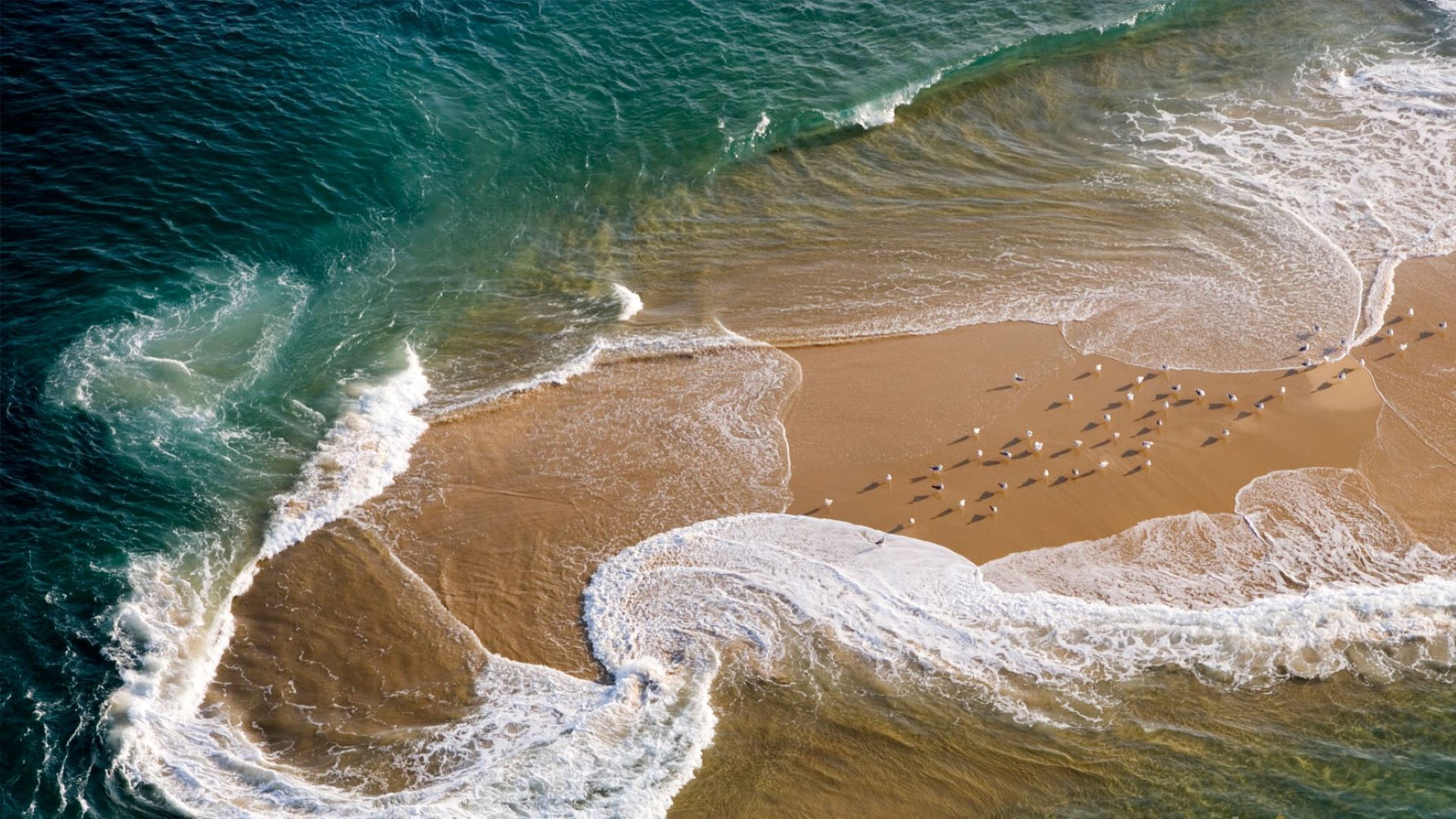

### 2025.jpg

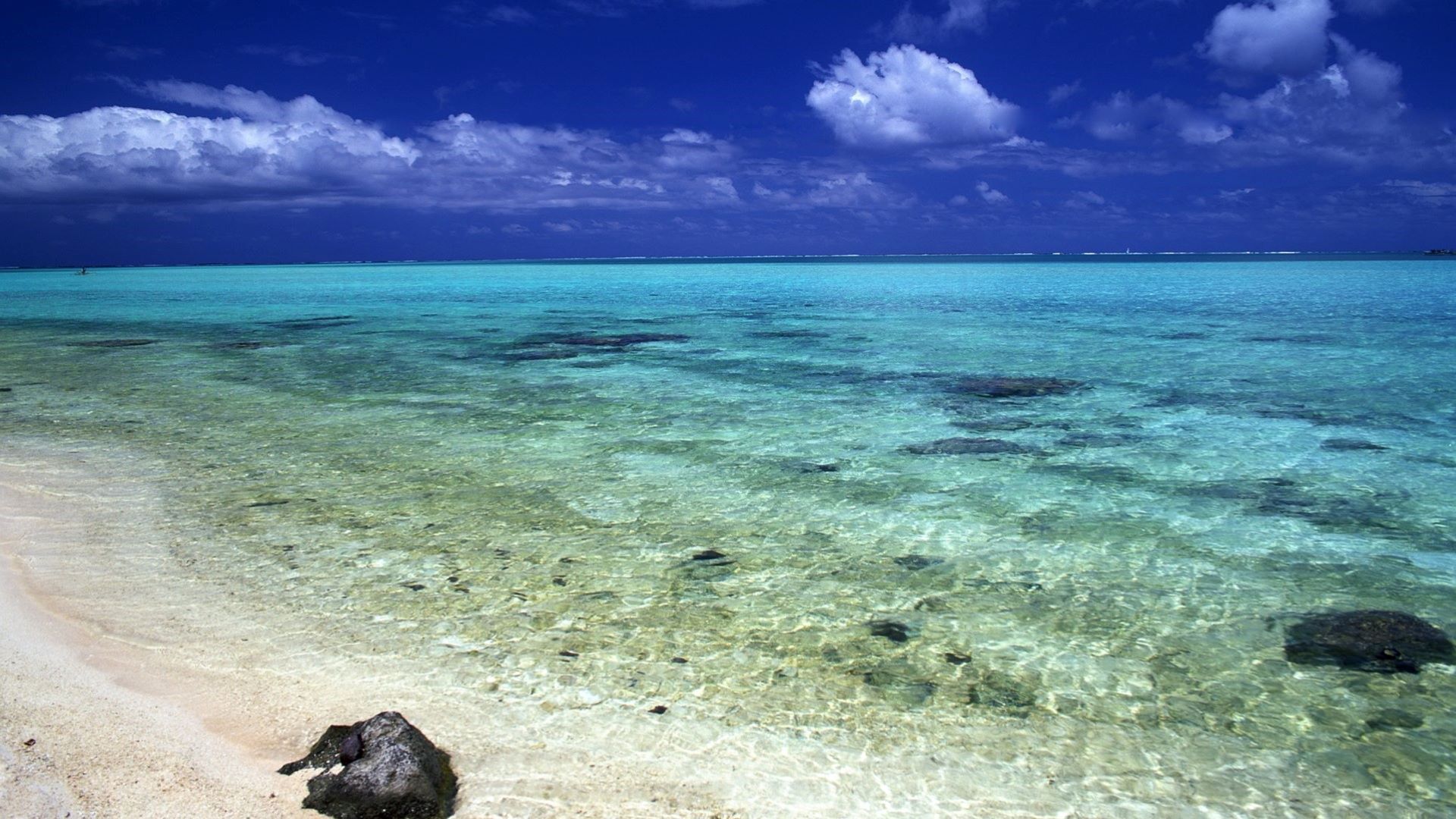

### 2041.jpg

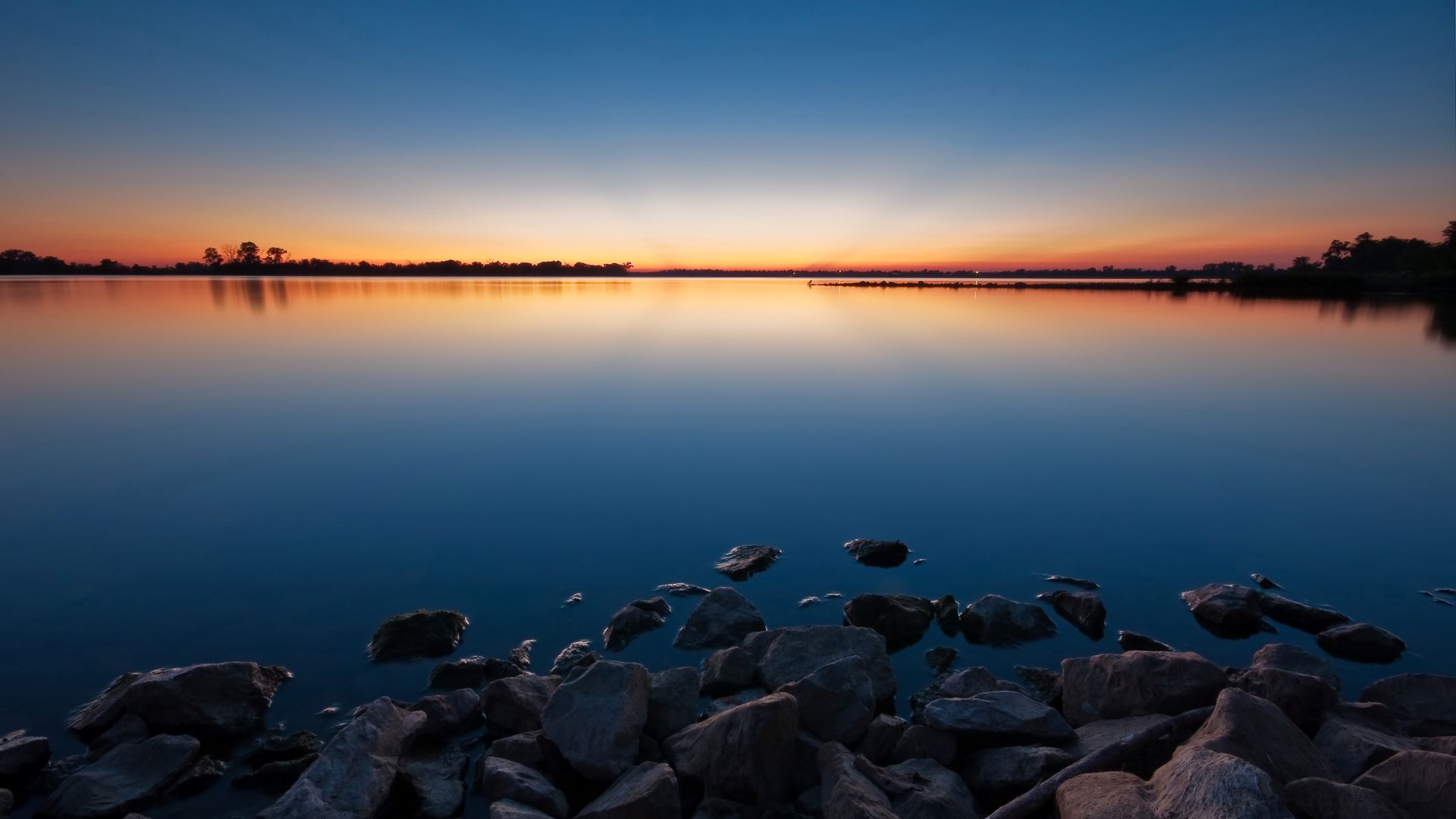

### 219179fgsdl.jpg

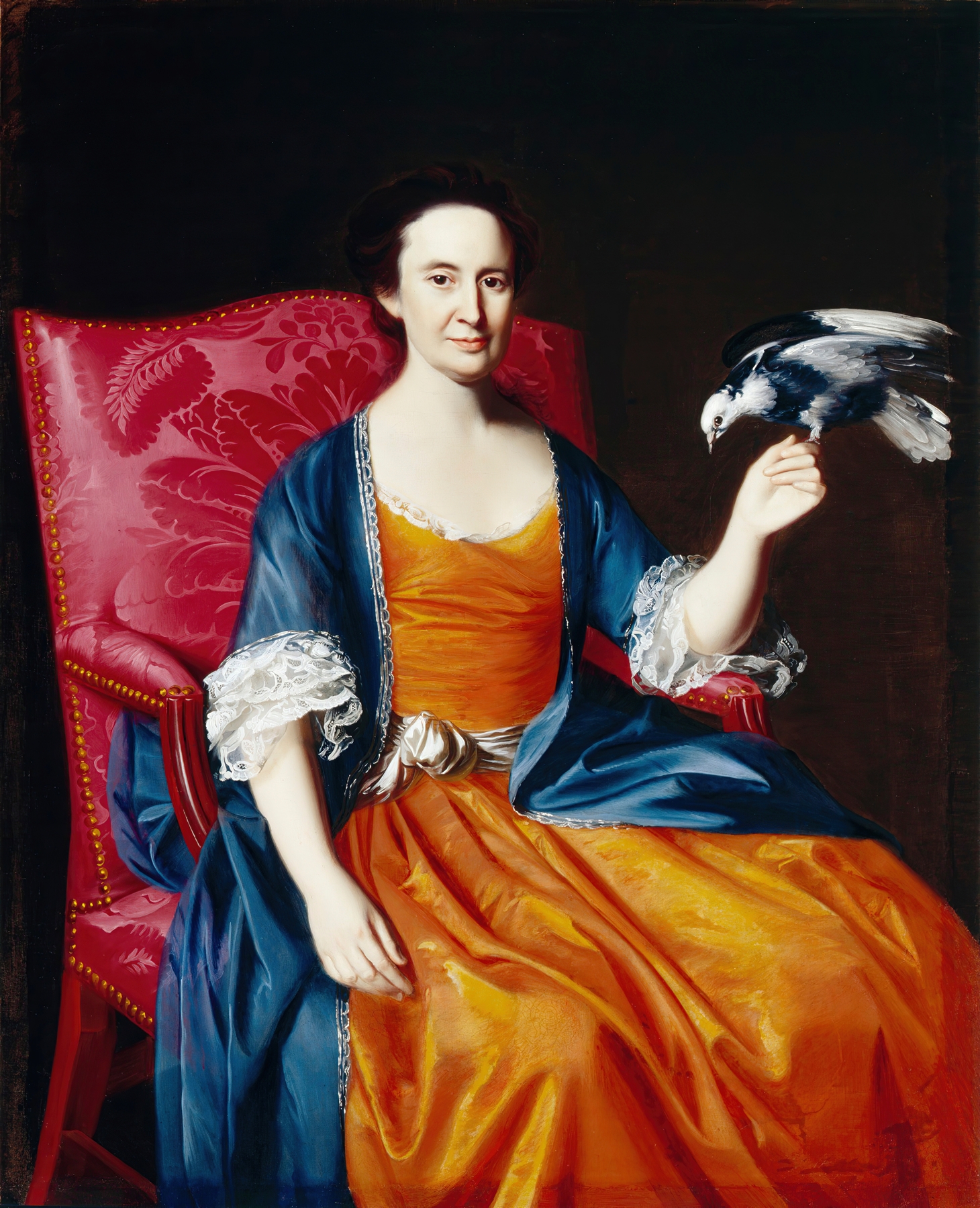

### 6408482614a1782e244e3434f3da5df9.png

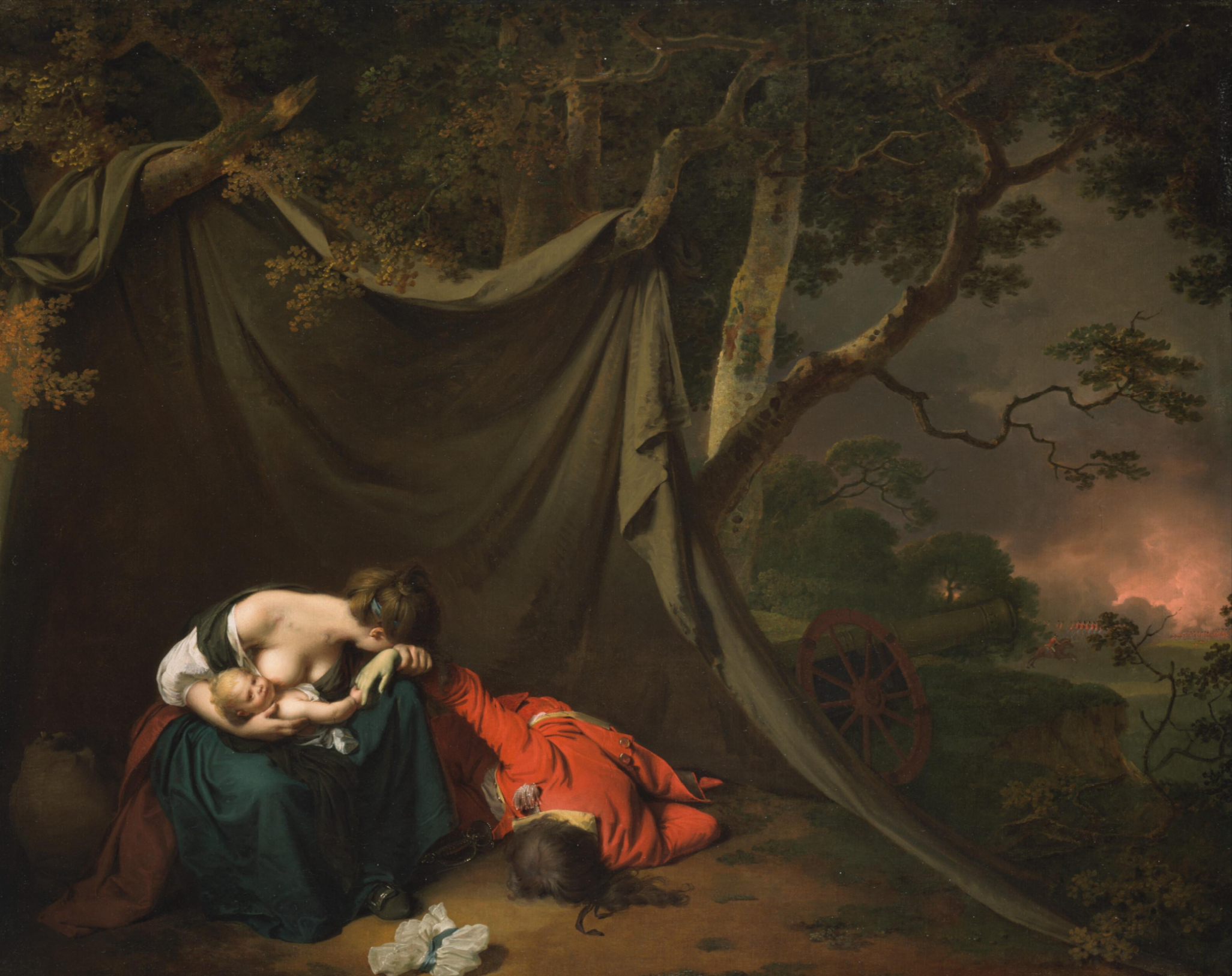

### al-held-i-beam-1961.jpg

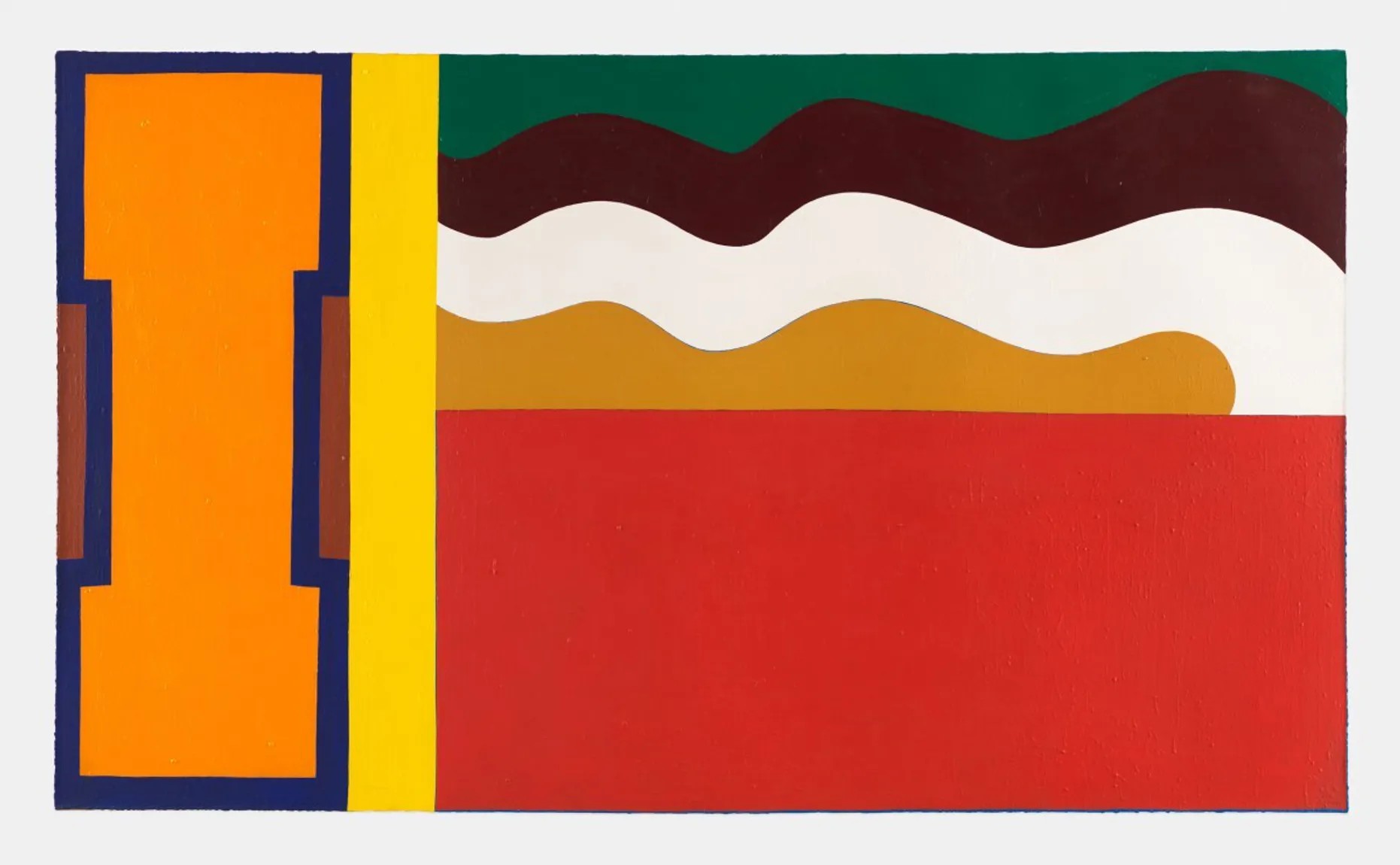

### cb3389b8-d606-42b1-a88f-97959280f6bb_2549.jpg

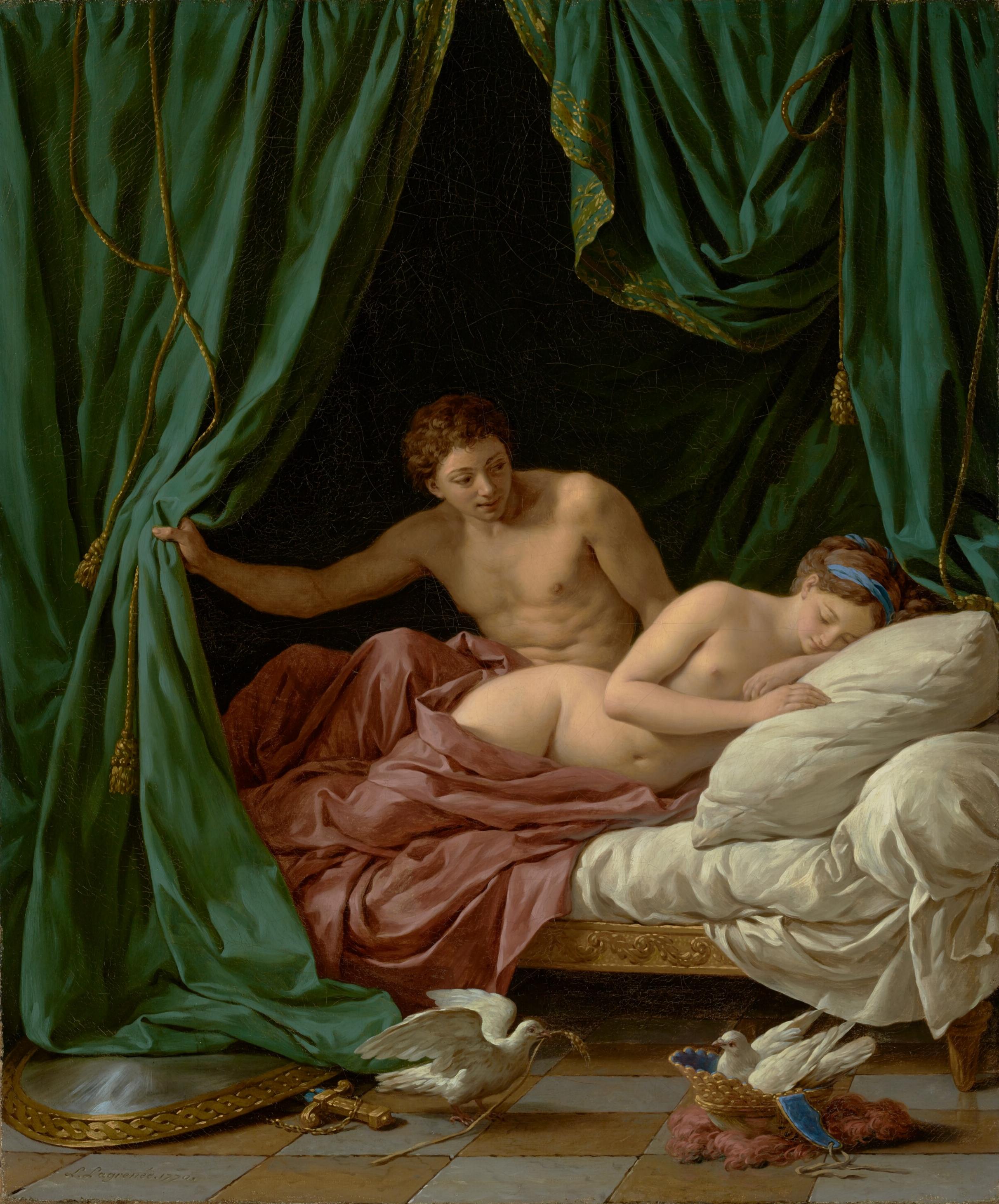

### Dante_Gabriel_Rossetti_-_Salutation_of_Beatrice_-_2.jpg

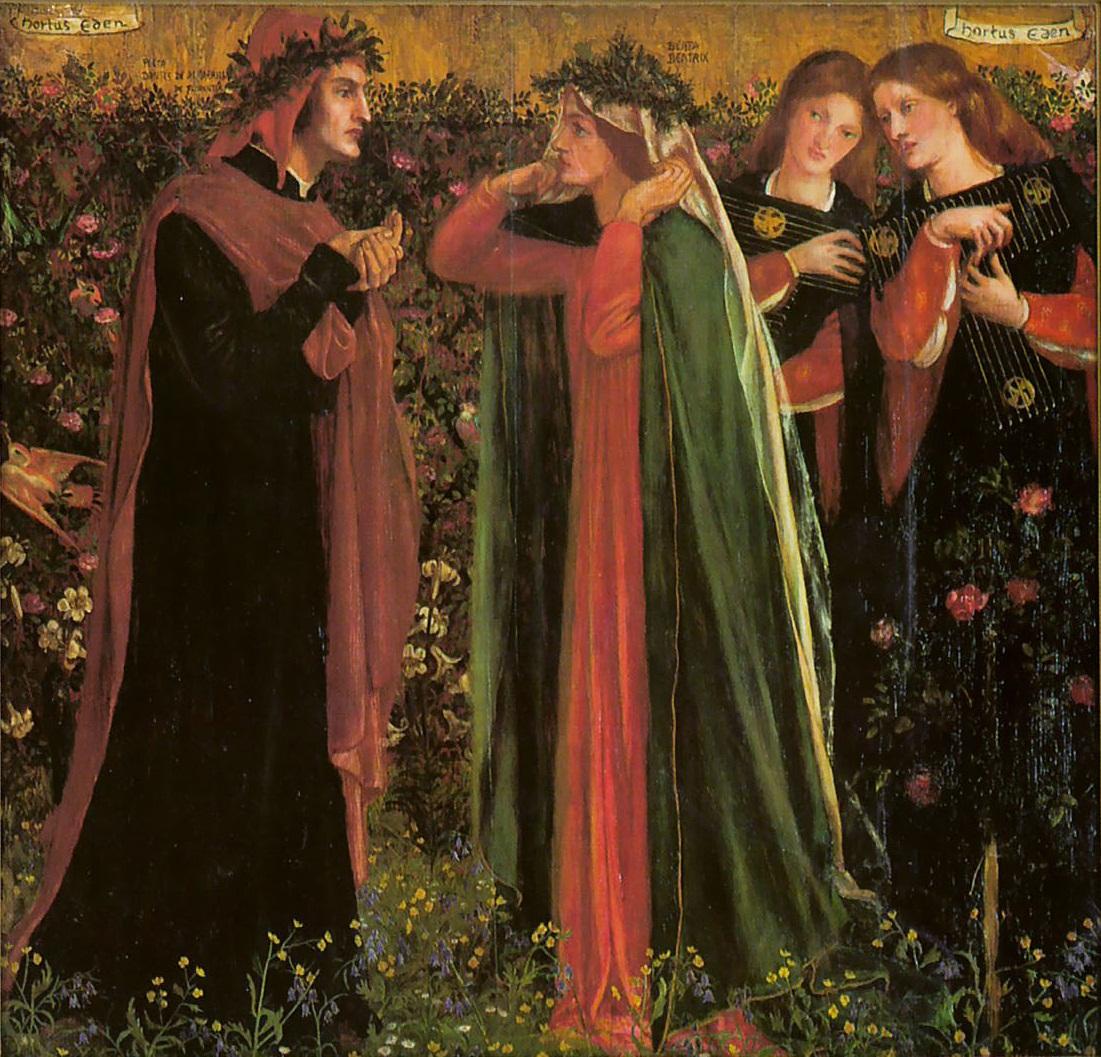

### default.jpg

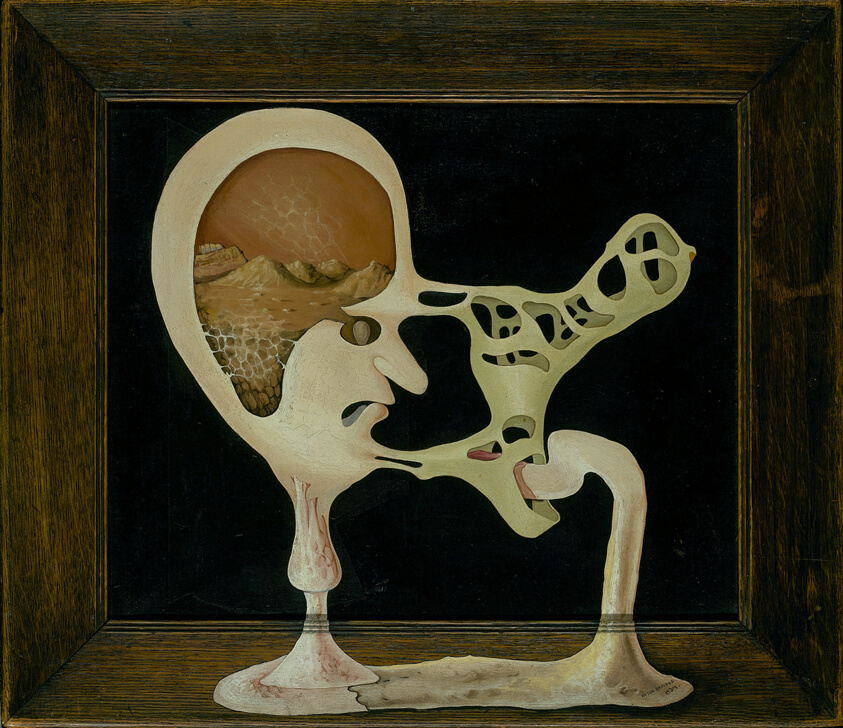

### DT75.jpg

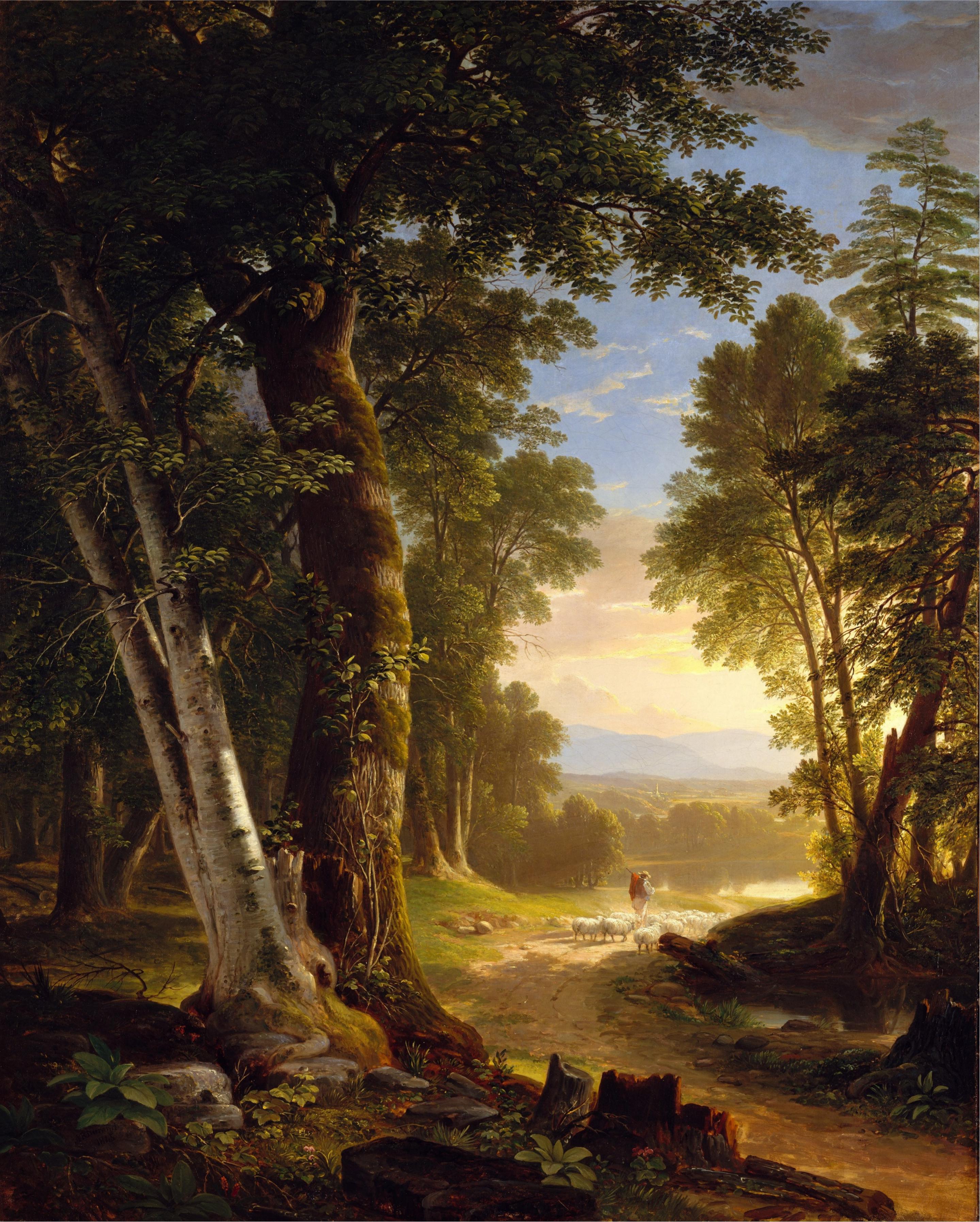

### Eugène_Louis_Gabriel_Isabey_-_The_Wreck_-_07.14_-_Detroit_Institute_of_Arts.jpg

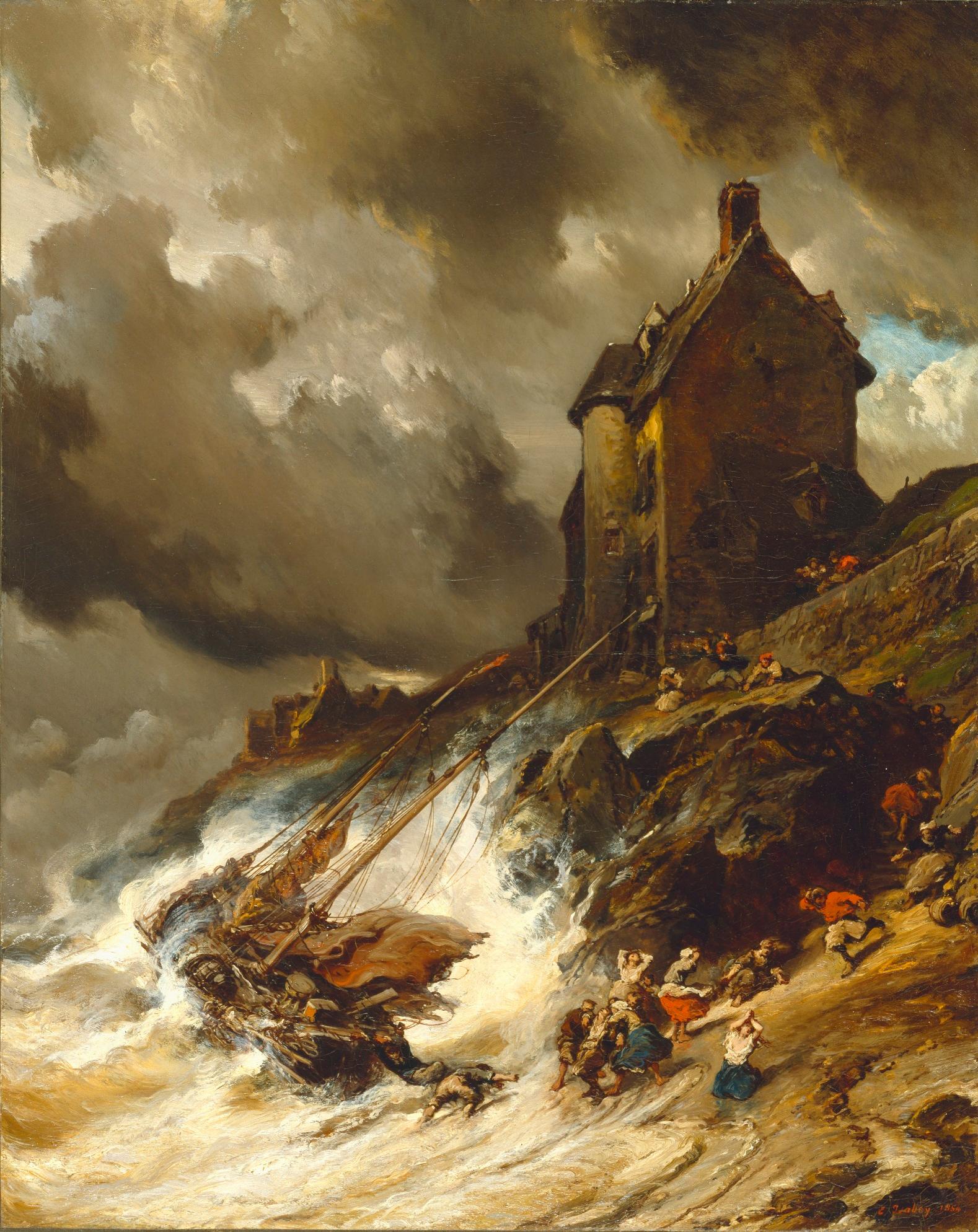

### Evocation of Butterflies.jpg

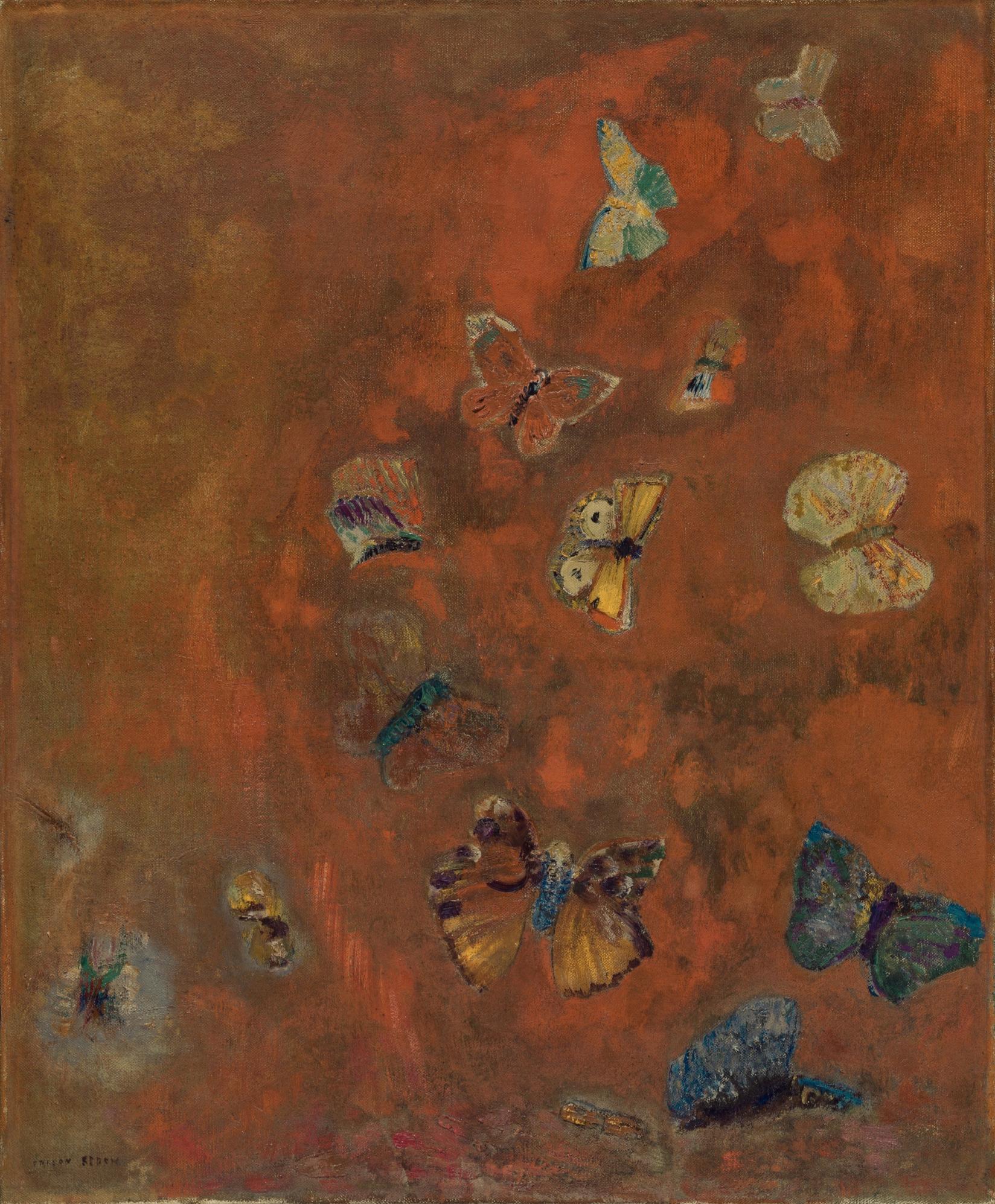

### Ferdinand-Victor-Eugène_Delacroix,_French_-_The_Death_of_Sardanapalus_-_Google_Art_Project.jpg

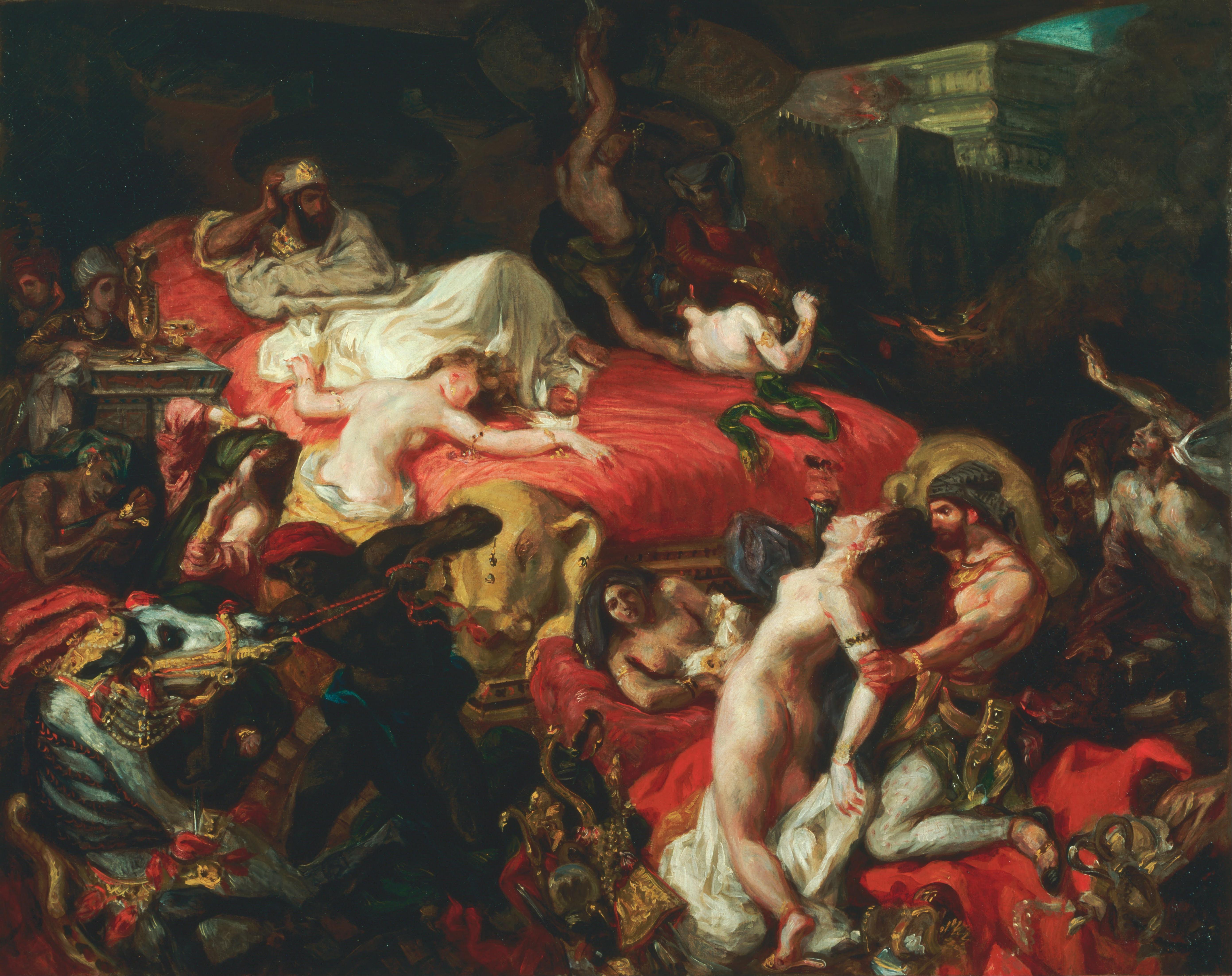

### forest_scene_1949.1.4.jpg

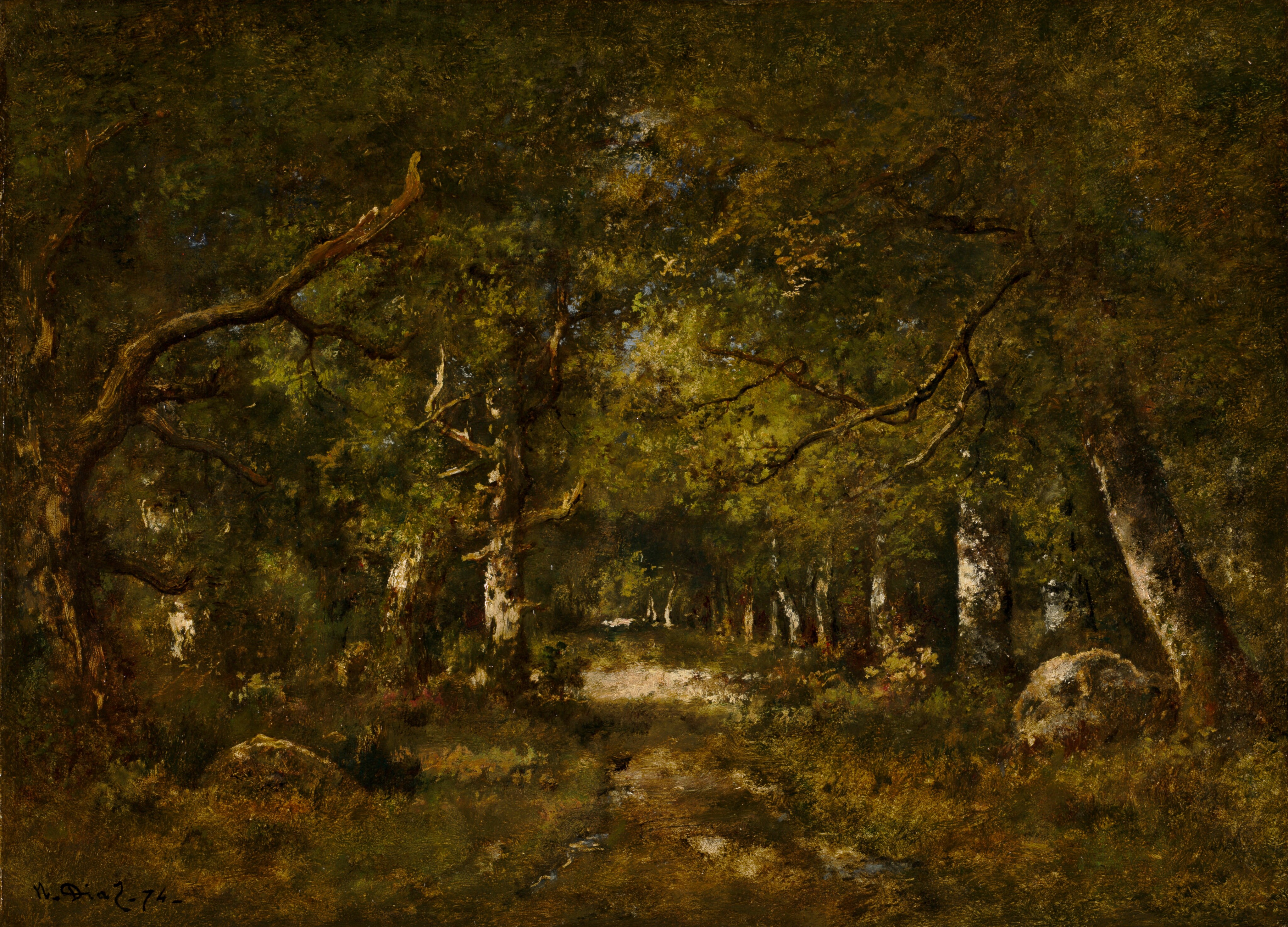

### Gustave_Courbet_-_La_vague_(c.1869-70).jpg

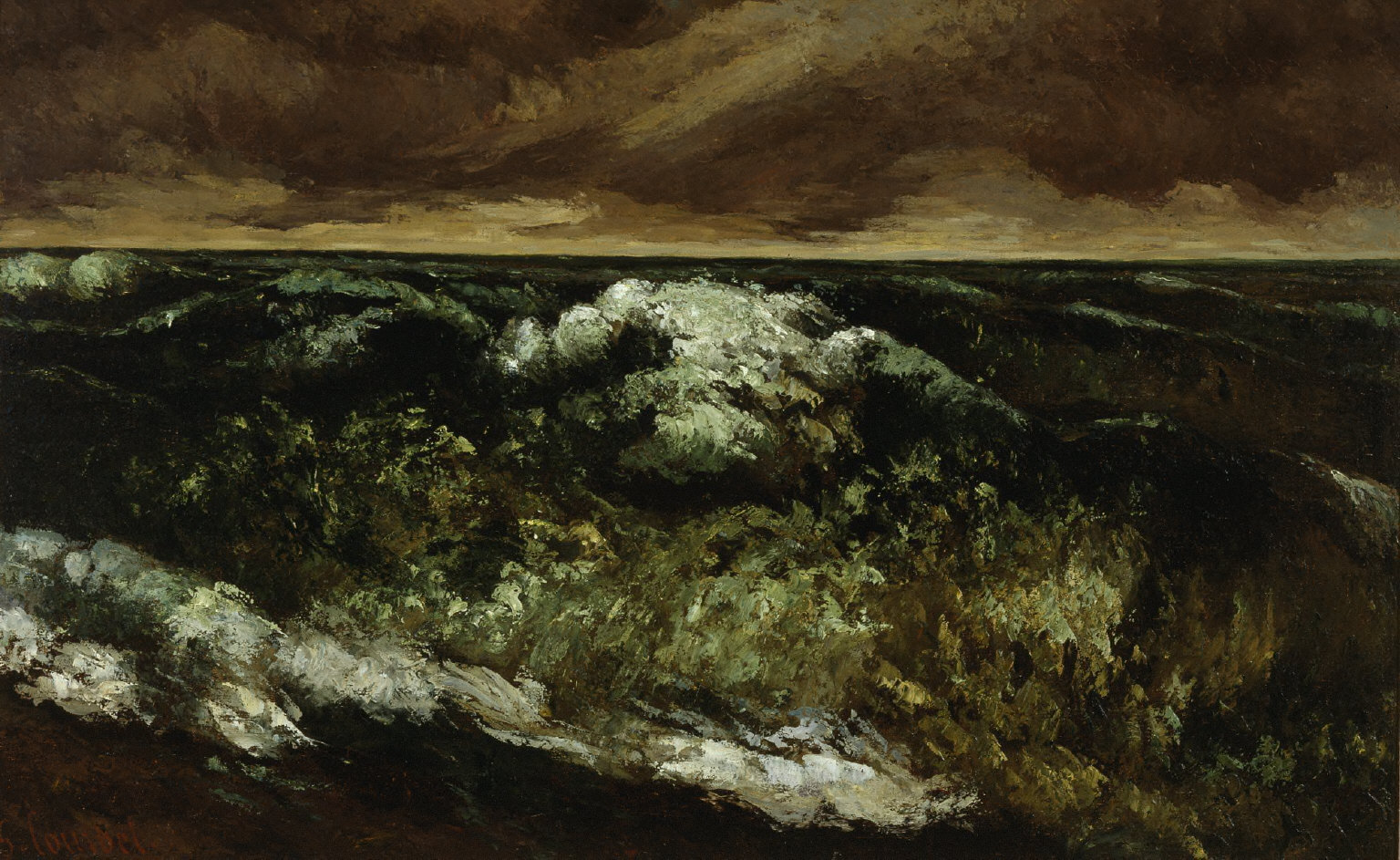

### hp1.jpg

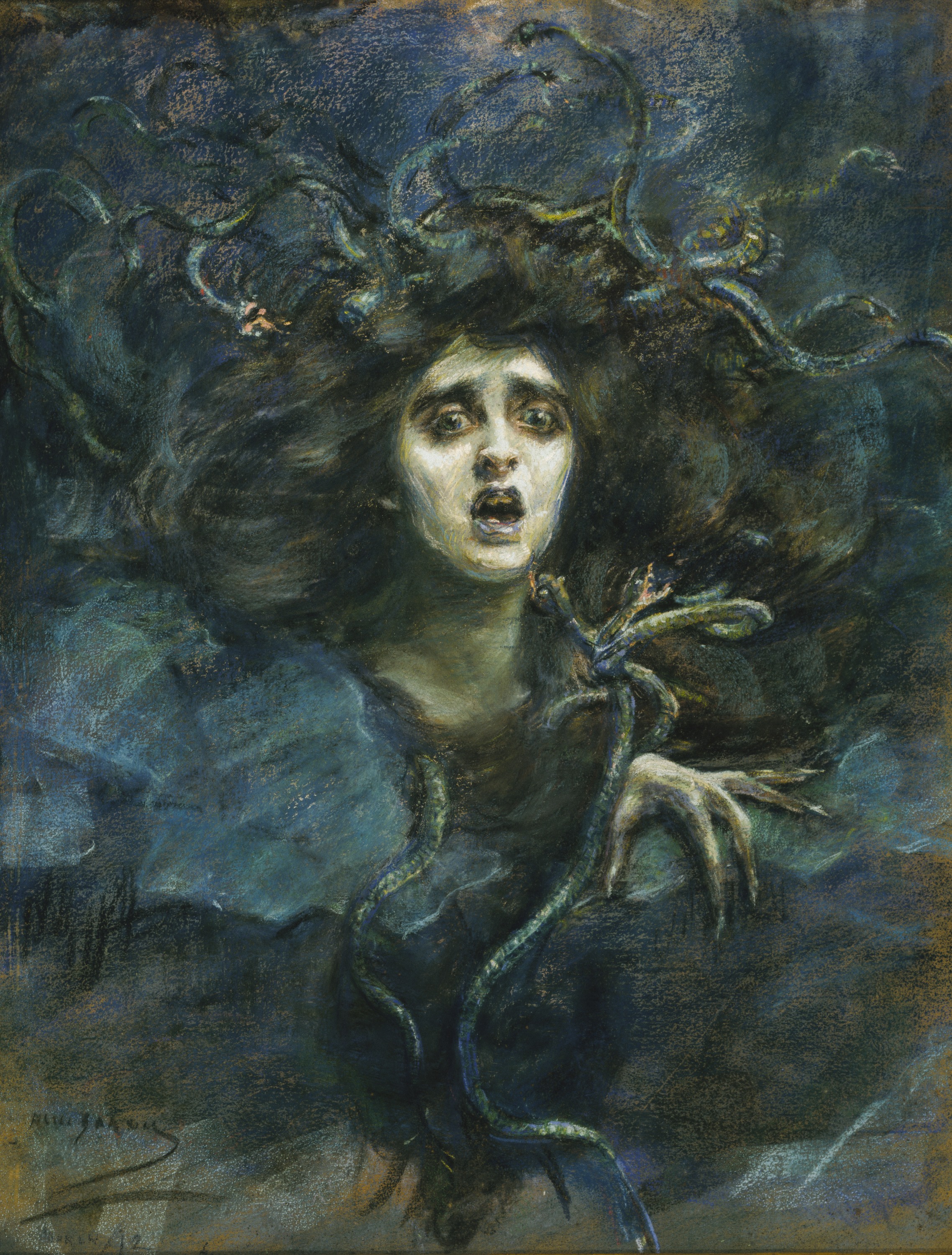

### hp7.jpg

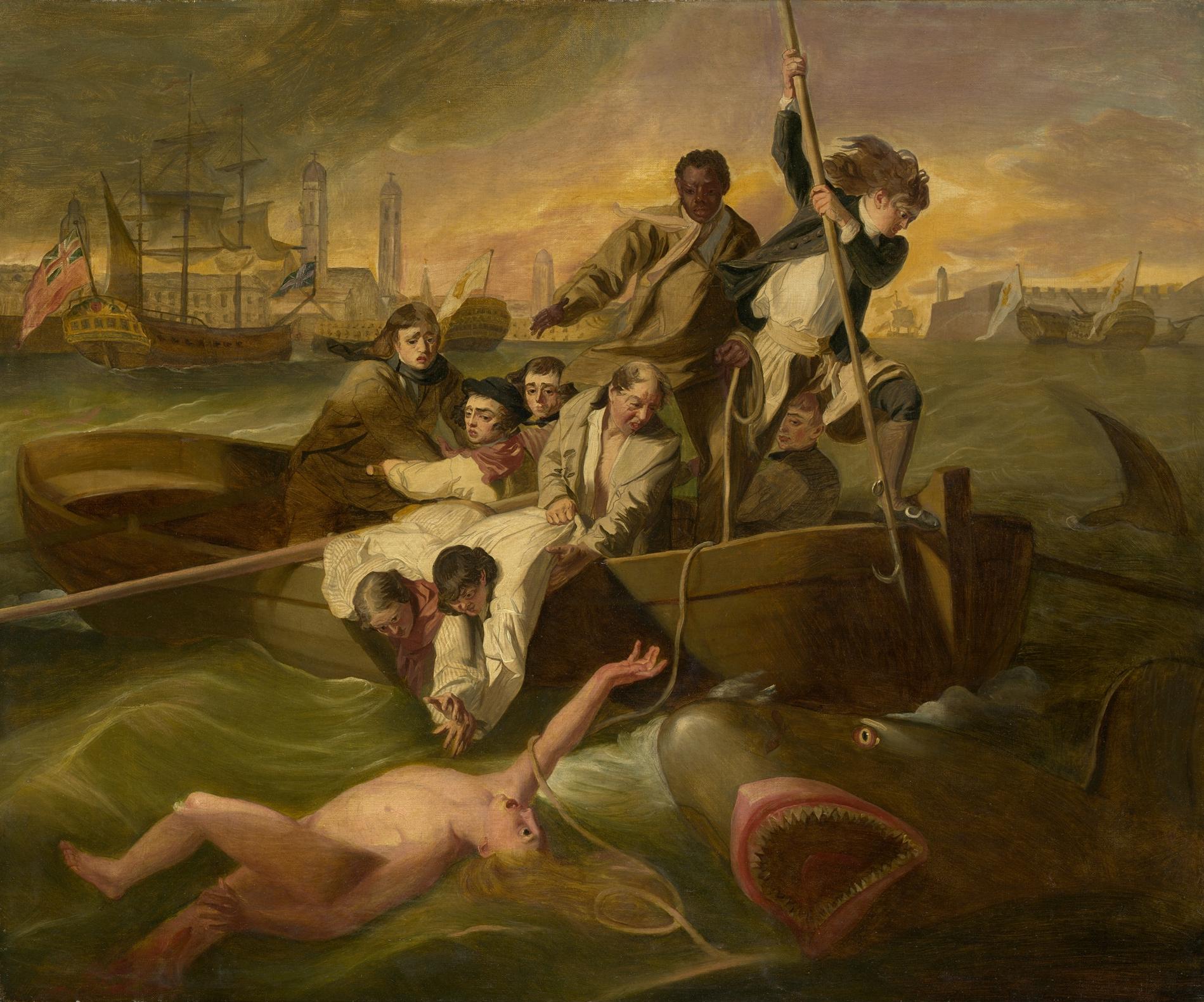

### hp8.png

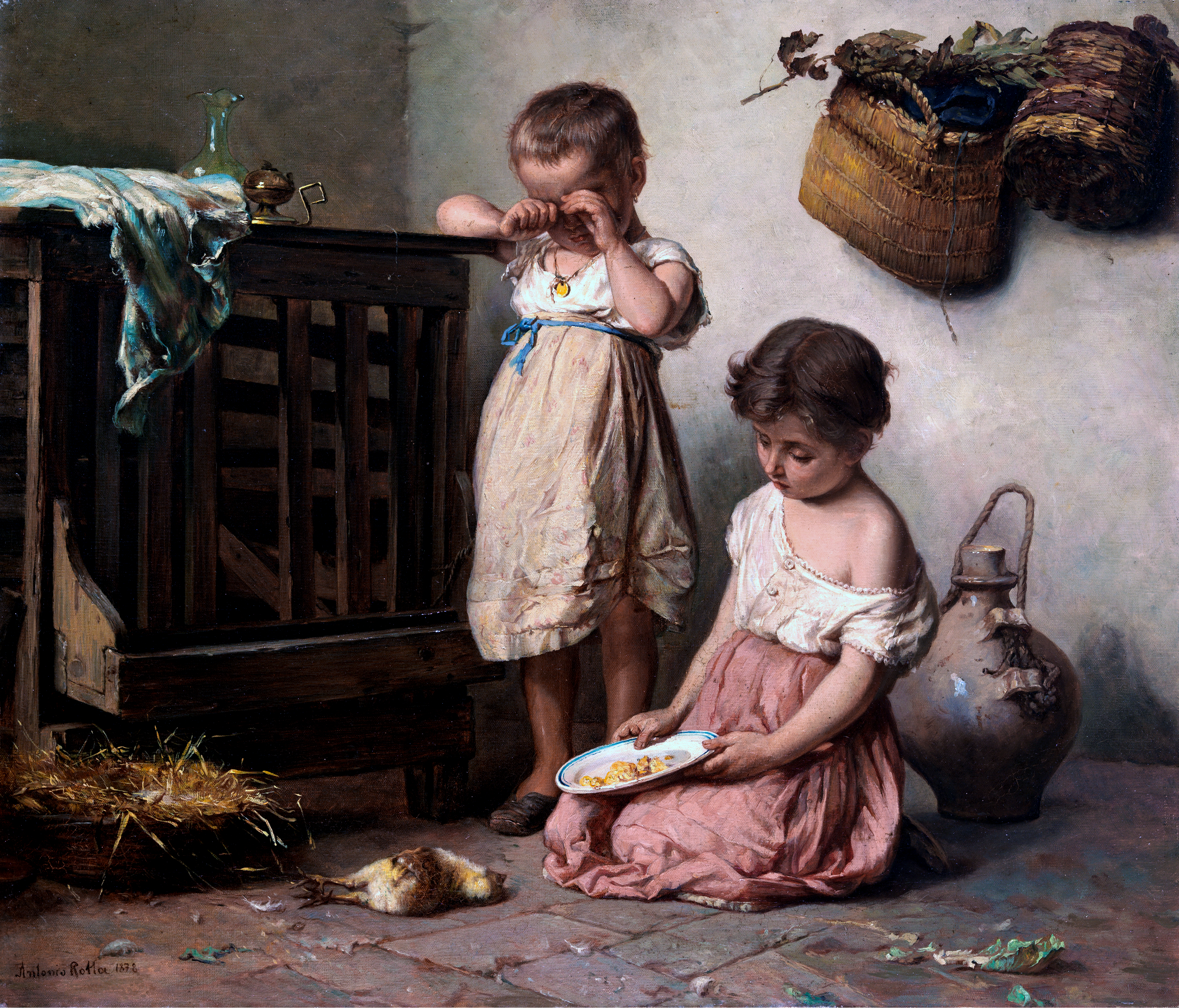

### John_Henry_Fuseli_-_The_Nightmare.jpg

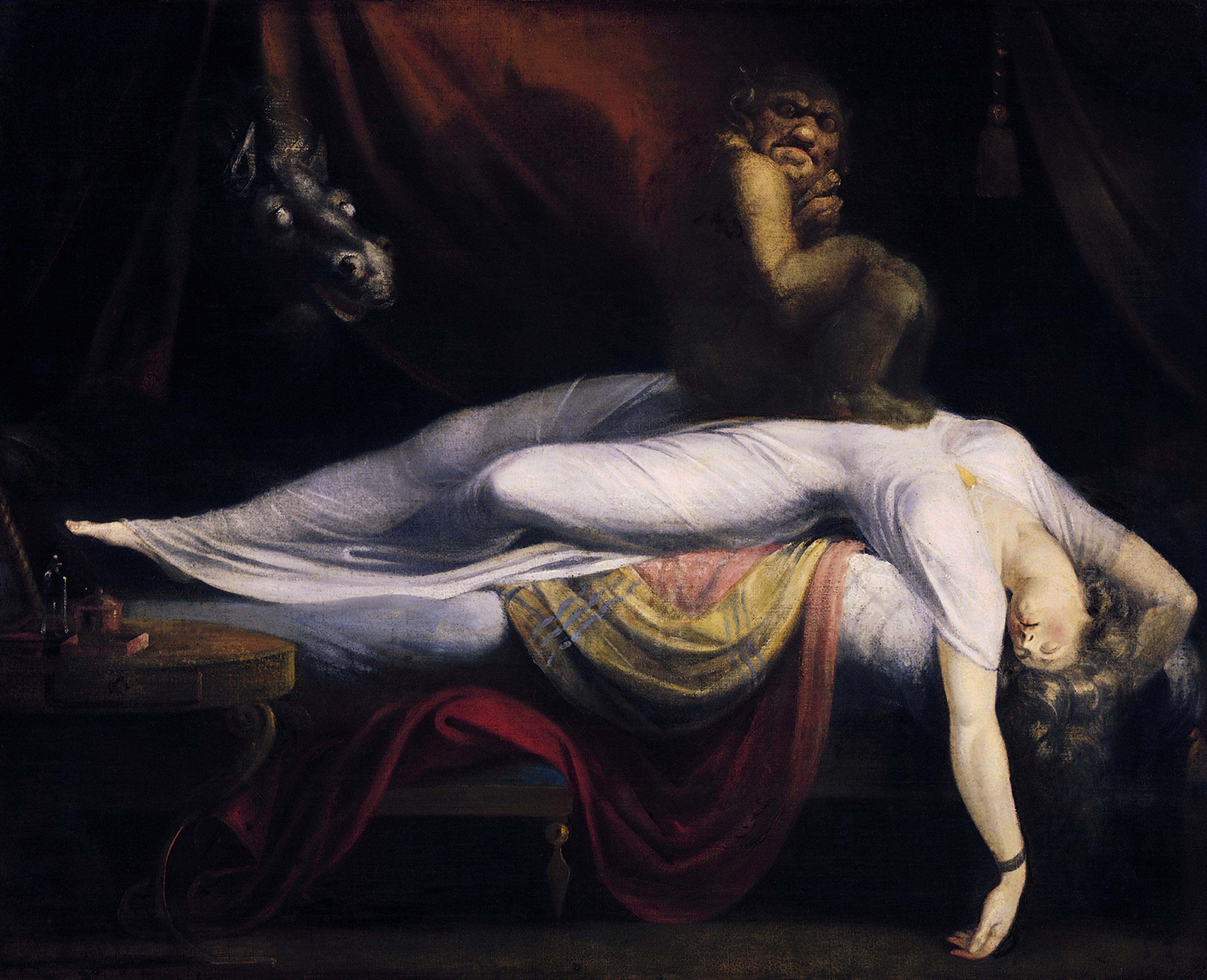

### n1.jpg

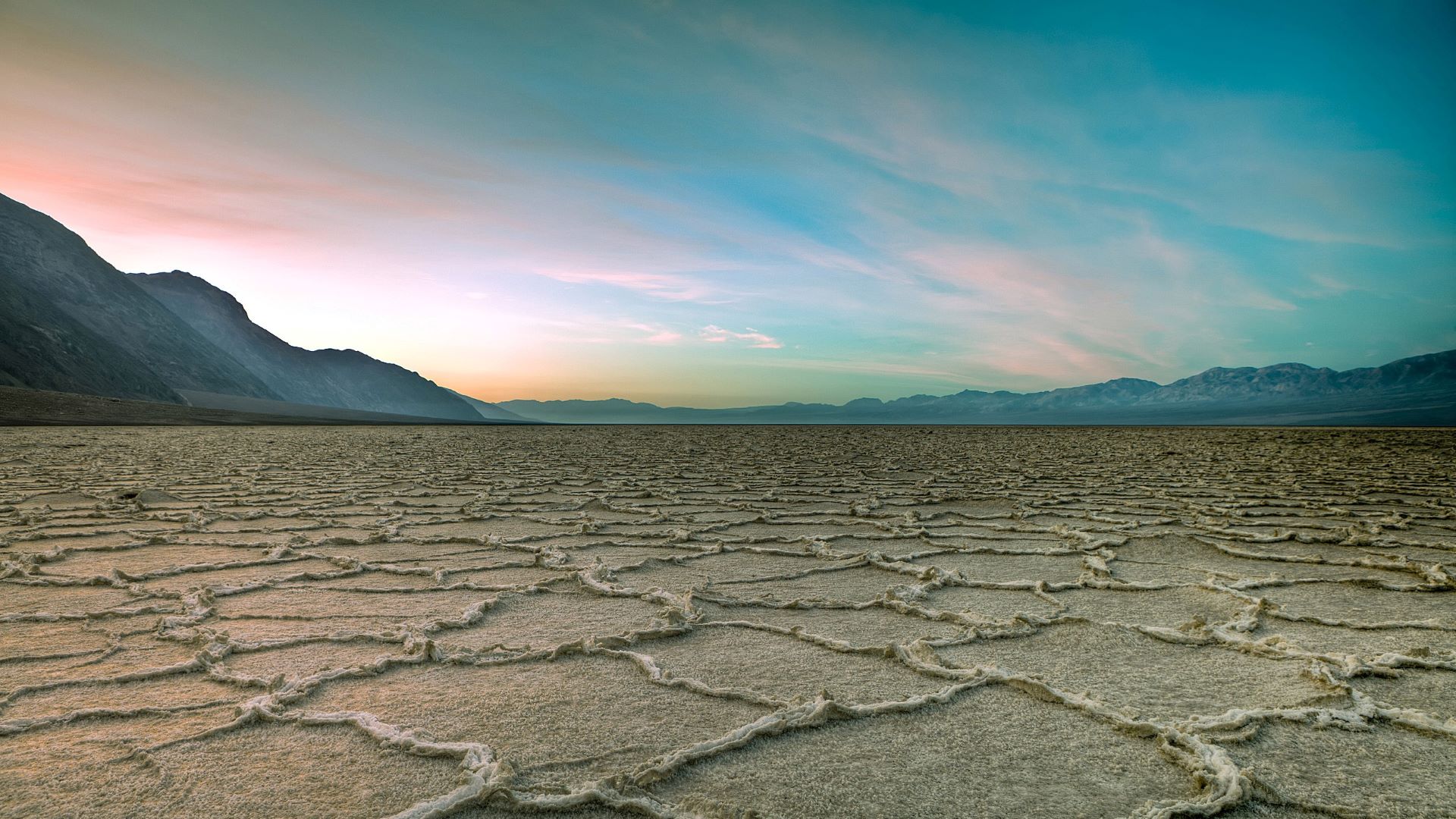

### Supplementary Material 2.docx

**Supplementary Material 2**

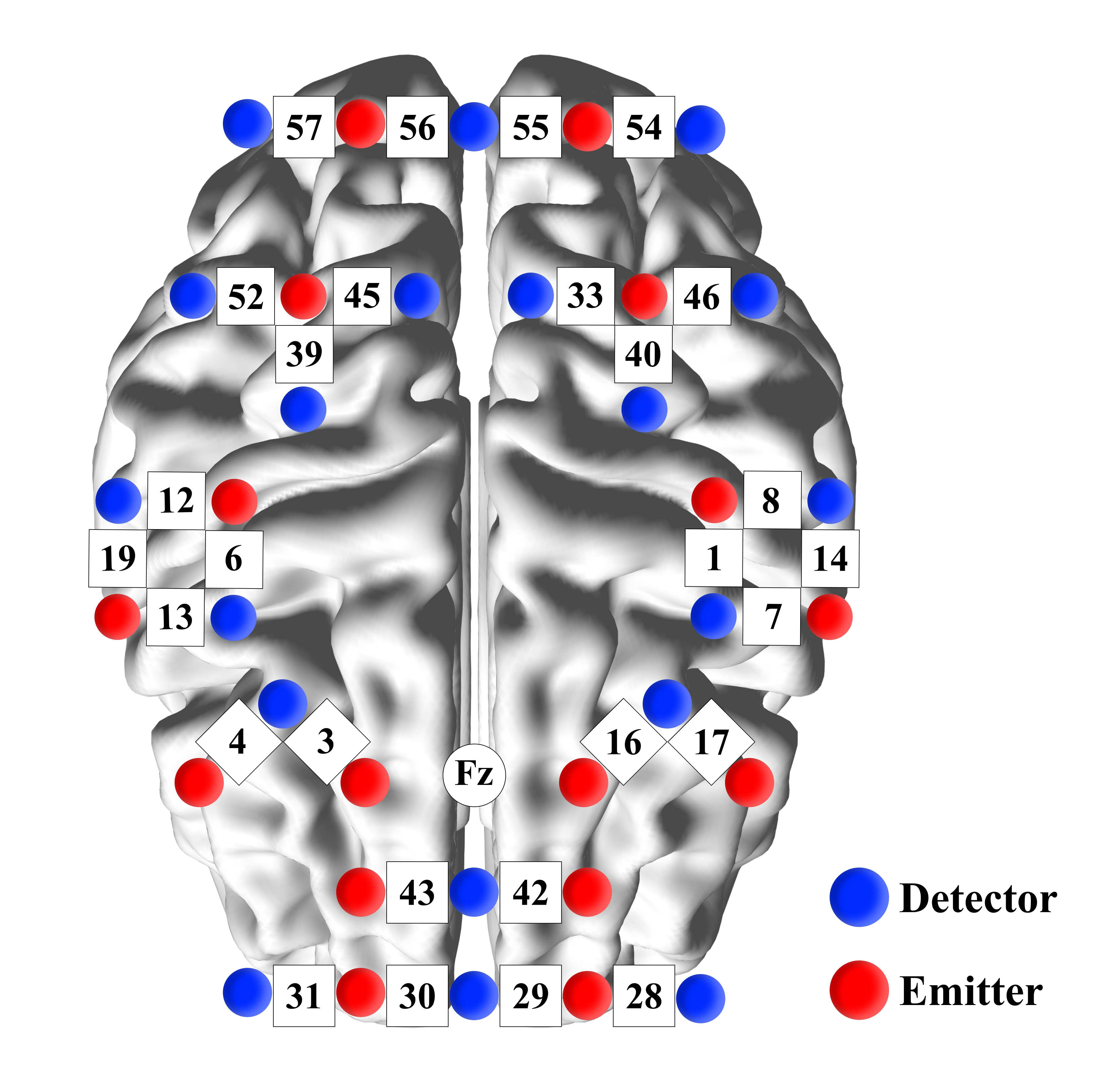

### 微信图片_20260122033314.jpg

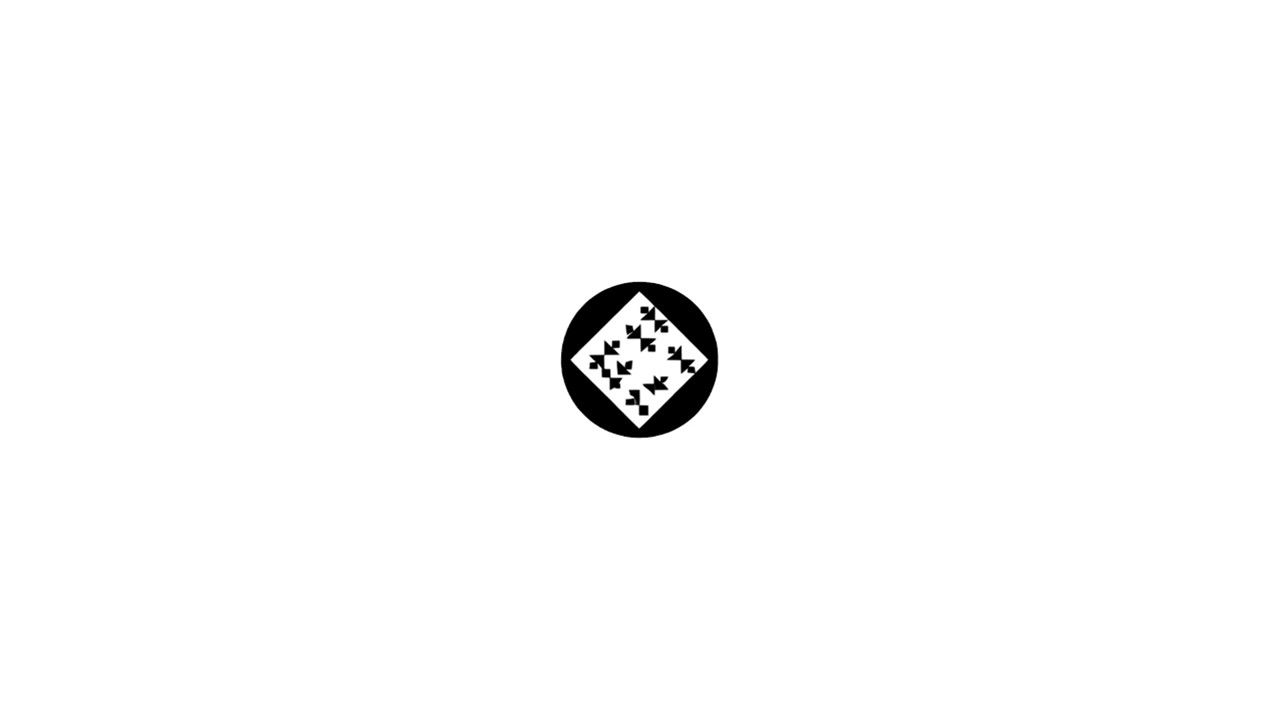

### 微信图片_20260122214853.jpg

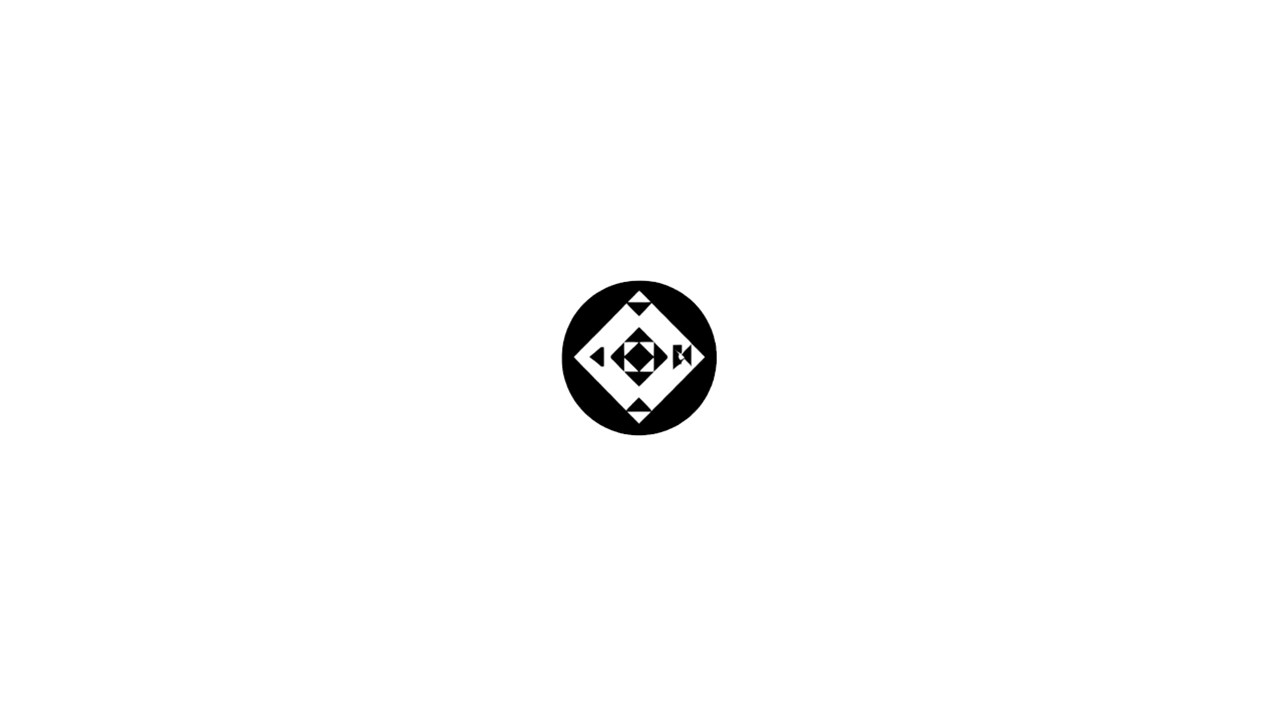

### 微信图片_20260122214903.jpg

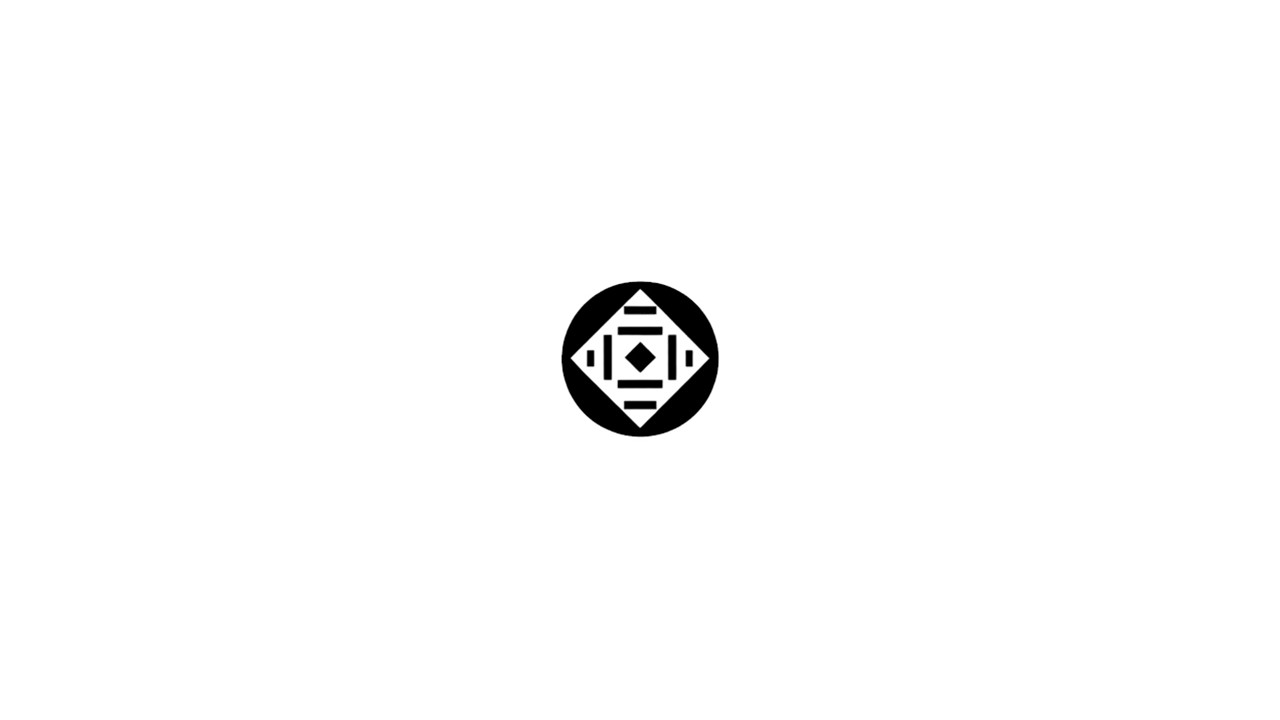

### 微信图片_20260122214907.jpg

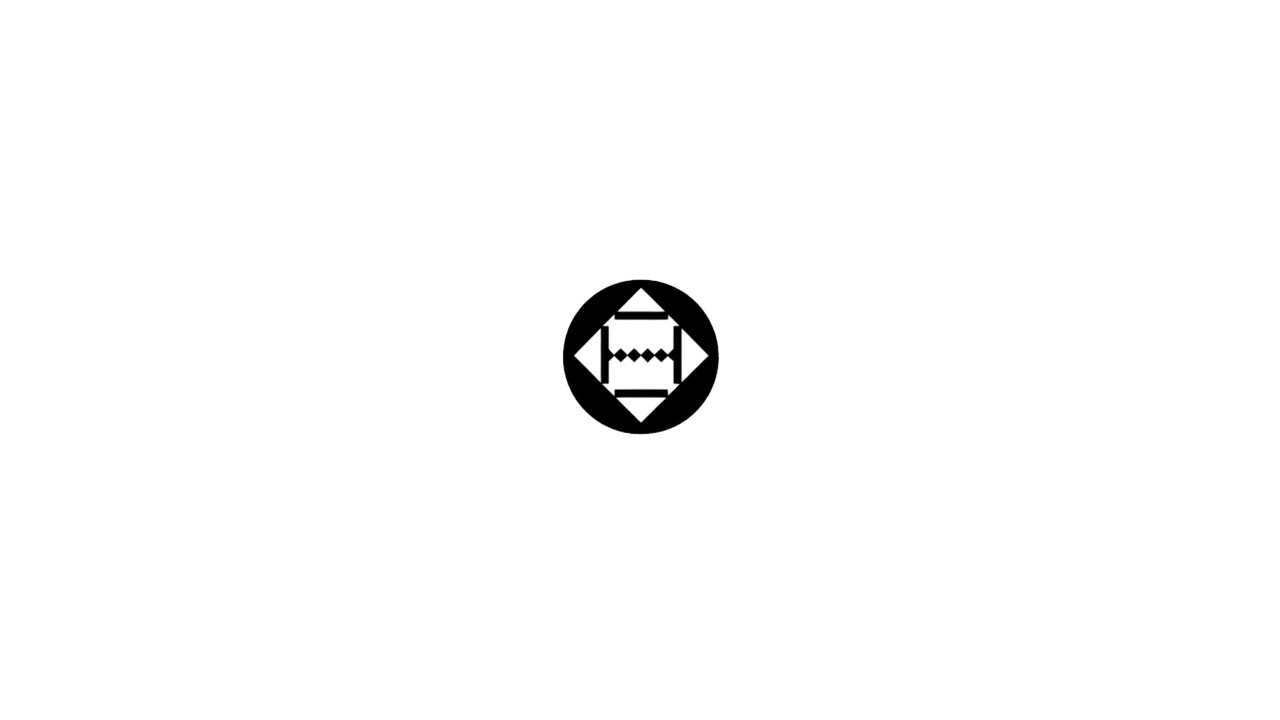

### 微信图片_20260122214912.jpg

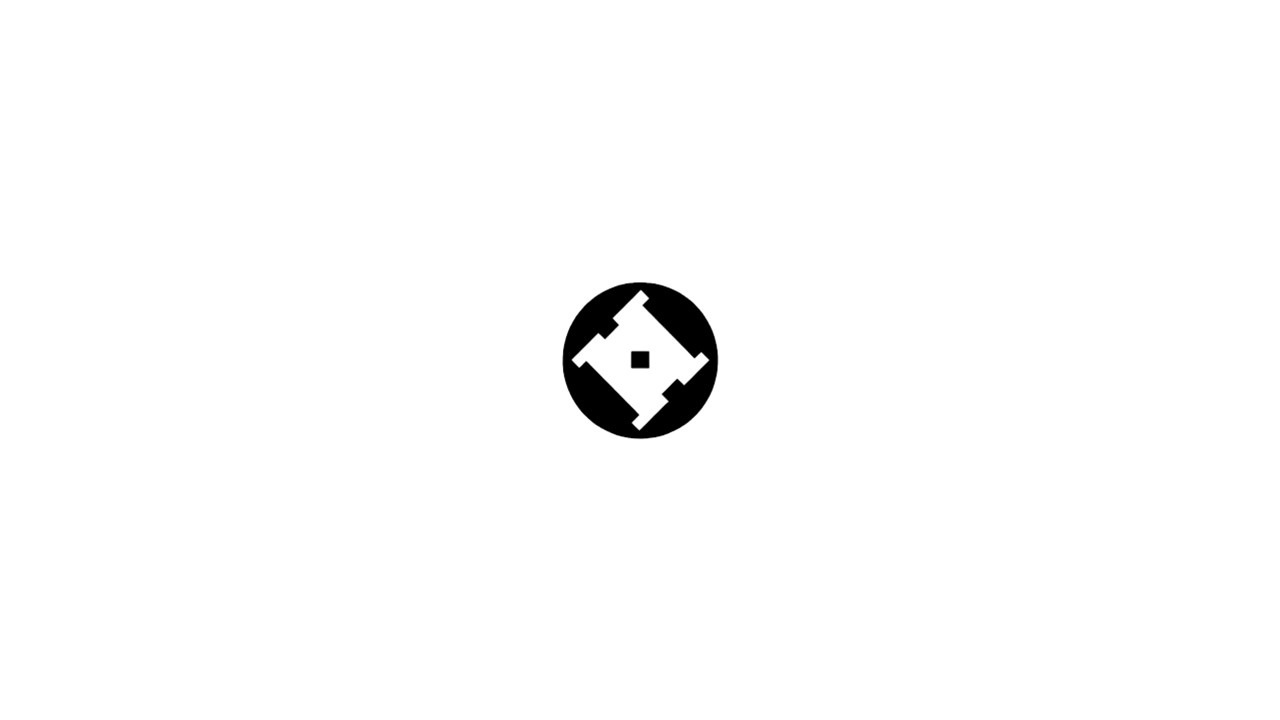

### 微信图片_20260122214916.jpg

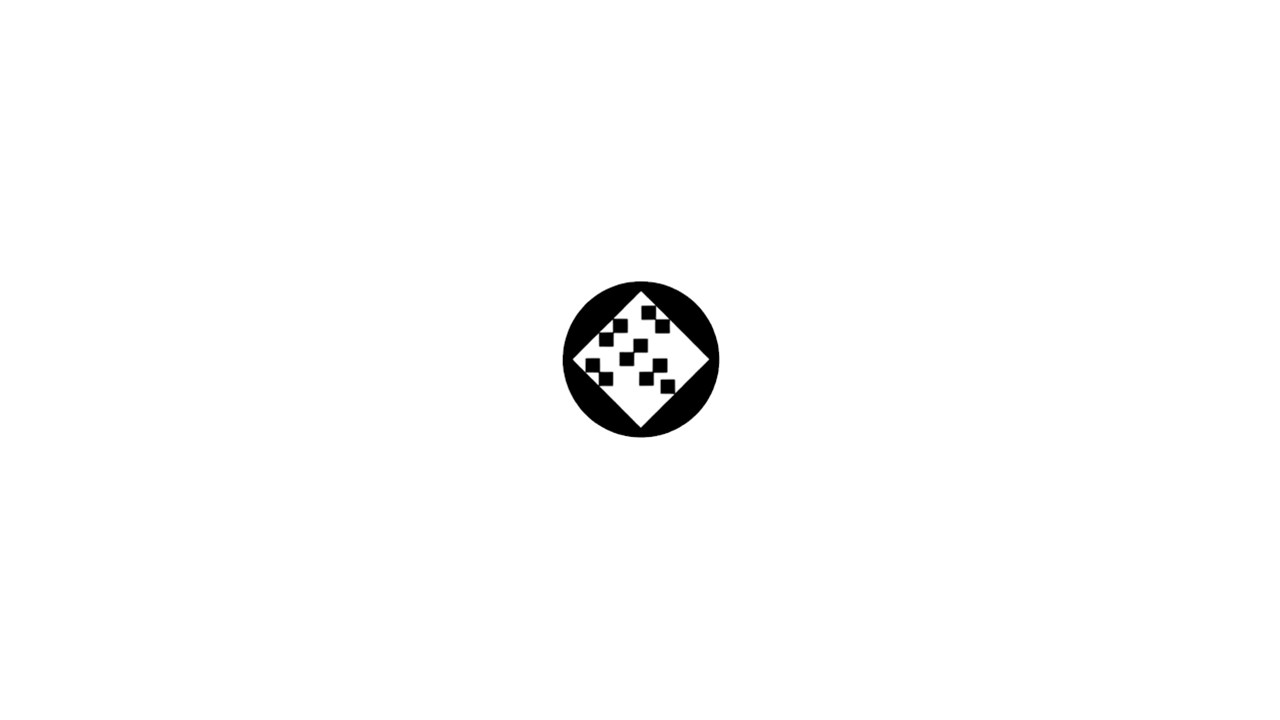

### 微信图片_20260122214920.jpg

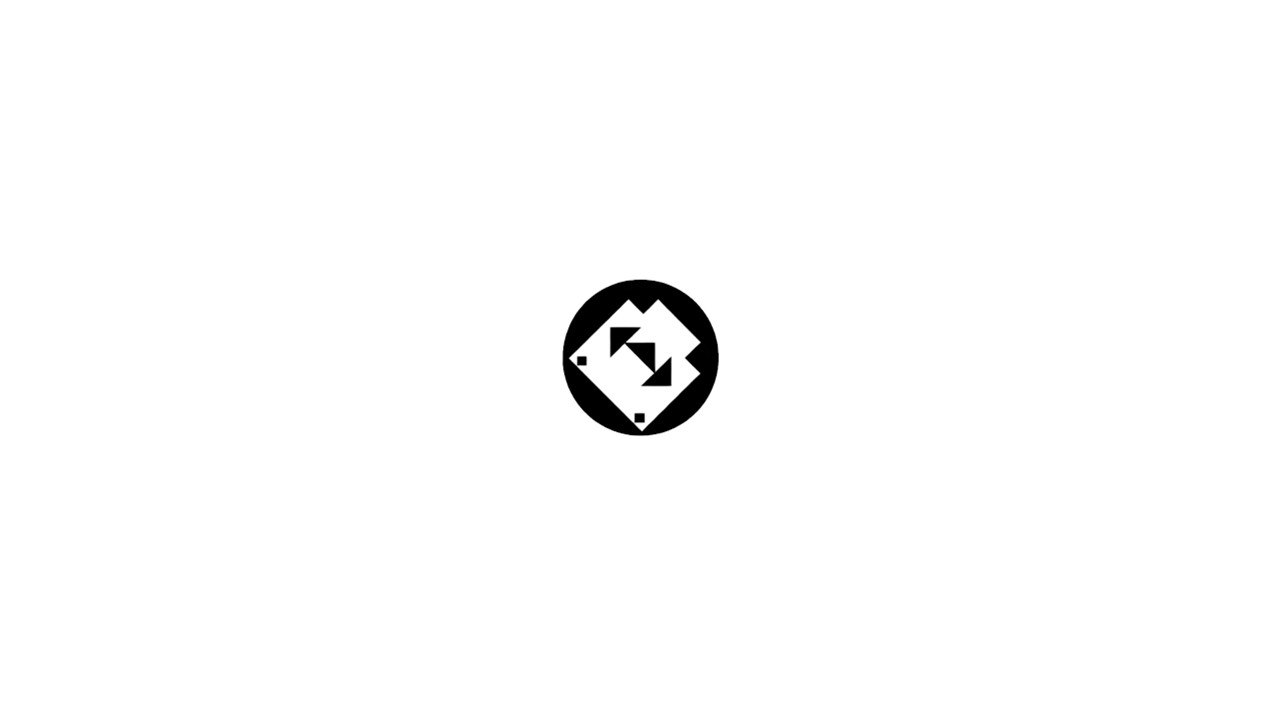

### 微信图片_20260122214923.jpg

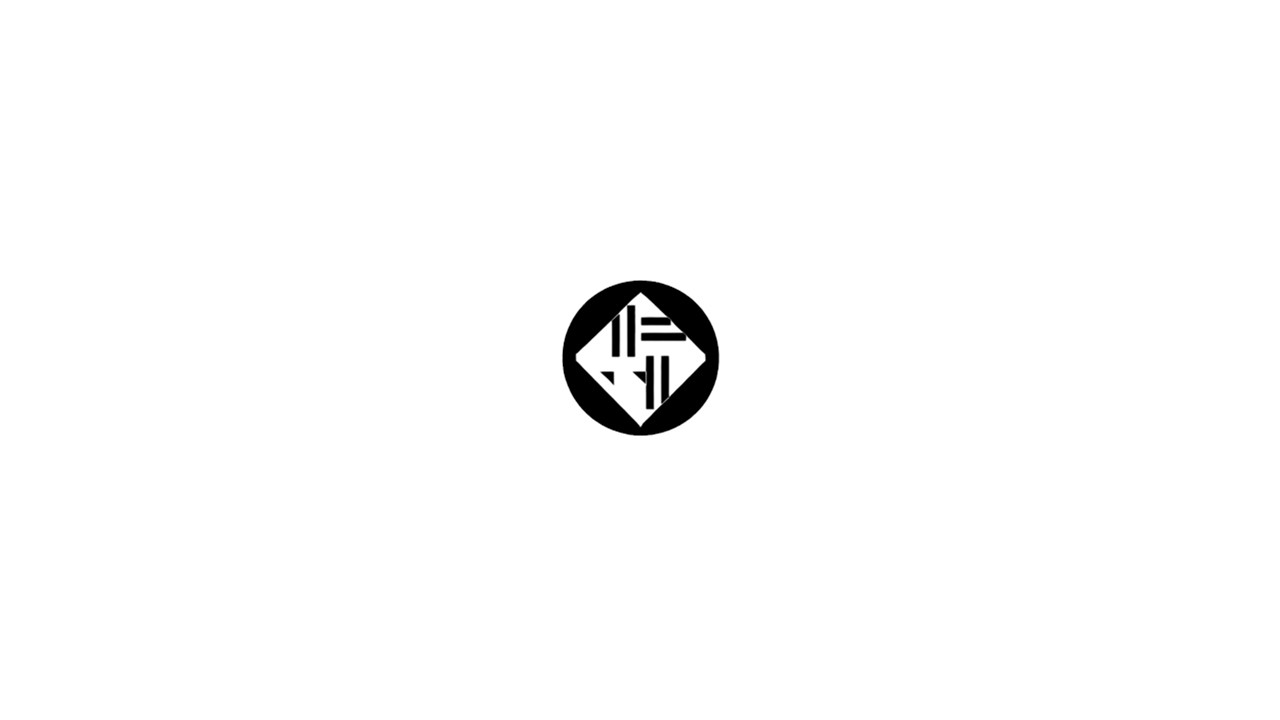
