## Supplementary Note S1 for "When Deeper Analysis Weakens Aesthetic Experience: Behavioral and Brain Network Evidence"

**Supplementary Note S1. Effect of the removed-rank asymmetry on sub-network entropy.**

Because K₂ was estimated separately per group (deep = 6, shallow = 5) and the baseline subspaces were disjoint, the residual subspace had 18 dimensions in the deep group and 19 in the shallow group. To determine whether this one-dimension asymmetry could by itself generate a group difference in topographic entropy, we ran a null simulation in which no true group difference existed and the two groups differed only in the number of dimensions removed (10 vs. 9). Twenty-eight-channel narrowband data were generated from a structured spatial covariance; the removed subspace was taken either as the leading eigenvectors of an independently generated structured baseline covariance (matching the real pipeline) or as a random orthonormal basis. Residual GED, forward-model recovery, and normalised Shannon entropy were computed exactly as in the main analysis, for sub-network sizes of 8, 12, and 16 channels (200 simulated participants per cell).

Under structured removal, the entropy difference attributable to the dimensionality asymmetry was +0.022, −0.005, and +0.000 for sub-network sizes of 8, 12, and 16 (|d| ≤ 0.19). Under random removal it was −0.014, −0.021, and −0.011 (|d| ≤ 0.24), i.e. opposite in sign to the observed effect. In every configuration the simulated bias was at least an order of magnitude smaller than the observed DMN difference (0.129, d = 0.84). The dimensionality asymmetry therefore cannot account for the reported effect.
