## Supplementary material for "When Deeper Analysis Weakens Aesthetic Experience: Behavioral and Brain Network Evidence": Table S1

**Table S1.** *Change in aesthetic rating relative to the passive-viewing baseline (Experiment 1).* Change = post − baseline on a 0–100 scale; *n* = 18 per group, df = 17. TOST = two one-sided tests against ±5 rating points; *B*_min_ = smallest symmetric bound at which equivalence would be established.

| **Image type** | **Dimension** | **Group** | ***M*** | ***SD*** | **95% CI** | ***t*** | ***p*** | ***d*** | **90% CI** | **TOST *p*** | ***B*_min_** |
| --- | --- | --- | --- | --- | --- | --- | --- | --- | --- | --- | --- |
| Visual Art | Perceptual | Shallow | 3.01 | 10.88 | [−2.40, 8.42] | 1.17 | .257 | 0.28 | [−1.45, 7.47] | .224 | 7.47 |
| Visual Art | Perceptual | Deep | 7.13 | 8.93 | [2.69, 11.57] | 3.39 | **.004** | 0.80 | [3.47, 10.79] | — | 10.79 |
| Visual Art | Semantic | Shallow | 3.65 | 10.56 | [−1.61, 8.90] | 1.46 | .161 | 0.35 | [−0.69, 7.98] | .297 | 7.98 |
| Visual Art | Semantic | Deep | 1.58 | 5.60 | [−1.20, 4.37] | 1.20 | .246 | 0.28 | [−0.71, 3.88] | **.010** | 3.88 |
| Visual Art | Affective | Shallow | 5.83 | 8.47 | [1.62, 10.05] | 2.92 | **.010** | 0.69 | [2.36, 9.31] | — | 9.31 |
| Visual Art | Affective | Deep | −0.86 | 8.09 | [−4.89, 3.16] | −0.45 | .657 | −0.11 | [−4.18, 2.46] | **.022** | 4.18 |
| Nat. Landscape | Perceptual | Shallow | −3.09 | 10.36 | [−8.24, 2.07] | −1.26 | .224 | −0.30 | [−7.33, 1.16] | .222 | 7.33 |
| Nat. Landscape | Perceptual | Deep | 4.66 | 7.75 | [0.80, 8.51] | 2.55 | **.021** | 0.60 | [1.48, 7.83] | — | 7.83 |
| Nat. Landscape | Semantic | Shallow | −4.14 | 11.51 | [−9.87, 1.58] | −1.53 | .145 | −0.36 | [−8.86, 0.58] | .378 | 8.86 |
| Nat. Landscape | Semantic | Deep | 4.28 | 7.43 | [0.59, 7.98] | 2.45 | **.026** | 0.58 | [1.24, 7.33] | — | 7.33 |
| Nat. Landscape | Affective | Shallow | −4.87 | 12.23 | [−10.96, 1.21] | −1.69 | .109 | −0.40 | [−9.89, 0.14] | .483 | 9.89 |
| Nat. Landscape | Affective | Deep | 5.00 | 6.59 | [1.72, 8.28] | 3.22 | **.005** | 0.76 | [2.29, 7.70] | — | 7.70 |

TOST reported only for cells that did not differ from zero.
